# Evolutionary constraints on mitochondrial bioenergetics are reshaped in cancer

**DOI:** 10.64898/2026.11.20.768050

**Authors:** Ji-Yong Sung, Jae-Ho Cheong

**Author notes:** Corresponding authors: Jae-Ho Cheong, M.D., Ph.D., Department of Surgery, Yonsei University College of Medicine, Seoul 03722, Republic of Korea.

## Abstract

Mitochondrial DNA encodes thirteen subunits of the oxidative phosphorylation system, many of which are subject to strong purifying selection in humans. In cancer this selection is relaxed, which is usually read as a weakening of the inherited filter. Comparing somatic variants from 8,886 tumours with population variation using a unified analytical framework and a dataset-specific mutational null, we show that the tumour filter has a different shape. ATP synthase, the least constrained complex in the germline, becomes the most constrained in tumours, an inversion that holds within the tumour data alone and is not explained by the measurement ceiling. Holding residue position fixed, the germline avoids substitutions that change side-chain charge at the proton-loading sites of complex I, as does the human phylogeny, and around the haems of complex III, whereas tumours no longer distinguish them. Damage at the redox centres of complexes III and IV is tolerated mainly at low variant allele fraction, including within single cohorts, and tumours retain constraint on the electron-tunnelling medium of complex IV. At a complex I proton bridge, density-functional calculations separate substitutions that retune the transfer potential from those that abolish it, and a gated-tunnelling model predicts that removing the charge of one tumour-mutated residue changes the rate far more than the kinetic isotope effect.

## Introduction

Mitochondrial DNA (mtDNA) encodes thirteen subunits of the oxidative phosphorylation system, and missense variation in them is removed from human populations by strong purifying selection ^1–3^. In tumours this selection is relaxed: somatic missense variants accumulate close to neutrality, truncating variants appear across the genes, and the complexes differ in the mutations they accumulate ^4–7^. The relaxation is usually read as a weakening of the same filter, in which somatic variation is a faded copy of inherited variation ^8,9^. That reading has not been tested, because the natural measure cannot test it. The ratio of somatic to germline dN/dS is bounded above by roughly the reciprocal of the germline estimate, so a gene near neutrality in the germline cannot show a large ratio, and one that tumours constrain more would look merely unrelaxed ^8^. Whether the two filters have the same shape must be asked with ranks and with comparisons made within one dataset.

The question also has a physical side. If the two filters differ in shape, the difference should be clearest at the steps where a single substitution acts directly on energy conversion, and these are steps of electron and proton transfer. Each complex couples chemistry to the membrane through electron and proton transfer steps whose rates are set by local structure: electron tunnelling between redox centres ^10,11^, oxygen reduction at the haem a3–CuB centre of complex IV ^12^ and proton transfer along hydrogen-bonded chains ^13–15^. At these steps quantum mechanics enters directly, because electrons and protons cross barriers by tunnelling at rates set by the donor–acceptor distance and its modulation by protein motion ^10,16^. A substitution can retune such a step, by changing a charge near a group that stays in place, or break it, by removing the group that carries the particle. A distant charge should shift the reactant and product energies but leave the shape of barrier, and with it the contribution of tunnelling, largely unchanged; a change in barrier geometry should alter both the rate and its sensitivity to isotopic mass. Earlier analyses of single complexes compared structural classes with the rest of their protein, which confounds chemistry with position, and in complex III could not separate electron-transfer from electrostatic effects ^17^. Which of the two tumours release (i.e., tolerate although the germline avoids it) has remained open. Dosage adds a third dimension. A damaging variant impairs respiration only once its fraction among the hundreds to thousands of mtDNA copies exceeds a threshold ^18,19^. If wild-type genomes mask the damage, tumour tolerance should fade as the mutant fraction rises; whether it does, and alike across the complexes, is unknown.

Here we analyse all four mtDNA-encoded complexes under one pipeline, with one mutational null per dataset, one set of resamples and proton-wire graphs built from four human cryo-EM structures with one set of parameters ^20,21^. The tumour filter is not a weakened copy of the germline filter. ATP synthase exchanges the extremes, least constrained in the germline and most constrained in tumours, also within the tumour data alone. Damage to complexes III and IV is tolerated mainly while wild-type genomes predominate. Holding residue position fixed, the germline avoids substitutions that change side-chain charge at the proton-loading sites of complex I ^15^, as does the human phylogeny, and around the haems of complex III, whereas tumours no longer distinguish them. Along computed electron-tunnelling pathways ^22^ the germline carries no charge change, fewer than expected even from matched residues in complex III, a protection that tumours lose; tumours in complex IV keep the tunnelling medium. Density-functional calculations show that a released charge shifts the wells of a proton-transfer potential while barely changing its barrier, whereas a substitution at the accepting group abolishes the step. In a gated-tunnelling model with a stiff protein scaffold, most of this proton transfer occurs by tunnelling, and the model predicts the isotope effect that would test it ^16,23^.

## Results

### The tumour filter is not a weaker copy

We assembled 16,152 somatic single-nucleotide variants (11,435 in coding sequence) from 8,886 tumours in TCGA, PCAWG and MSK-IMPACT, and population variation from HelixMTdb ^6,24,25^, each scored against a strand-aware, trinucleotide-resolved null estimated from that dataset alone (**Supplementary Fig. 1**); synonymous variants fell close to expectation in the classes analysed below (**Supplementary Fig. 2**). In the germline, missense dN/dS was low for complexes I (0.35), III (0.59) and IV (0.39) and close to neutral for complex V (0.92, 95% CI 0.81–1.04); in tumours it was 1.16, 1.25, 1.01 and 0.85 (0.68–1.10), respectively (**Fig. 1a and Supplementary Table 1**). Tumour-to-germline fold changes were 3.35 (3.08–3.63), 2.12 (1.76–2.52) and 2.59 (2.27–2.92) for complexes I, III and IV, against 0.93 (0.71–1.22) for complex V (**Fig. 1b**), which relaxed 0.35 times as much as the other three (0.27–0.47; bootstrap P ≤ 0.001); we refer to this as the exemption of complex V.

**Fig. 1.**
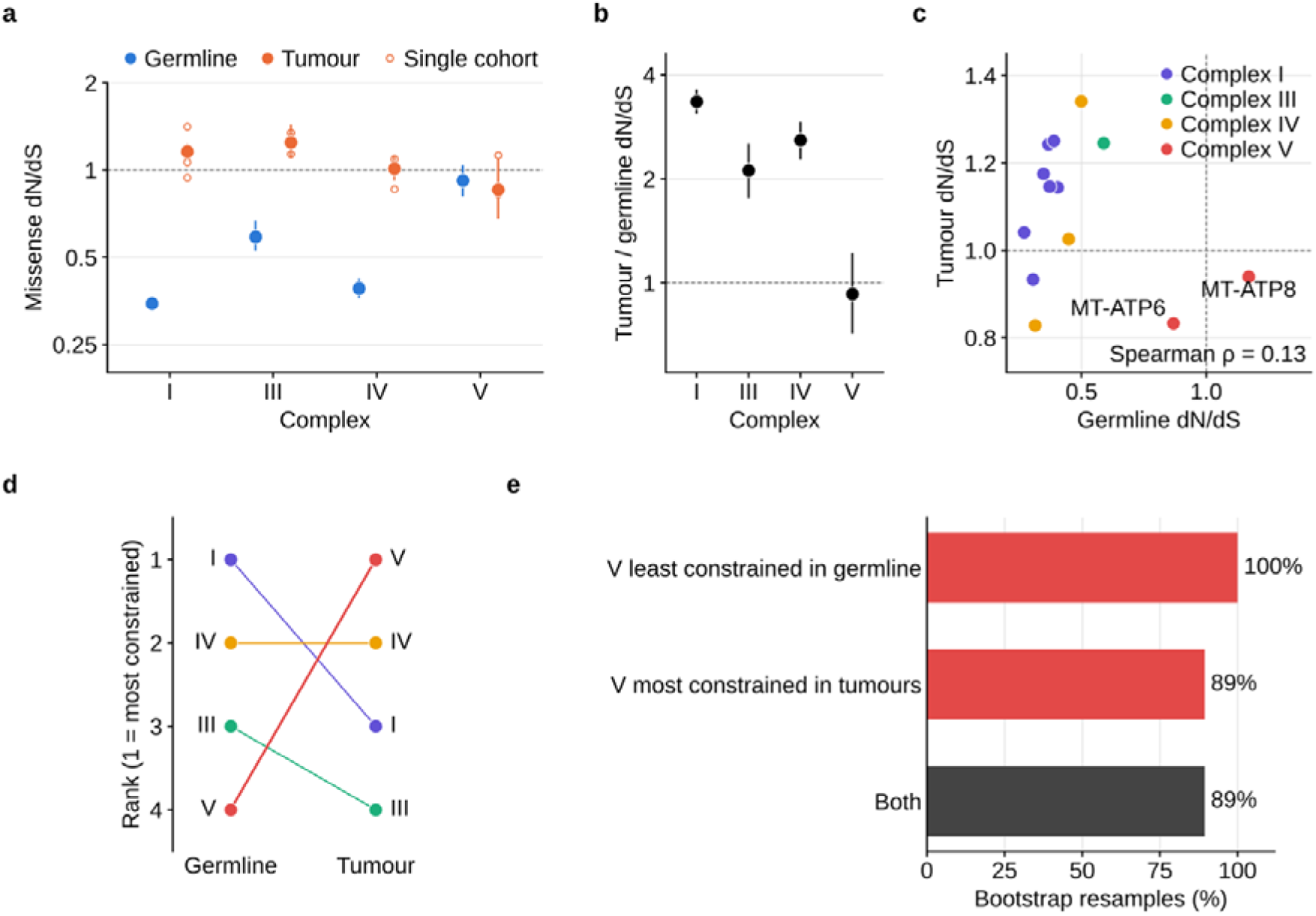
The germline and tumour filters are differently shaped, and complex V inverts its rank. a, Missense dN/dS of the mtDNA-encoded subunits of complexes I (MT-ND1–6, MT-ND4L), III (MT-CYB), IV (MT-CO1–3) and V (MT-ATP6, MT-ATP8). Germline estimates come from 12,224 HelixMTdb variant sites and tumour estimates from 16,152 somatic single-nucleotide variants in 8,886 tumours (TCGA, PCAWG and MSK-IMPACT), pooled (filled) or from each cohort alone (open). Each dataset is scored against its own strand-aware null of 192 trinucleotide substitution classes. b, Tumour-to-germline fold change in dN/dS: complex I 3.35 (95% CI 3.08–3.63), III 2.12 (1.76–2.52), IV 2.59 (2.27– 2.92) and V 0.93 (0.71–1.22). c, Germline against tumour dN/dS for each of the thirteen genes, coloured by complex. MT-ATP6 ranks twelfth of thirteen in the germline and second in tumours, and MT-ATP8 thirteenth and fourth. The rank correlation (Spearman ρ = 0.13, −0.25 to 0.53) cannot be resolved with thirteen genes. d, Rank of each complex under the two filters, from most (1) to least (4) constrained. e, Fraction of bootstrap resamples in which complex V was the least constrained complex in the germline, the most constrained in tumours, and both. Error bars show 95% percentile intervals from 1,000 bootstrap resamples. Tumours were resampled by patient and population variants by site, with one fixed set of resamples per dataset. Dashed lines mark dN/dS = 1 and no relaxation. The y axes in a and b are on a log scale.

Because fold changes are bounded by the germline estimate, we compared the filters on ranks. Ranked from most to least constrained, MT-ATP6 moves from twelfth of thirteen genes in the germline to second in tumours, and MT-ATP8 from thirteenth to fourth (**Fig. 1c and Extended Data Fig. 1a**); the gene-level rank correlation (Spearman ρ = 0.13, −0.25 to 0.53) cannot be resolved with thirteen genes, and the evidence is the inversion, not the absence of a correlation. At the level of complexes, complex V is the least constrained complex in the germline in 100% of bootstrap resamples, the most constrained in tumours in 89%, and both at once in 89% (**Fig. 1d, e**). The rank reflects relative constraint: tumour dN/dS of complex V itself (0.85, 0.68–1.10) does not differ significantly from neutrality, which the within-tumour comparisons below address directly. With gnomAD v3.1 ^3^ (56,434 genomes) as the reference, the relative relaxation of complex V is again 0.34 (0.26–0.46) and the inversion holds in 89% of resamples (**Extended Data Fig. 2**).

### The exemption survives the ceiling

Weakly constrained regions of the other complexes did not appear unrelaxed: the nine most weakly constrained positional bins of complexes I, III and IV, selected on one half of the population variants and evaluated on the other, rose from 0.54 (0.48–0.61) in the germline to 1.29 (1.17–1.41) in tumours, whereas complex V started at 0.92 and did not move (**Fig. 2a**; within-tumour ratio 0.67, 0.51–0.87; P = 0.005). Statistics computed within the tumour data contain no germline term. There, missense dN/dS of complex V was 0.75 times that of complexes I, III and IV (0.59–0.97; one-sided P = 0.017) and 0.71 times that of MT-CO2 and MT-CO3 (0.54–0.93; P = 0.012; **Fig. 2b**).

**Fig. 2.**
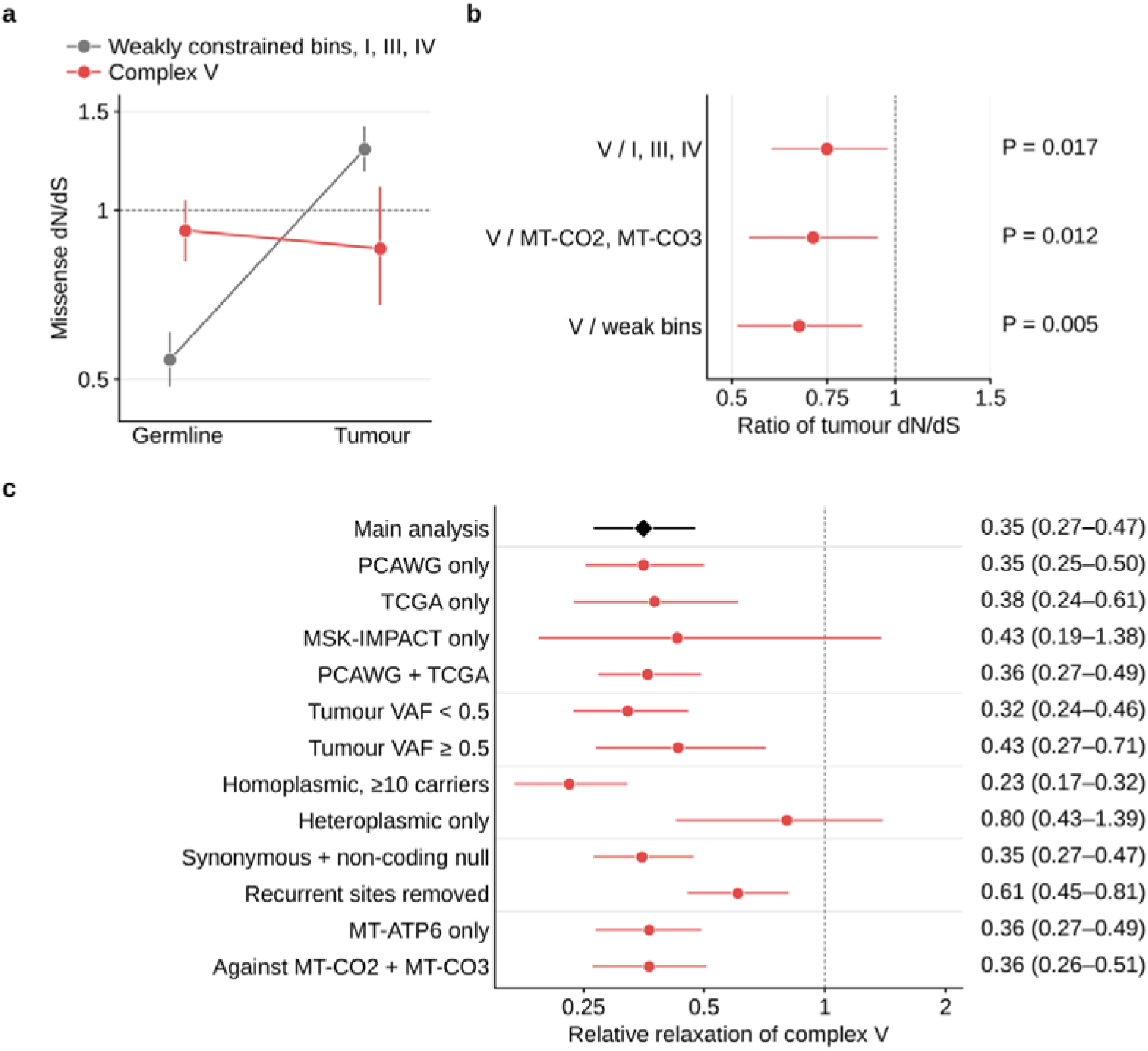
The exemption of complex V survives removal of the measurement ceiling and stays below 1 under every analysis choice varied. a, Matched-constraint control. The nine most weakly constrained positional bins of complexes I, III and IV (grey) were selected on one random half of population variants, and their germline dN/dS was estimated on the other half. These bins start at 0.54 (95% CI 0.48–0.61) and relax 2.37-fold past neutrality in tumours (1.29, 1.17–1.41). Complex V (red) starts at 0.92 and does not move (0.85). b, Comparisons within the tumour data alone, which contain no germline term and are therefore free of the ceiling. Shown is the missense dN/dS of complex V divided by that of complexes I, III and IV (0.75, 0.59–0.97), of the neighbouring genes MT-CO2 and MT-CO3 (0.71, 0.54– 0.93) and of the weakly constrained bins in a (0.67, 0.51–0.87). c, Relative relaxation of complex V, its tumour-to-germline fold change divided by the geometric mean of those of complexes I, III and IV, in the main analysis (diamond) and with one analysis choice changed per row (circles). The rows vary the tumour cohort, the tumour variant allele fraction (VAF), the germline reference, the null model and the gene sets compared. Error bars show 95% percentile intervals from the shared set of 1,000 bootstrap resamples, with tumours resampled by patient and population variants by site. P values are one-sided bootstrap probabilities that the ratio is at least 1. Dashed lines mark a ratio of 1. The y axis in a and the x axes in b and c are on a log scale.

The relative relaxation of complex V stayed below 1 in its point estimate under every analysis choice we varied (**Fig. 2c**), including each tumour cohort, both heteroplasmy strata, an alternative null, removal of recurrent sites (0.61, 0.45–0.81), MT-ATP6 alone (0.36, 0.27– 0.49) and a comparison with the neighbouring MT-CO2 and MT-CO3 (0.36, 0.26–0.51), and in each of eight cancer types (0.25–0.98; **Extended Data Fig. 1b**), although in five of them (breast, endometrial, hepatobiliary, ovarian and renal cell) the interval included 1. Among the other analysis choices, only MSK-IMPACT alone (32 complex V substitutions; 0.43, 0.19– 1.38) and a germline reference of heteroplasmic variants, which are too recent to have been filtered (0.80, 0.43–1.39), did not reach significance.

Truncating substitutions were relaxed in every complex, complex V included (**Extended Data Fig. 3**), so the exemption concerns missense variation. Tumour constraint in subunit a concentrated around Arg159 (0.36, 0.18–0.87), but not significantly after correction (family-wise P = 0.076; **Extended Data Fig. 4**), and synonymous variants were in excess in this shell in tumours (**Supplementary Fig. 2**), which would inflate its apparent constraint.

### Damage at redox centres is tolerated at low dosage

Damage at the redox centres, by contrast, depends on heteroplasmy. Charge-changing substitutions at the 94 MT-CYB residues within 8 Å of the b haems of complex III (PDB 9HZL; quinone sites and Rieske interface removed), which can shift haem redox potentials and so the driving force of electron transfer ^10,11^, were nearly absent from the germline (dN/dS 0.23, 0.09–0.39) and in excess in tumours (1.66, 1.34–2.05), in each cohort (**Extended Data Fig. 5a**). The excess lay below a variant allele fraction (VAF) of 0.5 (2.08, 1.63–2.71) and disappeared above it (1.04, 0.69–1.50; **Fig. 3a**), and membership of the class fell with VAF (cohort-adjusted logistic slope −1.15, −1.79 to −0.51; P = 4 × 10) whereas charge-preserving substitutions at the same residues did not (**Extended Data Fig. 5d**). The pattern held at 6 Å, in PDB 9CG3 and with our earlier residue set ^17^ (**Extended Data Fig. 5c** and **Supplementary Fig. 3**), and substitutions predicted to change a Q-cycle electron-transfer rate at least twofold ^22,26^ were enriched below a VAF of 0.5 (1.70) and not above it (1.08; **Supplementary Fig. 4**). Because VAF distributions differ between cohorts (median 0.04 in PCAWG against 0.51 in TCGA and MSK-IMPACT), we also divided the calls of each cohort into thirds of VAF: the excess fell from 2.00 (1.40–2.80) in the lowest third to 1.08 (0.71– 1.50) in the highest (ratio 0.54, 0.32–0.86; one-sided P = 0.004), and membership of the class fell with VAF within PCAWG (slope −2.72; P = 0.006) and within TCGA (−1.25; P = 0.03), without a cohort-by-VAF interaction (P = 0.08; **Supplementary Fig. 11a, b** and **Supplementary Table 1**). In TCGA alone the two strata were not resolved (**Supplementary Fig. 5**).

**Fig. 3.**
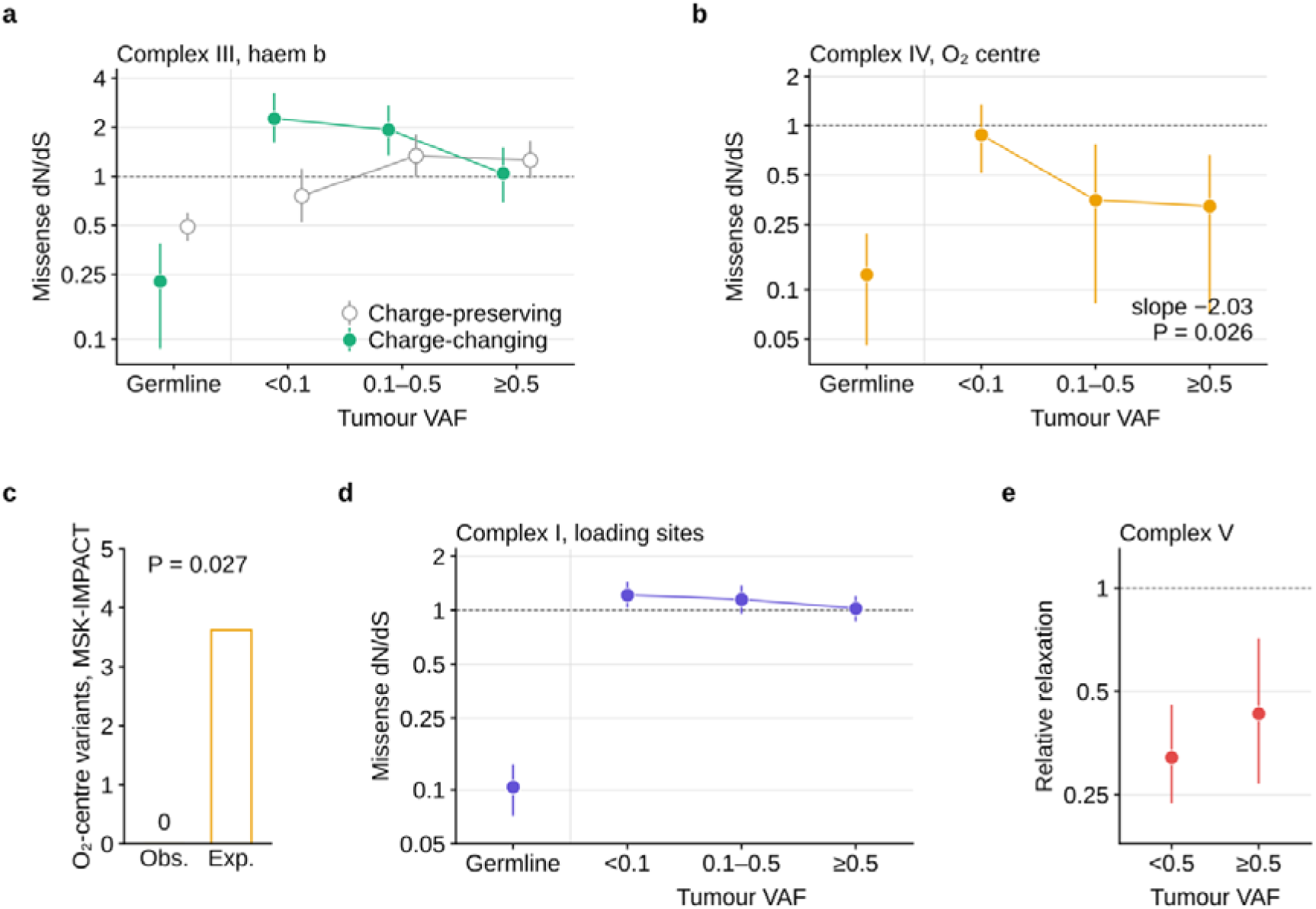
Damage to complexes III and IV, and more weakly complex I, is tolerated mainly while wild-type mtDNA predominates. a, Complex III. Missense dN/dS of MT-CYB substitutions at the 94 residues within 8 Å of the two b haems (PDB 9HZL), excluding the quinone sites and the Rieske interface. Substitutions that change formal side-chain charge (filled) are shown with those at the same residues that preserve it (open), in the germline and in tumours by variant allele fraction (VAF). Among MT-CYB missense variants in tumours, membership of the charge-changing class falls with VAF (cohort-adjusted logistic slope −1.15, 95% CI −1.79 to −0.51; P = 4 × 10□□), whereas membership of the charge-preserving class does not (+0.39; P = 0.10). b, Complex IV. Missense dN/dS at the 24 MT-CO1 residues within 8 Å of the haem a3 iron or CuB (PDB 9I7U), in the germline and in tumours by VAF, with the cohort-adjusted logistic slope. c, Observed and expected numbers of oxygen- reducing-centre variants among the 63 MT-CO1 missense mutations of MSK-IMPACT, whose median VAF is 0.50. The expected number is derived from the spectrum-weighted fraction of MT-CO1 missense opportunity at the centre; the P value is a one-sided Poisson probability. d, Complex I. Missense dN/dS of charge-changing substitutions at proton-loading sites (PDB 9I4I), in the germline and in tumours by VAF. e, Relative relaxation of complex V below and above a VAF of 0.5. dN/dS is taken against the synonymous changes of the same gene, with each dataset’s own null. In tumour strata, rates come from the whole tumour spectrum and observations from the stratum. Error bars show 95% percentile intervals from the shared set of 1,000 bootstrap resamples. Dashed lines mark dN/dS = 1 or a ratio of 1. The y axes are on a log scale except in c.

The oxygen-reducing centre of complex IV follows the same rule. Missense variants at the 24 MT-CO1 residues within 8 Å of the haem a3 iron or CuB were strongly depleted in the germline (0.12, 0.05–0.22; **Supplementary Fig. 6**) and less so in tumours (0.59, 0.38–0.82; **Extended Data Fig. 6**), where depletion deepened with VAF, from 0.88 below 0.1 to 0.32 above 0.5 (cohort-adjusted slope −2.03, −3.83 to −0.24; P = 0.026; **Fig. 3b**). In thirds of VAF within each cohort the depletion deepened from 0.93 to 0.17 (ratio 0.18, 0.04–0.49; P = 0.002), although MSK-IMPACT contributes no variant at the centre. Synonymous variants were, if anything, in excess there (**Supplementary Fig. 2**), and the bovine structure gave the same picture (**Extended Data Fig. 6a**). Of the 63 MT-CO1 missense mutations of MSK-IMPACT, a panel cohort whose variants have a median VAF of 0.50, none fell at the centre, against 3.6 expected (P = 0.027; **Fig. 3c**). In the germline, variants at the centre were concentrated among heteroplasmic-only sites (odds ratio 8.4; P = 0.027; **Supplementary Fig. 5a**), and in TCGA the depletion above a VAF of 0.5 persisted after correction for purity (**Supplementary Fig. 5b**).

Charge-changing substitutions at the proton-loading sites of complex I, defined below, followed the same direction more weakly (1.18, 1.05–1.33 below a VAF of 0.5; 1.03, 0.87– 1.20 above; **Fig. 3d**) and were concentrated among heteroplasmic-only germline sites (odds ratio 3.2; P = 0.001; **Supplementary Fig. 5a**). In thirds of VAF within each cohort the class fell from 1.26 to 0.89 (ratio 0.70, 0.55–0.88; P = 0.004), in PCAWG and MSK-IMPACT but not in TCGA (1.69, 0.98–3.04). This is the pattern expected if wild-type genomes buffer a cost that is enforced again once they no longer predominate ^18,19^. The exemption of complex V held in both strata (0.32, 0.24–0.46 and 0.43, 0.27–0.71; **Fig. 3e**).

### The germline avoids charge changes that tumours no longer distinguish

We built hydrogen-bonded proton-wire graphs for every mtDNA-encoded subunit from four human cryo-EM structures with one set of parameters (complex I, PDB 9I4I; complex III, 9HZL; complex IV, 9I7U; ATP synthase, 8H9F) ^20,21^, generating cavity water identically in each (Methods). They give 229 proton-loading sites and 122 cut vertices in complex I, 47 and 18 in complex III, 145 and 66 in complex IV, and 14 and 23 in ATP synthase (**Fig. 4a** and **Supplementary Table 2**). A substitution at a loading site can change the charge of a group that stays in place; one at a cut vertex can sever the pathway ^27,28^. Candidate substitutions in subunit a of ATP synthase on the two axes are listed in **Supplementary Table 3**.

**Fig. 4.**
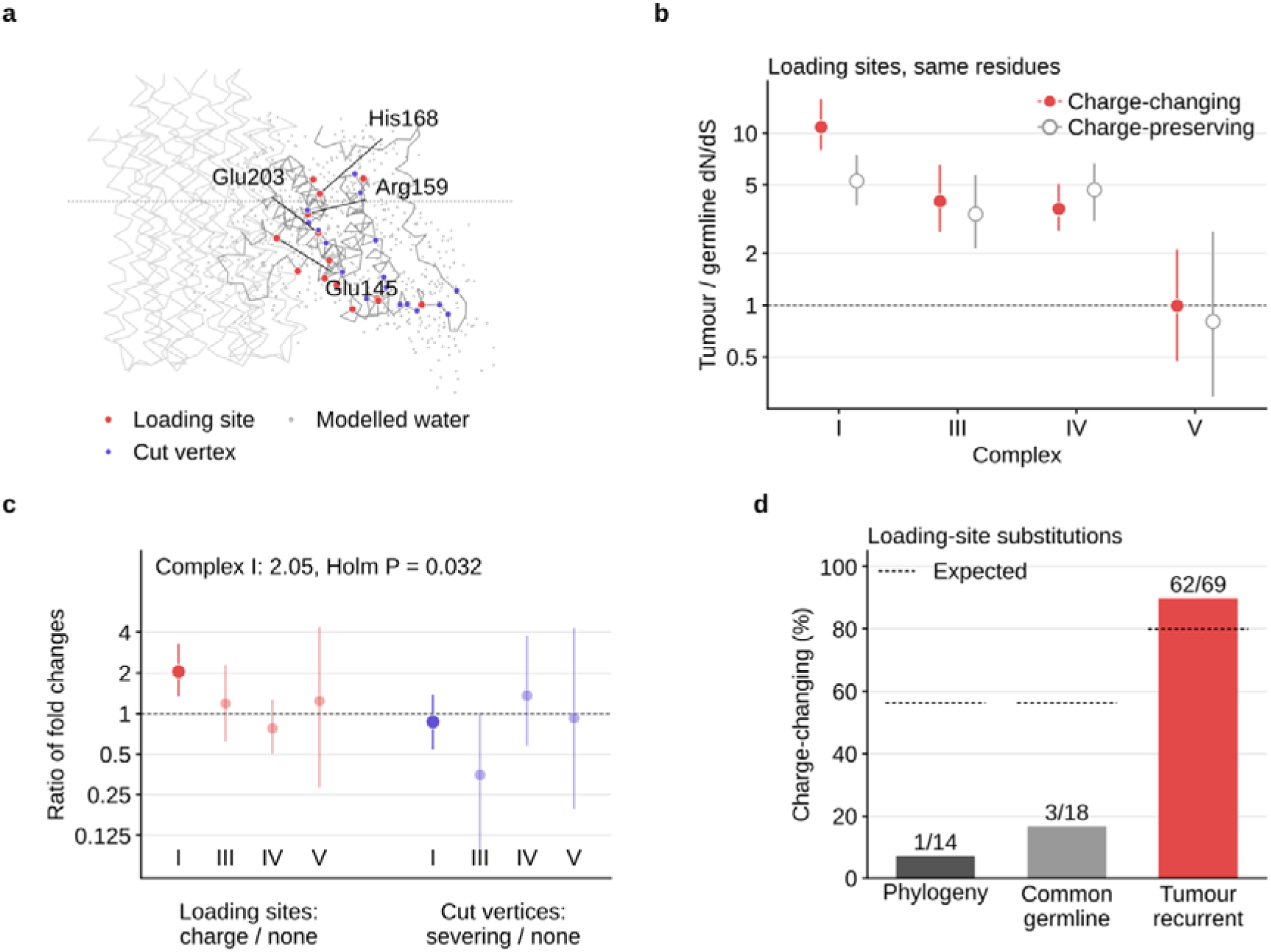
At the proton-loading sites of complex I, the germline avoids charge changes that tumours no longer distinguish. a, Proton-wire graph of subunit a of ATP synthase (PDB 8H9F) as an example, projected on the membrane normal (dotted line, membrane centre). Loading sites are shown in red, cut vertices in purple and modelled cavity water in grey, with the half-channel residues and Arg159 labelled. Graphs were built for the mtDNA-encoded subunits of all four complexes from PDB 9I4I (complex I), 9HZL (complex III), 9I7U (complex IV) and 8H9F (ATP synthase), with identical parameters and cavity water generated identically. b, Tumour-to-germline fold change in dN/dS at loading sites, for substitutions that change formal side-chain charge (filled) and for substitutions at the same residues that preserve it (open). c, Ratio of the two fold changes at loading sites (charge-changing over charge-preserving) and at cut vertices (wire-severing over non-severing), by complex; complex I is drawn solid. At complex I loading sites the ratio is 2.05 (95% CI 1.34–3.27; one-sided P = 0.002), the only one of the eight contrasts to survive Holm correction (adjusted P = 0.032). d, Percentage of complex I loading-site substitutions that change charge among those defining branches of the human mtDNA phylogeny (PhyloTree build 17), those carried by more than ten HelixMTdb individuals and those recurrent in four or more tumours. Dashed lines mark the percentage expected from the germline (first two bars) or tumour (third bar) spectrum over the same residues; binomial P = 2 × 10□□, 7 × 10□□ and 0.02. Error bars show 95% percentile intervals from the shared set of 1,000 bootstrap resamples. Dashed lines in b and c mark a ratio of 1. The y axes in b and c are on a log scale.

Comparing a wire class with the rest of its complex confounds chemistry with position: charge-changing substitutions at loading sites appear relaxed 3.46-fold in complex I, 2.23-fold in complex III and 1.62-fold in complex IV, but this ordering disappears once background positions are matched on germline constraint (**Extended Data Fig. 7a, b**), although the match is incomplete in complex I, where no background position is as constrained as the class. We therefore compared, within the loading sites, substitutions that change formal side-chain charge with those at the same residues that do not.

In complex I, charge-changing substitutions at loading sites relaxed 10.84-fold (7.99–15.82) and charge-preserving substitutions at the same residues 5.30-fold (3.81–7.47), a ratio of 2.05 (1.34–3.27; one-sided P = 0.002; **Fig. 4b, c**). Within tumours alone the two classes had similar dN/dS (1.12 and 1.10; ratio 1.02, 0.86–1.21), indicating that the strong differential constraint observed in the germline is substantially attenuated in tumours. By contrast, substitutions severing the wire at cut vertices relaxed no more than others (0.87, 0.54–1.38). The contrast was 2.32 (1.42–3.84) with gnomAD and stayed above 1 in its point estimate under every analysis choice, with each ND gene removed, with coverage weighting (**Supplementary Fig. 7**) and without the 46 calls of the most recurrent substitution, m.3380G>A (1.94, 1.27–3.12). With heteroplasmic-only germline variants, which have had little time to be filtered, as the reference it fell to 1.18 (0.59–2.25), and within tumours alone the two classes differed in TCGA (1.50, 1.01–2.43) but not in PCAWG (0.83, 0.64–1.09) (**Extended Data Fig. 7c**). The hydration model can dilute this contrast but not create it: checked against a mouse complex I structure that resolves its waters ^29^, it recovers 94.9% of buried waters and 88 of 90 loading sites, and at the 74 human loading sites that the observed waters confirm after alignment the ratio is 3.27 (1.32–8.27; **Extended Data Fig. 8**).

The same avoidance of charge is visible on the timescale of human evolution. Of the 14 ND missense substitutions at complex I loading sites that define branches of the human mtDNA phylogeny ^30^, 13 preserve formal charge, mostly exchanging one titratable residue for another; only ND4 R340H changes it, against 56% expected (P = 2 × 10 ; **Fig. 4d**). Substitutions carried by more than ten HelixMTdb individuals behave alike (3 of 18; P = 7 × 10), whereas loading-site substitutions recurrent in four or more tumours change charge more often than expected (62 of 69 against 80%; P = 0.02; **Supplementary Table 4**). These recurrences do not exceed mutational opportunity and are no more clonal than other ND missense variants (**Supplementary Fig. 8**), which fits tolerance rather than positive selection. The same positions matter in inherited disease: 8 of the 29 confirmed pathogenic ND missense variants in MITOMAP are charge-changing substitutions at loading sites, against 6% expected (P = 2 × 10 ; **Supplementary Fig. 9**).

On the proton wires of complexes III and IV the contrast is absent (1.19, 0.62–2.29 and 0.78, 0.50–1.27), and of the eight contrasts, four complexes on two axes, only that of complex I survives Holm correction (adjusted P = 0.032). Charge is nonetheless released in complex III at its electron-transfer cofactors: around the b haems, charge-changing substitutions relaxed 7.36-fold and charge-preserving substitutions at the same residues 2.33-fold, a ratio of 3.17 (1.80–8.26; one-sided P = 0.001), concentrated below a VAF of 0.5 (**Extended Data Fig. 5b**). Because the predicted electron-transfer effect of these substitutions is computed from their electrostatic shift and is almost collinear with it (r = 0.94; **Supplementary Fig. 4c**), the two cannot be separated, and we interpret this pattern as consistent with electrostatic tuning of the haem driving force, although other coupled effects cannot be excluded. At the oxygen-reducing centre of complex IV no charge-changing variant survives in the germline, so the comparison cannot be made. The germline thus avoids, and tumours tolerate, charge changes where it tunes a transfer step.

### A released charge can retune or abolish proton transfer

A charge change can act on a transfer step by shifting the protonation free energy of the donor relative to the acceptor, which moves the wells of the potential, or by changing the geometry of the barrier, in which case rate and isotope sensitivity move together ^13,16^; changes in barrier geometry are expected to alter tunnelling probability and isotope sensitivity more directly, although energetic shifts can also affect transfer. To find out which the released substitutions do, we computed the potential felt by the transferring proton with density-functional theory ^31^ on a cluster at the Asp179–water–Glu145 bridge of MT-ND5, with charge-changing substitutions in tumours at Asp179, Glu145 and the neighbouring Arg176 (Asp179Asn in twelve, Glu145Lys in ten, Arg176His in four, Glu145Gly in three and Glu145Gln in one). We modelled the loss of the distant Arg176 charge (the Arg 176 charge-removal model, a proxy for Arg176His) and Glu145Gln and Glu145Lys, at the accepting group; Asp179Asn and Glu145Gly were not modelled.

At the structural geometry the potential is a double well with a barrier of 10.40 kcal mol ¹ and an acceptor well 1.63 kcal mol ¹ above the donor well (**Fig. 5a, b**). Removing the charge of Arg176, 6.8 Å from the proton (modelled by deleting the guanidinium group, which approximates the electrostatic effect of Arg176His but not its full structural or protonation-state consequences), acts as a classical electrostatic shift: the acceptor well drops to 0.71 kcal mol ¹, the barrier only to 9.84 kcal mol ¹, and the separation of the wells stays at 0.827 Å. Glu145Gln and Glu145Lys instead remove the acceptor well, the potential rising monotonically by 42 and 35 kcal mol ¹, so the step is abolished (**Fig. 5a**). The two kinds of substitution that tumours release are thus distinguishable. With a larger basis set and dispersion correction (B3LYP-D3(BJ)/def2-TZVP), the barrier was 10.9 kcal mol ¹, the Arg176 charge-removal model lowered it by 0.66 kcal mol ¹ and both Glu145 substitutions still abolished the step (**Supplementary Table 1, sheet S2q**).

**Fig. 5.**
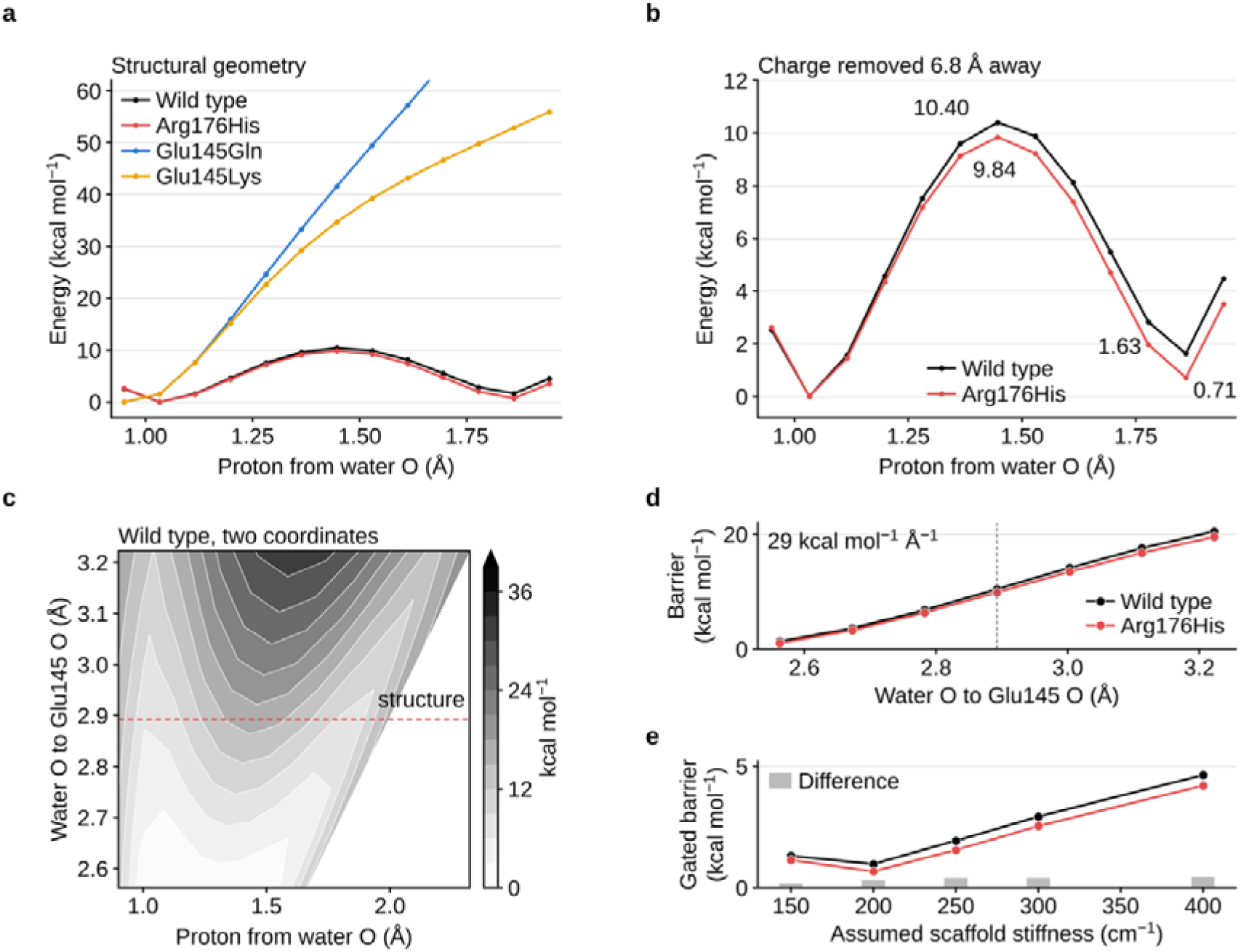
A charge change at a distance shifts the transfer step without changing the shape of its barrier, and the barrier is gated by the width of the hydrogen bond. Density-functional calculations on a cluster taken from PDB 9I4I at the Asp179–water–Glu145 bridge of MT-ND5 (B3LYP/def2-SVP, density fitting, conductor-like continuum with dielectric constant 4). The step shown is the transfer from the hydronium formed in the first, barrierless step to Glu145. a, Potential felt by the transferring proton at the geometry of the structure, for the wild type and three substitutions observed in tumours. Glu145Gln and Glu145Lys remove the acceptor well, and the potential rises monotonically by 42 and 35 kcal mol□¹. b, Wild type and the Arg176 charge-removal approximation of Arg176His, in which the guanidinium charge 6.8 Å from the transferring proton is removed. The barrier falls from 10.40 to 9.84 kcal mol□¹ and the acceptor well from 1.63 to 0.71 kcal mol□¹ above the donor well. The separation of the wells is 0.827 Å in both. c, Wild-type potential against proton position and water-to-Glu145 distance (77 points; 7 separations from 2.56 to 3.22 Å × 11 proton positions), relative to its minimum. The dashed line marks the structural separation (2.89 Å). d, Transfer barrier against the water-to-Glu145 distance for the two charge states. The barrier rises from 1.3 to 20.5 kcal mol□¹ over the sampled range, 29 kcal mol□¹ Å□¹; the dashed line marks the structural separation. e, Height of the gated barrier against the stiffness assumed for the harmonic scaffold term, for the two charge states, with their difference (bars). The 100 cm□¹ setting, at which the relaxed path leaves the sampled range, is omitted. The difference between charge states stays between 0.17 and 0.44 kcal mol□¹ and in the same direction throughout.

Because donor and acceptor approach one another in a protein, we scanned the proton at seven water-to-Glu145 distances (2.56–3.22 Å; **Fig. 5c**). The barrier fell from 20.5 kcal mol ¹ at 3.22 Å to 10.5 at the structural 2.89 Å (10.45 in the scan; 10.40 at the unmodified structural geometry) and 1.3 at 2.56 Å, 29 kcal mol ¹ per ångström (**Fig. 5d**), the signature of gated transfer in which protein motion compresses donor and acceptor into the range where the proton can tunnel ^16^. A second bridge, in MT-ND1, gave 40 kcal mol ¹ Å ¹ and a nearly coincident curve, although its potential has no product well at widths up to 0.11 Å beyond the structure, and a third was barrierless (**Extended Data Fig. 9a**); the tunnelling factor of a fixed geometry (**Extended Data Fig. 9b**) is therefore not a property of the enzyme. With a harmonic scaffold standing in for the protein, the gated barrier was 0.97–4.63 kcal mol ¹ over assumed stiffnesses of 150–400 cm ¹, and the difference between the two charge states stayed at 0.17–0.44 kcal mol ¹ in the same direction (**Fig. 5e**). Averaging proton and deuteron transfer on the scanned surface over a thermal distribution of widths (**Extended Data Fig. 10**), the predicted kinetic isotope effect at 308 K was 7.5 (6.6–10.7 over stiffnesses), against 31 at fixed geometry, falling from 9.6 at 278 K to 7.0 at 318 K with an isotopic difference in activation energy of 1.4 kcal mol ¹ (1.2–1.9). In the model, tunnelling carries most of this transfer: at 250 cm ¹, 69% of the gated proton rate passed below the barrier top (53–88% over stiffnesses; 53% for the deuteron), at a donor–acceptor distance of 2.66 Å, and the rate was 2.4 times (1.7–5.5) that with zero-point energy but no tunnelling. Without tunnelling the ratio of Arrhenius prefactors would be 0.94–0.98, inside the semiclassical range; with it, the ratio fell from 0.92 to 0.44 as the scaffold stiffened (**Fig. 6a, b**). Removing the Arg176 charge raised the proton rate 2.6-fold (2.2–2.9) but changed the isotope effect by only 8–20% and the share of tunnelling little (72%), as expected for a shift of the wells rather than of the barrier. A multidimensional reaction-path treatment, in which the separation relaxes along the path and the tunnelling path can cut the corner, gave tunnelling shares of 84–97%, isotope effects of 4.5–14 and a ratio of Arrhenius prefactors of 0.90–1.08 at stiffnesses of 300–400 cm ¹, whereas at 150–250 cm ¹ it gave isotope effects of 0.8–2.1 and tunnelling shares of 0–50% for the wild type, so in this treatment tunnelling dominates only if the scaffold is stiff; in it removing the Arg176 charge raised the rate 1.6– 2.2-fold and changed the isotope effect by −17% to +31%, so the separation of rate from isotope effect is less clean than in the one-dimensional model (**Supplementary Table 1**). The model rests on one proton coordinate, a cluster and an assumed stiffness, so we report ratios, not absolute rates. Because the thermal distribution of separations weights compressed geometries with low barriers, the apparent activation energy of the gated proton rate is small (−0.06 to 1.2 kcal mol ¹ over stiffnesses) and absolute rates (10¹ –10¹² s ¹) far exceed enzyme turnover; the model describes one transfer step within a cluster, not the rate-limiting step of complex I.

**Fig. 6.**
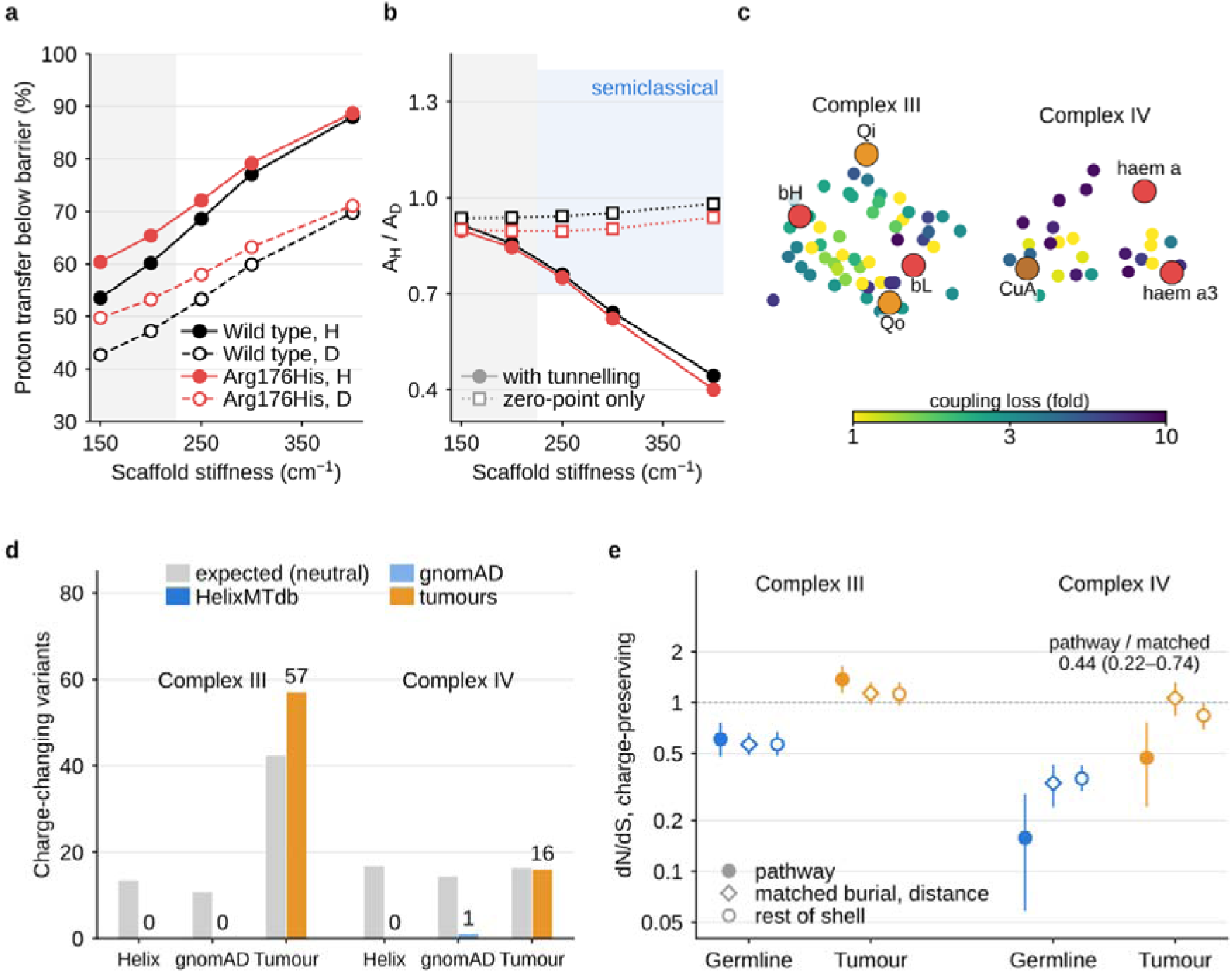
In the one-dimensional gated model, tunnelling contributes substantially to proton transfer, and electron-tunnelling pathways carry constraint beyond their surroundings. a, Share of the gated proton (H, filled) and deuteron (D, open) transfer rate at the MT-ND5 Glu145 bridge that passes below the barrier top at 308 K, against assumed scaffold stiffness, for wild type (black) and the Arg176 charge-removal model (red). b, Ratio of Arrhenius prefactors A_H_/A_D_ over 278–318 K with tunnelling (filled) and with zero-point energy only (open); the blue band marks the semiclassical range (0.7–1.4). Grey regions in a and b mark stiffnesses at which the distance distribution extends past the sampled range. c, Electron-tunnelling pathways projected onto their principal plane: Cα atoms of residues within tenfold of the best Beratan–Onuchic coupling of any step (colour, loss of coupling; metal ligands excluded) and the redox centres. Complex III (PDB 9HZL): Qo → bL, bL → bH, bH → Qi and bL → bL′ (50 MT-CYB residues). Complex IV (PDB 9I7U): CuA → haem a → haem a3 (24 MT-CO1 and MT-CO2 residues). d, Neutral expected (grey) and observed numbers of charge-changing missense substitutions in the pathways, in HelixMTdb (blue), gnomAD (light blue) and tumours (orange). e, Missense dN/dS of charge-preserving substitutions in the pathways (filled), at residues matched on burial and distance to the cofactors (diamonds; three per pathway residue) and at the rest of the 8 Å cofactor shell (open), in the germline (blue) and tumours (orange), with the pathway-to-matched ratio for complex IV in tumours. Error bars show 95% percentile intervals from the shared resamples; log scale in e.

### Electron-tunnelling pathways carry constraint beyond their surroundings

Charge released at the haems tunes the driving force of electron tunnelling; the medium through which the electron tunnels is a separate quantity. We computed Beratan–Onuchic tunnelling pathways ^22^ for every electron-transfer step between redox centres bound by mtDNA-encoded subunits, four in complex III and two in complex IV, taking as each pathway the residues through which the electron can tunnel at no more than a tenfold loss of coupling, metal ligands excluded (Methods): 50 MT-CYB residues and 24 MT-CO1 and MT-CO2 residues (**Fig. 6c**). The germline carries no charge-changing substitution in either pathway, against 13.5 and 16.8 expected under neutrality (Poisson P = 1 × 10 and 5 × 10 ; gnomAD 0 of 10.7 and 1 of 14.3), whereas tumours carry them at or above expectation (57 against 42 and 16 against 16; **Fig. 6d**). Charge changes are, however, also rare in the rest of the 8 Å cofactor shell (germline dN/dS 0.29 in complex III and 0.02 in complex IV). Against residues matched on burial and distance to the cofactors, the complex III pathway carries fewer charge changes than expected (4.4 expected at the matched constraint, none observed; P = 0.0123), a difference that tumours do not keep (pathway/matched 0.92, 0.68–1.19; **Supplementary Fig. 11c and Supplementary Table 1**); in complex IV the matched residues carry none either, so this comparison cannot be made. Charge-preserving substitutions, which can change the tunnelling medium without shifting a redox potential, separate the two complexes. In complex III they behaved in the pathway as at residues matched on burial and distance to the cofactors (germline ratio 1.08, 0.81–1.36). In complex IV the pathway from CuA through haem a to haem a3 was constrained in the germline (0.16, 0.06–0.29) and stayed constrained in tumours (0.47, 0.24–0.76), whereas matched residues relaxed fully (1.06, 0.84–1.32; ratio 0.44, 0.22–0.74; P = 0.001); adjusting for burial, distance and the changes in side-chain volume and hydropathy gave 0.48 (0.24– 0.98), and the result held with the oxygen-reducing centre removed (0.55, 0.26–0.91; **Fig. 6e**). A similar pattern was seen in the bovine structure at 1.5 Å, where the 24 pathway residues of lowest coupling loss (14 shared with the human set; a tenfold threshold there gives only nine residues, with tumour dN/dS 0.73, 0.29–1.32) gave 0.17 (0.08–0.31) in the germline and 0.47 (0.25–0.71) in tumours, and in a permutation test against 10,000 sets of residues matched on burial and distance the human pathway was more constrained than the matched sets in the germline (P = 0.011) and in tumours (P = 0.025). Tumours thus tolerate charge around the redox cofactors, including along the tunnelling pathways, and, in complex IV, keep the medium through which the electron tunnels. The analysis identifies paths of electronic coupling; it does not show that any substitution changes a tunnelling rate.

## Discussion

Relaxed selection on mtDNA in tumours is usually treated as a dimmer switch on the germline filter ^4,5^. It is not: complex V trades places between the extremes, and its relative relaxation stays below 1 under every analysis choice and without the measurement ceiling. One interpretation is that cancer cells tolerate greater impairment of the respiratory complexes that generate the proton-motive force than of ATP synthase, which can either harness this force for ATP production or consume ATP to help maintain membrane potential through reverse operation ^6,32,33^. Damage at the haems of complex III and the oxygen-reducing centre of complex IV is affordable mainly while wild-type genomes predominate; as they recede, the excess disappears at the haems and turns into depletion at the oxygen-reducing centre, so tumours change how strictly the cost of damage is enforced, not the cost itself. Around the haems of complex III, charge-changing substitutions also exceed neutral expectation in tumours (dN/dS 1.66, and 2.08 below a VAF of 0.5), which could reflect positive selection for altered electron transfer at low dosage as well as tolerance; these data do not separate the two. These data cannot say whether the requirement for ATP synthase is for synthesis, for consumption of the gradient or for the integrity of the complex. Other explanations also remain open, including dominant-negative effects of subunit a variants on the assembled enzyme and hyperpolarization with increased production of reactive oxygen species when ATP synthase is impaired.

CRISPR knockouts of nuclear-encoded subunits in 1,361 cancer cell lines ^34^ support this only in part: with the c-subunit paralogues set aside ^35^, ATP synthase is more essential than complex I but not than complexes II–IV (**Supplementary Fig. 10**), and growth in culture is a poor model of the filter acting on mtDNA in a tumour ^36^.

The decisive comparison for the physical question is made within a residue, which removes the alternative that loading sites simply lie where tumours tolerate change. The avoidance of charge on the human phylogeny and the concentration of disease variants in the same class indicate that tumours reverse a long-standing selection rather than exploit a quirk of somatic calling. The same contrast is strong around the haems of complex III ^10,26^ and absent on the proton wires of complexes III and IV, which suggests that what tumours release is not proton translocation as such but particular transfer steps whose rate a single charge can retune, such as the chain of charged residues that couples quinone chemistry to pumping in complex I ^37,38^ and the haem driving force in complex III.

These results connect quantum biology to cancer genomics in two ways, both testable. Electron and proton transfer are the steps at which tunnelling enters bioenergetics ^10,16^. For electrons, tunnelling pathways are only partly specific: the germline carries no charge change along them (and, in complex III, fewer than at matched residues); tumours remove this extra protection in complex III and keep the tunnelling medium itself in complex IV. For protons, the one-dimensional gated model predicts substantial transfer below the barrier at the Glu145 bridge, although the multidimensional estimate depends strongly on scaffold stiffness. Tumours carry substitutions of both kinds at this bridge: removing the distant Arg176 charge shifts the wells while leaving the barrier and the share of tunnelling nearly unchanged, whereas charge changes at Glu145 itself abolish the step. Selection at the level of the class therefore does not distinguish retuning from abolition; the calculations separate them and make the distinction testable. Neither result is a measurement of tunnelling, and the predictions depend on the assumed scaffold stiffness: an isotope effect of about 7–11 that falls with temperature and a ratio of Arrhenius prefactors below 0.7 if the scaffold is stiff (both in the one-dimensional model; 4.5–14 and 0.90–1.08 in the multidimensional one with a stiff scaffold, and smaller isotope effects with a soft one), and, if Arg176His acts by removing the Arg176 charge, an isotope effect changed much less than the rate by this substitution (by 8–20% in the one-dimensional model and by at most a third in the multidimensional one, against a 1.6–2.9-fold change in rate). Because the multidimensional treatment places the prefactor ratio inside the semiclassical range even when tunnelling dominates, the magnitude and temperature dependence of the isotope effect, rather than the prefactor ratio, are the more robust test. Such a temperature-dependent isotope effect would place the step among the enzymatic proton transfers in which protein motion gates tunnelling ^16,39^; MT-ND5 Arg176His, carried by four tumours, is a natural candidate for such a test by solvent isotope effects on complex I isolated from cybrids or from cells edited with mtDNA base editors, although a single step of the proton pathway need not limit turnover.

Several limitations remain. Complex V contributes two short genes, so its subunit-level analyses rest on small counts. The hydration model adds water as well as recovering it, so the loading-site class is a superset of the real one, and cut vertices suffer more. Germline and tumour data differ in ascertainment, which per-dataset nulls and alternative references address without making them interchangeable. Cross-sectional VAF does not measure a biochemical threshold or longitudinal purging and may reflect copy number and subclonal structure. VAF distributions also differ between cohorts (median VAF of ND missense calls about 0.04 in PCAWG against about 0.5 in TCGA and MSK-IMPACT), but the dependence on VAF persists when each cohort is divided into thirds of VAF and within single cohorts, except for complex I in TCGA and, for complex III, when TCGA alone is split at a VAF of 0.5. Low-VAF calls are also enriched for sequencing and DNA-damage artefacts, which would pull dN/dS towards 1 in the lowest stratum; at the oxygen-reducing centre of complex IV, where no within-residue control is available, this could contribute to the weaker depletion at low VAF. Conversely, the most recurrent step-abolishing substitution, MT-ND5 Glu145Lys, is carried at a median VAF of 0.73, which the dosage-buffering reading does not explain. The tunnelling analyses identify coupling paths and predict ratios from one bridge, one proton coordinate and an assumed stiffness, not rates. Finally, dN/dS measures the filter that acted on a set of variants, not the fitness effect of any one substitution ^40^, and none of these results has been tested functionally.

## Methods

### Variant data

Somatic mtDNA variants and HelixMTdb population variants ^41^ were taken from the harmonised resource of Gorelick and colleagues, which assembles TCGA, PCAWG and MSK-IMPACT calls under a common pipeline ^6,24,25^. We retained single-nucleotide variants between rCRS positions 577 and 16,023 whose reference allele matched rCRS ^42^, restricted MSK-IMPACT and TCGA calls to those classified as somatic or likely somatic, and deduplicated by patient, position and alternative allele within and across cohorts, so that a variant called in both TCGA and PCAWG for one of the 187 shared donors is counted once, from TCGA. This gave 16,152 variants, 11,435 in coding sequence, in 8,886 tumours (TCGA 2,300 patients, PCAWG 2,184, MSK-IMPACT 4,589 before removal of the 187 shared donors). Population variants (12,224 sites) were filtered identically and treated as distinct sites with heteroplasmic and homoplasmic carrier counts. The population arm was repeated with gnomAD v3.1 chrM (56,434 genomes) ^3^, discarding variants failing release filters (9,324 usable sites). No germline-polymorphism filter was applied to somatic calls; deduplication and the somatic classification of the resource were used instead.

### Annotation and mutational null

Consequences were assigned under the vertebrate mitochondrial code ^43^. Every possible single-nucleotide change at every coding position was enumerated (33,786 changes), excluding positions in overlapping reading frames (MT-ATP8/MT-ATP6, 8,527–8,572; MT-ND4L/MT-ND4, 10,760–10,766). Each change was assigned to one of 192 classes defined by its L-strand trinucleotide context and alternative allele ^44^. For each dataset separately, the per-site rate of a class was the number of observed variants in that class, coding and non-coding, divided by the number of reference positions with that context, and the expected count for any group of changes is the sum of these rates. Estimating the spectrum from each dataset in turn keeps the strand asymmetry of mtDNA mutation from propagating into selection estimates (**Supplementary Fig. 1**). Synonymous calibration in every structural class is shown in **Supplementary Fig. 2**.

### dN/dS, relaxation and resampling

Missense dN/dS is the observed-to-expected ratio of missense changes divided by that of synonymous changes in the same genes ^40,45^. The fold change is tumour dN/dS divided by germline dN/dS, and the relative relaxation of complex V its fold change divided by the geometric mean of those of complexes I, III and IV. Ranks across the four complexes and thirteen genes were recomputed in every resample. All intervals and P values come from one set of 1,000 bootstrap resamples per dataset, generated once from a fixed seed and shared by every analysis; tumour datasets were resampled by patient and population datasets by site. Intervals are 2.5th and 97.5th percentiles, and one-sided P values are (replicates at or beyond the null + 1)/(finite replicates + 1). Sensitivity analyses changed one choice at a time: tumour cohort, VAF stratum, germline reference, null model (synonymous and non-coding variants only), removal of sites recurrent in five or more tumours, and gene sets. The analysis by cancer type used TCGA and MSK-IMPACT, for which harmonised diagnoses are available, and cancer types with at least 400 substitutions. Truncating dN/dS was computed in the same way for nonsense changes (**Extended Data Fig. 3**).

### Controls for the ceiling

First, missense dN/dS of complex V was compared with that of complexes I, III and IV, and with that of MT-CO2 and MT-CO3, within the tumour data alone. Second, population variants were split at random into two halves with a fixed seed; each gene was divided into three positional bins, the bins of complexes I, III and IV in the top quartile of germline dN/dS in the first half were selected, and their germline and tumour dN/dS were estimated on the held-out half. The same design with ten bins per gene matched background positions to the germline constraint of each wire class (**Extended Data Fig. 7b**).

### Cofactor environments of complexes III and IV

The haem b environment comprises the 94 MT-CYB residues of PDB 9HZL (chain J) with any heavy atom within 8 Å of either b haem, after removal of residues within 4.5 Å of the bound quinones (Qo and Qi sites) or of the Rieske chains; a 6 Å radius, PDB 9CG3 and the residue set of our earlier analysis were used as sensitivity analyses. The oxygen-reducing centre comprises the 24 MT-CO1 residues of PDB 9I7U (chain A) with any heavy atom within 8 Å of the haem a3 iron or CuB; PDB 5B1A (bovine, 1.5 Å), whose numbering coincides with the human sequence at every catalytic residue, gave 26. Residues lining the gas-accessible cavity of complex IV were those within 5 Å of the four xenon sites within 25 Å of the haem a3 iron in PDB 9KUK, excluding the centre. The predicted effect of each MT-CYB substitution on three Q-cycle electron-transfer steps was computed in PDB 9CG3 from the screened Coulomb shift of its net charge change at each redox centre, the non-adiabatic Marcus expression ^26^ and Pathways electronic couplings ^22^, with an effective dielectric constant of 8 (4–16 in sensitivity analyses).

### Proton-wire graphs and the hydration model

Graphs were built with identical parameters for the mtDNA-encoded subunits of PDB 9I4I (complex I; chains s, i, j, r, k, l and m for MT-ND1, -ND2, -ND3, -ND4, -ND4L, -ND5 and - ND6), 9HZL (complex III; chain J), 9I7U (complex IV; chains A–C) and 8H9F (ATP synthase; chains N and Q) ^21^. Nodes are Grotthuss-competent side-chain heteroatoms ^27^ of these subunits and of any residue of another chain within 6 Å, together with water positions; edges join nodes 2.2–3.5 Å apart, excluding pairs within one residue or adjacent residues. Connected components of at least four nodes are clusters, and are major if they have at least 12 nodes or span at least 10 Å along the membrane normal, fitted from the hydrophobic belt of each structure. Loading sites are titratable residues (Asp, Glu, His, Lys, Arg, Tyr, Cys) with a node in a major cluster; cut vertices are residues with a node that is an articulation point of its component. Because none of the four structures models water, cavity water was generated with a buried-cavity model on a 0.7 Å grid within 12 Å of the target subunits, retaining points at least 2.6 Å from protein heavy atoms, within 3.4 Å of a nitrogen or oxygen and with at least 75 heavy atoms within 8 Å, thinned to a minimum spacing of 2.5 Å ^29^. The model was checked on mouse complex I (PDB 8OM1, 2.39 Å) by building the graph from modelled and from observed buried waters and transferring confirmed loading sites to human residues by Needleman–Wunsch alignment with BLOSUM62 ^46,47^. The ATP synthase graph was rebuilt over nine combinations of edge cutoff and burial threshold.

### Within-site contrasts

A substitution changes charge if the formal side-chain charge differs between wild type and mutant (Lys and Arg +1, Asp and Glu −1, others 0), and severs the wire if it replaces a Grotthuss-competent residue with one that is not. At loading sites, charge-changing substitutions were compared with charge-preserving substitutions at the same residues, and at cut vertices severing with non-severing substitutions, each against the synonymous changes of the same genes; we report the ratio of their tumour-to-germline fold changes. The eight within-site contrasts (four complexes, two mechanistic axes) were treated as one family for multiple-testing correction, and family-wise P values are two-sided and Holm-adjusted. The same contrast was applied to the haem b environment. Robustness analyses removed each ND gene in turn from both the class and the synonymous denominator, removed the 46 calls of m.3380G>A, and weighted expected counts in TCGA by the number of tumours callable at each position.

### Electron-tunnelling pathways

Electronic couplings were computed with the Beratan–Onuchic Pathways model ^22^ on the heavy atoms of the structures of the main analysis, with a decay of 0.6 per covalent bond, 0.6² exp[−1.7(R − 2.8)] for hydrogen bonds between N and O atoms 2.5–3.5 Å apart and 0.3 exp[−1.7(R − 1.4)] for through-space jumps up to 6 Å. Donors and acceptors were the metal and conjugated porphyrin atoms of each haem, the copper ions and the quinone head groups. The steps were Qo quinone → haem bL, haem bL → haem bH, haem bH → Qi quinone and haem bL → haem bL′ of the other monomer in PDB 9HZL, and CuA → haem a and haem a → haem a3 in PDB 9I7U; haem bL and haem a3 were identified as the haems nearest the Qo quinone and CuB. The best path was found with Dijkstra’s algorithm on −ln ε, and each residue was assigned the loss of coupling of the best path forced through any of its atoms. The pathway of a step comprises the mtDNA-encoded residues with a loss of at most tenfold (threefold and thirtyfold in sensitivity analyses), excluding residues with an N, O or S atom within 2.6 Å of a metal. Charge-preserving and charge-changing missense substitutions in the pathways were compared with those at the other residues within 8 Å of the same cofactors, with dN/dS against the synonymous changes of the genes of each complex and the shared resamples. For classes with no observed germline variant, the expected count under neutrality is the spectrum-weighted opportunity scaled by the synonymous observed-to-expected ratio, with a Poisson probability of observing none. Matched controls were, for every pathway residue, the three nearest non-pathway residues of the same genes in standardised burial (heavy atoms within 10 Å of the Cβ) and minimum distance to a redox cofactor, drawn without replacement. The same matched residues were used for charge-changing substitutions; with no germline variant on the pathway, the test is the Poisson probability of observing none given the neutral expectation on the pathway multiplied by the dN/dS of the matched residues. In a quasi-Poisson regression over every possible charge-preserving missense change of the genes, with the spectrum-weighted expectation as offset, the pathway term was adjusted for burial, distance and the absolute changes in side-chain volume and Kyte–Doolittle hydropathy. For replication, pathways were recomputed in PDB 5B1A (bovine complex IV) and PDB 9CG3 (human complex III, without quinones), carrying bovine residues to human numbering only where the residue was identical at the same number (91% of MT-CO1 and 73% of MT-CO2 positions) and taking the same number of residues as in the main structure. For the permutation test, each pathway residue was replaced by a random draw from its ten nearest non-pathway residues in standardised burial and distance, 10,000 times, and the empirical P value is the fraction of sets with dN/dS at or below that of the pathway.

### Heteroplasmy and purity

Dependence on heteroplasmy was assessed by dN/dS within VAF strata (<0.1, 0.1–0.5 and ≥0.5, and either side of 0.5), with mutation rates from the whole tumour spectrum, observations restricted to the stratum and intervals from the shared patient-level resamples. Among the missense variants of each gene, membership of a class was regressed on VAF by logistic regression with cohort as a covariate. In TCGA, VAFs were divided by the consensus purity estimate ^48^ and capped at 1. To separate VAF from cohort, the calls of each cohort were also divided into thirds of VAF, with the same rates and resamples, and the logistic models were refitted with a cohort-by-VAF interaction, with VAF as a within-cohort percentile, and within each cohort. The expected number of oxygen-reducing-centre variants among the 63 MT-CO1 missense mutations of MSK-IMPACT is their number multiplied by the spectrum-weighted fraction of MT-CO1 missense opportunity at the centre, with a Poisson probability for the observed count. Germline heteroplasmy was compared between sites seen only in the heteroplasmic state and sites homoplasmic in at least one carrier by two-sided Fisher’s exact tests.

### Phylogeny, recurrence and disease variants

Haplogroup-defining substitutions of PhyloTree build 17 ^30^ were mapped to possible coding changes, and the fraction of those at complex I loading sites that change formal charge was compared with the fraction expected from the germline spectrum over the same residues by a binomial test; the same test was applied to HelixMTdb substitutions carried by more than ten individuals and to tumour substitutions recurrent in four or more patients. Expected recurrence came from 2,000 multinomial draws of each cohort’s ND missense calls over all possible ND missense substitutions, weighted by the tumour spectrum. MITOMAP ND missense variants ^49^ (release of 25 May 2022), all reported and confirmed pathogenic, were assigned to wire classes and compared with spectrum-weighted expectation by binomial tests.

### Structural regions of ATP synthase and essentiality

MT-ATP6 and MT-ATP8 residues of PDB 8H9F were classified as the Arg159 shell, the a/c-ring interface, the stator interface, lipid-exposed ^50,51^ (relative accessibility ≥0.25) or buried core, at shell radii of 6–12 Å, with a family-wise permutation test over the nine definitions, and the analysis was repeated on PDB 8H9S. CRISPR gene effects of nuclear-encoded structural subunits were taken from DepMap ^34^ and compared between complexes by one-sided Mann–Whitney tests, with and without the three c-subunit paralogues.

### Density-functional calculations

Clusters were built from PDB 9I4I at the positions given by the structure. At the MT-ND5 bridge, Asp179 and Glu145 were modelled as formate, Arg176 as guanidinium, and the bridging water at its modelled position as hydronium carrying the proton transferred in the first, barrierless step. Glu145Gln and Glu145Lys were modelled as amide and ammonium on the same carbon, and the electrostatic component of Arg176His by removing the guanidinium charge, without modelling the full structural or protonation-state consequences of the substitution. Energies were computed with B3LYP ^52,53^ and def2-SVP ^54^ in PySCF ^55^, with density fitting and a conductor-like continuum of dielectric constant 4 ^56^. As a check of basis set and dispersion, single points at the structural geometry (the reactant well, three points bracketing the barrier top and the acceptor well of the scan for wild type and the charge-removal model, and three points for each Glu145 substitution) were recomputed at B3LYP-D3(BJ)/def2-TZVP on the same clusters, after confirming that the released code reproduces the def2-SVP barrier of 10.40 kcal mol ¹. The two-dimensional surface translated the Glu145 fragment rigidly to seven separations (2.56–3.22 Å) and scanned the proton at eleven positions at each. A harmonic scaffold restraint, calibrated so that the potential is stationary at the structural separation with the proton in its reactant well, stood in for the protein, with stiffness varied from 150 to 400 cm ¹. The MT-ND1 Glu202–water–Ser209 and MT-ND4L/MT-ND2 Glu70–water–Glu34 bridges were treated in the same way. Fixed-geometry nuclear motion was solved on a Colbert–Miller grid ^57^ and tunnelling factors estimated semiclassically at 310 K; these are reported only to show what a fixed-geometry treatment yields. The fixed-geometry isotope effect is 65 from the potential at the structural geometry, 33 from the surface scan and 31 from the rate grid at 308 K (**Supplementary Table 1**), a further reason these values are not interpreted (**Extended Data Fig. 9b**).

### Gated proton tunnelling

For each separation R of the MT-ND5 surface and each charge state, the 11-point proton potential was interpolated with a cubic spline and closed with quadratic walls. Reactant vibrational levels for the proton and the deuteron were obtained by a discrete-variable representation ^57^ confined between the inner wall and the barrier top. The rate at fixed R was k(R) = (hQ_r_) ¹ ∫ P(E) exp(−E/k_B_T) dE, where Q_r_ is the partition function of the reactant levels and P(E) is the semiclassical (WKB) transmission below the barrier top, the parabolic-barrier transmission above it and zero below the product minimum. ln k(R) and the reactant-well energy E_r_(R) were interpolated over R, and the gated rate was the average of k(R) over w(R) exp{−[½K(R − R_c_)² + E_r_(R)]/k_B_T_eff_}, with T_eff_ = (ħω/2k_B_)coth(ħω/2k_B_T) for a donor– acceptor mode of frequency ω ^39,58^. K and R_c_ are the force constant and restraint centre used at each stiffness in the density-functional scan (**Fig. 5e**). Stiffnesses of 150, 200, 250, 300 and 400 cm ¹ and temperatures from 278 to 318 K in 5 K steps were sampled, and activation energies and prefactors came from linear fits of ln k against 1/T. At 150 and 200 cm ¹, 13% and 3% of the distance weight lay at the edge of the sampled range. We report the isotope effect, its temperature dependence and ratios between charge states; absolute rates depend on the single proton coordinate, the adiabatic and semiclassical approximations, the cluster model and the assumed stiffness, and are not interpreted. To measure the contribution of tunnelling, the same averaging was applied to the part of the flux below the barrier top and to a semiclassical rate with the same quantised reactant levels but unit transmission above the barrier top and none below it; the ratio of the full to the semiclassical rate is the tunnelling factor, and the rate-weighted mean of R is the distance at which transfer takes place. As a multidimensional check, the minimum-energy path was found on the surface plus the scaffold term in mass-weighted proton and separation coordinates, with the reduced mass of the separation mode given by K = μω²; the potential along the path was raised by the zero-point energy of the transverse mode, and rates were computed from reactant levels along the path with zero-curvature and small-curvature tunnelling ^59^. Results of this treatment are reported at every assumed stiffness (150–400 cm ¹).

### Statistics and reproducibility

No statistical method was used to predetermine sample size; all available variants meeting the filters were analysed. Unless stated otherwise, intervals are 95% bootstrap percentile intervals from the shared resamples and P values are one-sided bootstrap probabilities in the direction stated, used where the comparison tests a directional hypothesis stated in the text (two-sided values are twice these); Fisher’s exact, binomial, Poisson and Mann–Whitney tests are as described above. Every estimate, with its interval and the counts it rests on, is given in Supplementary Table 1. Multiplicity correction was applied to the family of eight within-site axis contrasts (Holm); the analyses of cofactor environments, electron-tunnelling pathways and their controls were not part of the pre-specified analysis plan and are reported with nominal P values.

## Data availability

Somatic and germline variant tables are available from github.com/reznik-lab/mtdna-mutations and the gnomAD v3.1 mtDNA release from gnomad.broadinstitute.org. Structures are available from the Protein Data Bank under accessions 9I4I, 9HZL, 9I7U, 8H9F, 8H9S, 8OM1, 9CG3, 5B1A and 9KUK; CRISPR data from depmap.org; PhyloTree build 17 (phylotree.org) and the MITOMAP disease variants (mitomap.org; release of 25 May 2022) as distributed at github.com/leklab/mitochondrial_constraint [2]; and TCGA purity estimates from ref. 48. The density-functional surfaces on which the tunnelling calculations rest are given in **Supplementary Table 1**. Source data for every figure are provided in **Supplementary Table 1**.

## Code availability

All analysis code, including the density-functional and gated-tunnelling calculations, is available to editors and reviewers during peer review through a private link provided with the submission and will be deposited on GitHub and archived at Zenodo upon publication; it reproduces every value reported here from the public inputs listed above.

## Acknowledgements

We thank the TCGA Research Network, the ICGC/TCGA PCAWG Consortium and the Helix team for making their data publicly available, and Gorelick and colleagues for releasing their harmonised mtDNA calls. This work was supported by the Korean ARPA-H Project through the Korea Health Industry Development Institute (KHIDI), funded by the Ministry of Health & Welfare, Republic of Korea (RS-2025-25456722), and by the ANCHOR through the Seoul ANCHOR Center, funded by the Ministry of Education and the Seoul Metropolitan Government (2026-ANCHOR-01-022-05), to J.-H.C.

## Author contributions

**J.-Y.S.** and **J.-H.C.** conceived and designed the study. **J.-Y.S.** performed all analyses, interpreted the results, generated the figures and wrote the manuscript. **J.-H.C.** provided the conceptual framework linking quantum-mechanical principles of mitochondrial bioenergetics to human genetic variation and cancer biology, acquired funding and critically reviewed the manuscript. **J.-Y.S.** and **J.-H.C.** jointly supervised the study.

## Additional information

Supplementary Information is available for this paper. Correspondence and requests for materials should be addressed to Ji-Yong Sung or Jae-Ho Cheong.

---

**Extended Data Fig. 1.**
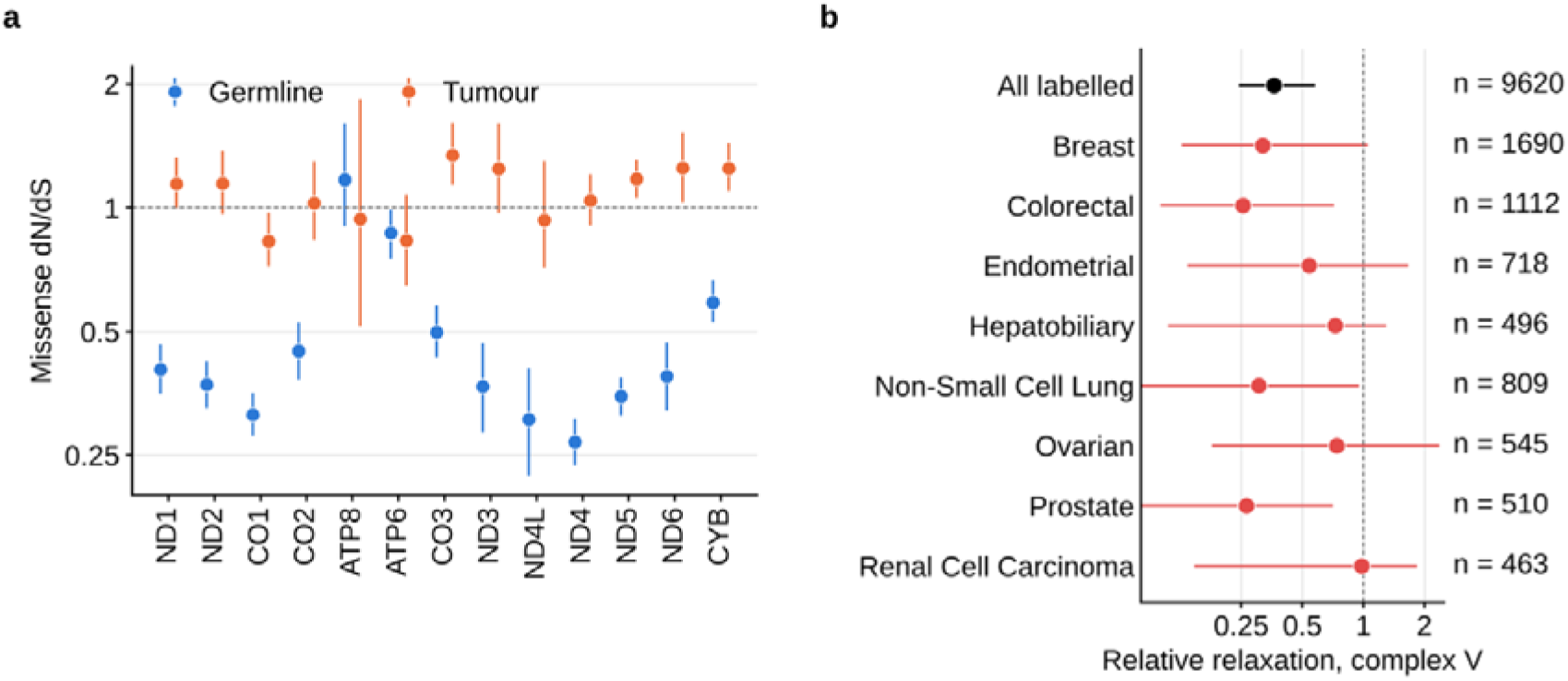
Gene-level estimates and the exemption of complex V by cancer type. a, Missense dN/dS of each of the thirteen mtDNA-encoded genes in the germline (blue) and pooled tumours (orange). b, Relative relaxation of complex V within each cancer type with at least 400 substitutions, using TCGA and MSK-IMPACT, for which harmonised diagnoses are available. The estimate lies below 1 in every type, from 0.25 in colorectal cancer to 0.98 in renal cell carcinoma, and its interval includes 1 in breast, endometrial, hepatobiliary, ovarian and renal cell cancers; all labelled tumours together give 0.36 (0.24– 0.58). n, tumour single-nucleotide variants. Error bars show 95% percentile intervals; log scales.

**Extended Data Fig. 2.**
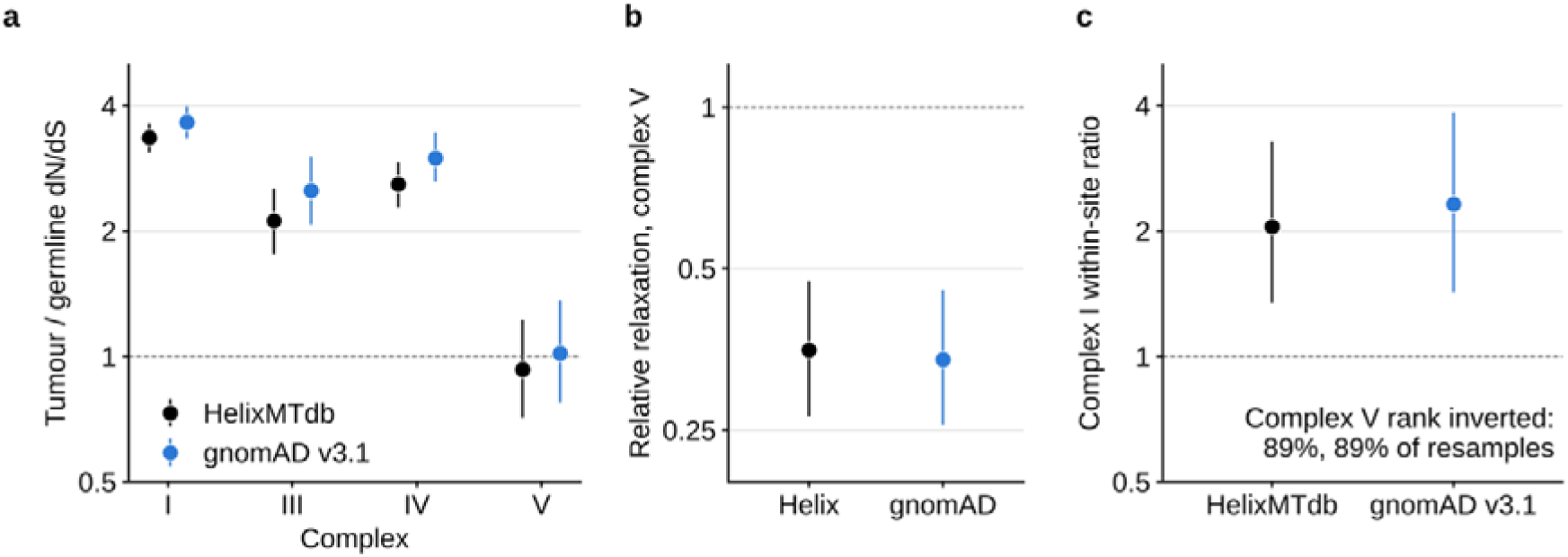
Replication with gnomAD v3.1 as the population reference. The population arm was repeated with the gnomAD v3.1 chrM release (56,434 genomes; 9,324 usable variant sites) under the same annotation, null model, bootstrap and contrasts as HelixMTdb. a, Tumour-to-germline fold change in missense dN/dS by complex, with HelixMTdb (black) or gnomAD (blue) as the germline reference. b, Relative relaxation of complex V: 0.35 (95% CI 0.27–0.47) with HelixMTdb and 0.34 (0.26–0.46) with gnomAD; P ≤ 0.001 for both. c, Within-site charge contrast at complex I loading sites: 2.05 (1.34–3.27) with HelixMTdb and 2.32 (1.42–3.84) with gnomAD. The text gives the fraction of resamples in which complex V was both least constrained in the germline and most constrained in tumours. Error bars show 95% percentile intervals from 1,000 bootstrap resamples. Dashed lines mark a ratio of 1. The y axes are on a log scale.

**Extended Data Fig. 3.**
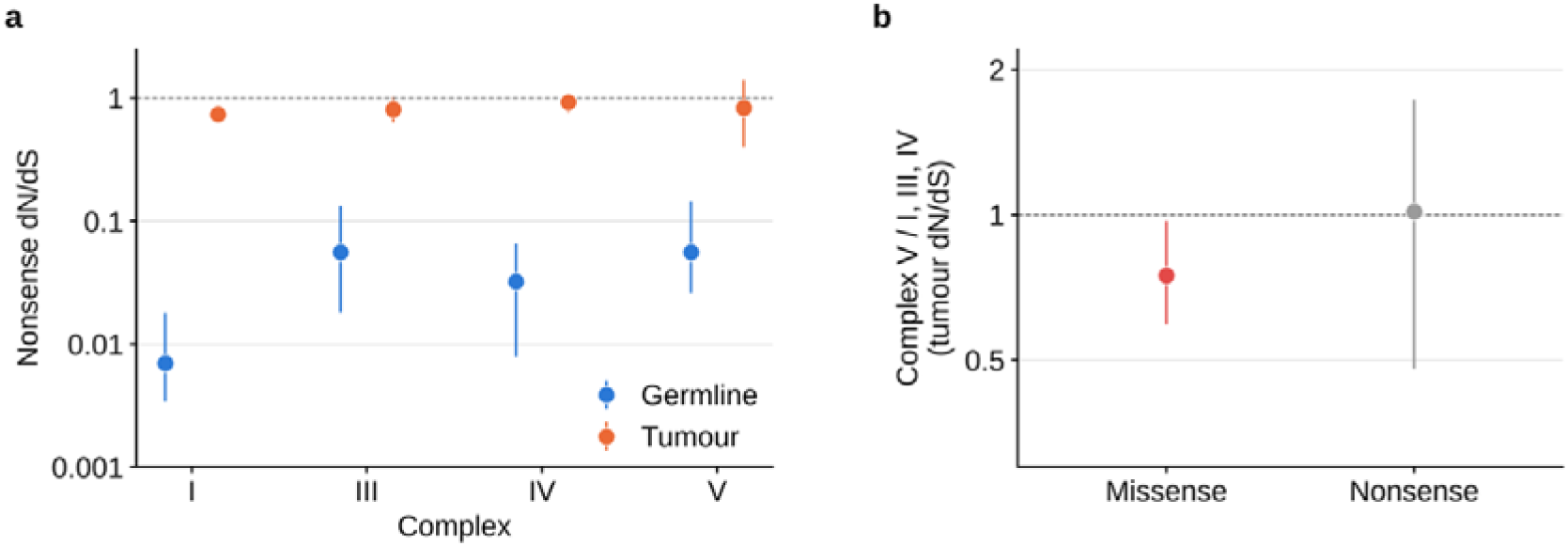
Truncating substitutions. a, Nonsense dN/dS by complex in the germline and pooled tumours. Truncating substitutions are strongly purged from the germline in every complex (0.007–0.056) and relaxed in tumours in every complex, complex V included (0.84, 0.40–1.42, from 14 substitutions). b, Within-tumour ratio of complex V to complexes I, III and IV for missense (0.75, 0.59–0.97) and nonsense (1.01, 0.48–1.73) dN/dS. There is no evidence that tumours purge truncating complex V substitutions, and little power to detect it. Error bars show 95% percentile intervals; log scales.

**Extended Data Fig. 4.**
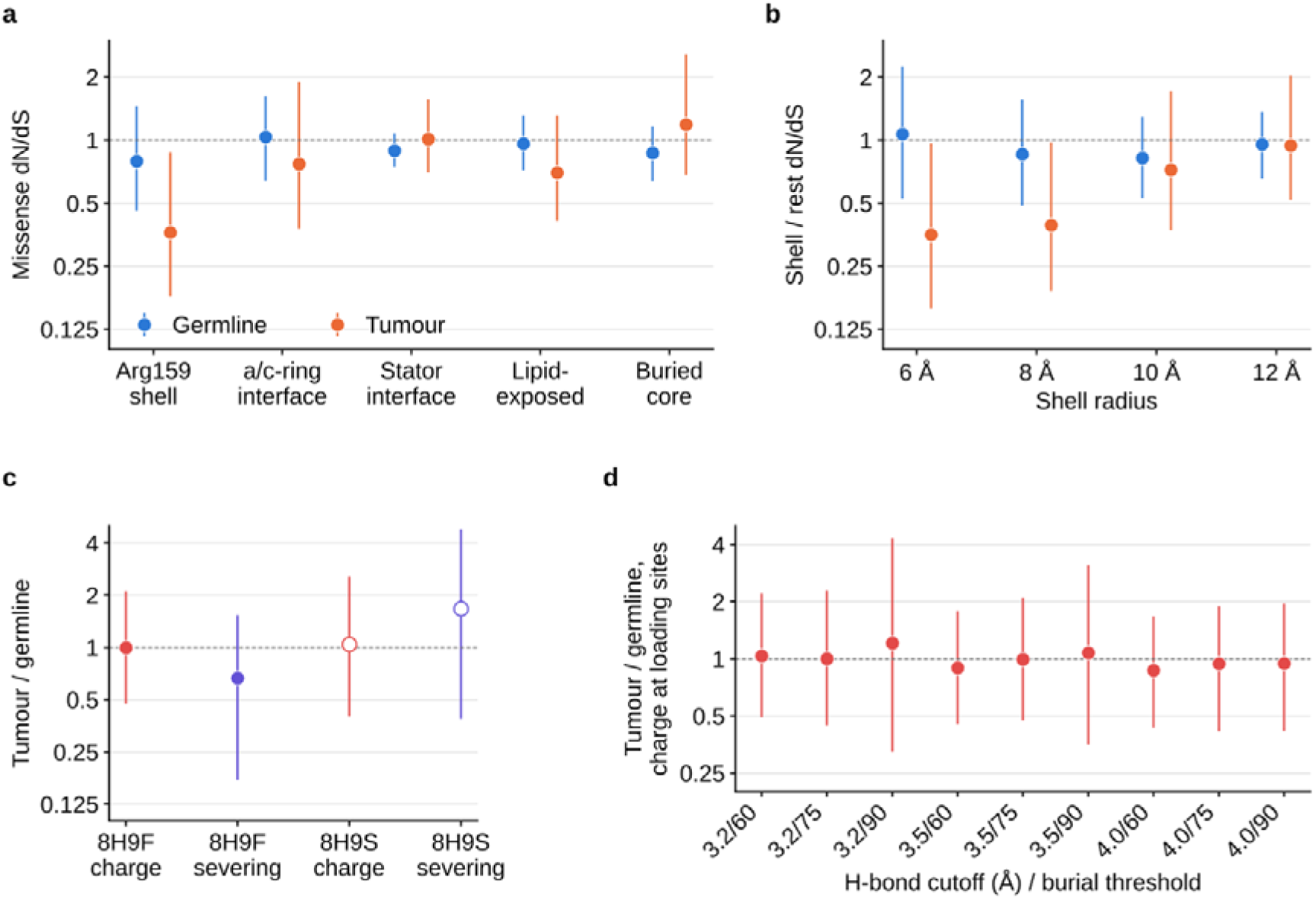
Tumour constraint in subunit a around Arg159. a, Missense dN/dS of MT-ATP6 and MT-ATP8 by structural region (PDB 8H9F), in the germline and pooled tumours. Each residue took the first region it qualified for, in this order: residues within 8 Å of the Arg159 guanidinium, a/c-ring interface, stator interface, lipid-exposed, buried core. Tumour dN/dS in the Arg159 shell is 0.36 (0.18–0.87). b, Shell-to-rest ratio at radii of 6, 8, 10 and 12 Å. The 8 Å tumour ratio (0.39, 0.19–0.98) does not survive a family-wise permutation test over the nine region definitions examined (P = 0.076). c, Tumour-to-germline fold change for charge-changing substitutions at loading sites and wire-severing substitutions at cut vertices of ATP synthase, from PDB 8H9F (filled) and the independent structure PDB 8H9S (open). d, The loading-site fold change after rebuilding the graph under nine combinations of hydrogen-bond cutoff and burial threshold. Error bars show 95% percentile intervals; log scales.

**Extended Data Fig. 5.**
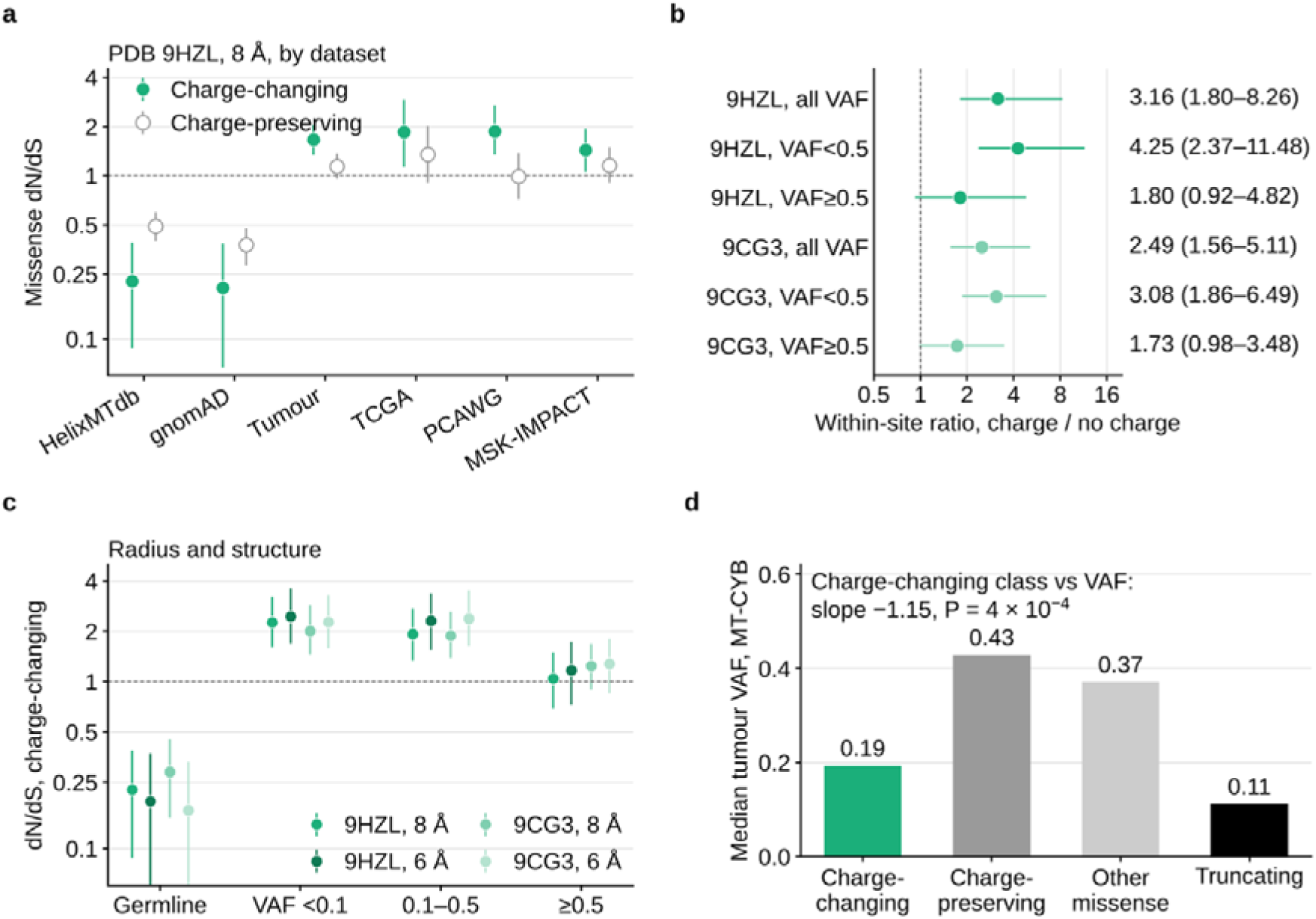
Complex III: charge-specific relaxation around the b haems. Residues of MT-CYB (PDB 9HZL, chain J) with any heavy atom within 8 Å of either b haem, after removal of residues within 4.5 Å of the bound ubiquinones or the Rieske chains (94 residues). a, Missense dN/dS of charge-changing (filled) and charge-preserving (open) substitutions at these residues, by dataset. b, Within-site ratio (charge-changing over charge-preserving fold change) for all tumour variants and below or above a variant allele fraction (VAF) of 0.5, in PDB 9HZL and in the independent structure PDB 9CG3. c, dN/dS of charge-changing substitutions in the germline and by tumour VAF, at radii of 6 and 8 Å in both structures. d, Median tumour VAF of MT-CYB missense substitutions by class, and of truncating substitutions. The text gives the cohort-adjusted logistic slope of charge-changing class membership on VAF. Error bars show 95% percentile intervals from the shared resamples; log scales in a–c.

**Extended Data Fig. 6.**
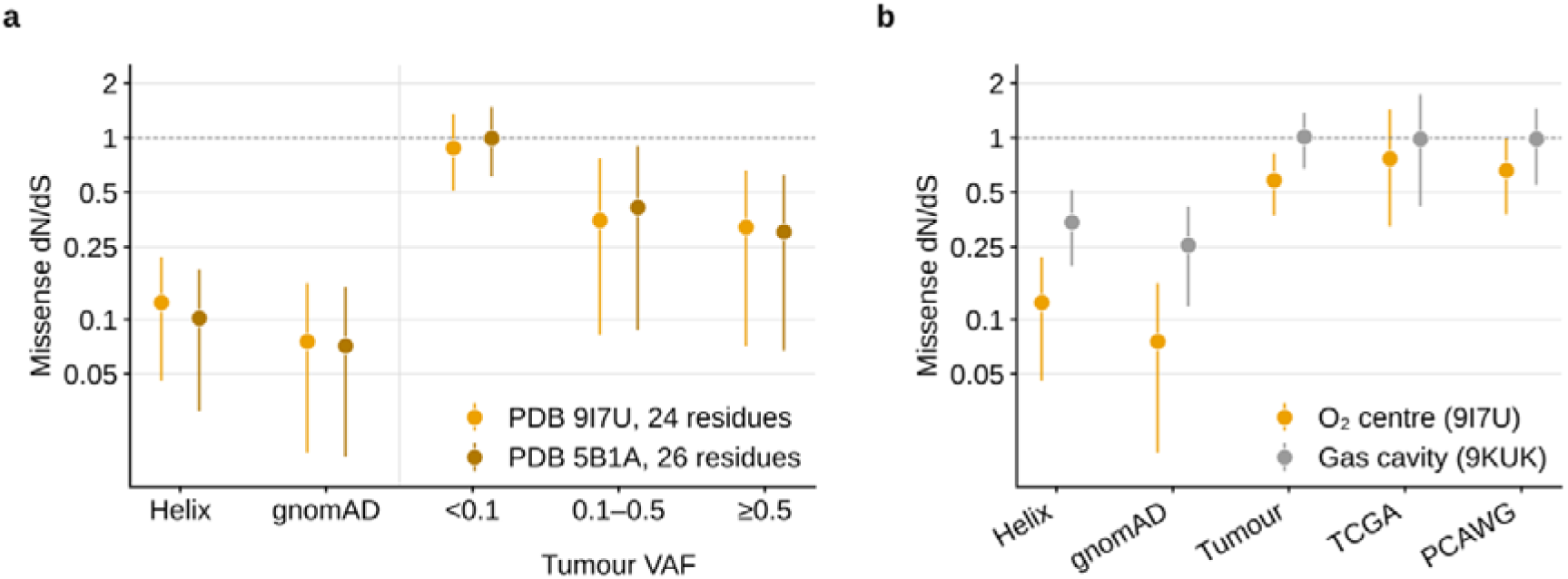
Complex IV: the oxygen-reducing centre in two structures, and the gas-accessible cavity. a, Missense dN/dS of MT-CO1 residues within 8 Å of the haem a3 iron or CuB, in the human structure (PDB 9I7U, 24 residues) and the bovine structure at 1.5 Å (PDB 5B1A, 26 residues; numbering coincides with the human sequence at every catalytic residue). Values are shown in both population references and by tumour VAF. b, The same centre (9I7U) beside the residues lining the gas-accessible cavity. The cavity is defined by the four xenon sites within 25 Å of the haem a3 iron in PDB 9KUK (23 residues within 5 Å of them, excluding the centre). Under the unified null the cavity is itself constrained in the germline (0.34, 0.20–0.52) and unconstrained in tumours (1.01, 0.68–1.38), so it does not serve as a negative control. Error bars show 95% percentile intervals; log scales.

**Extended Data Fig. 7.**
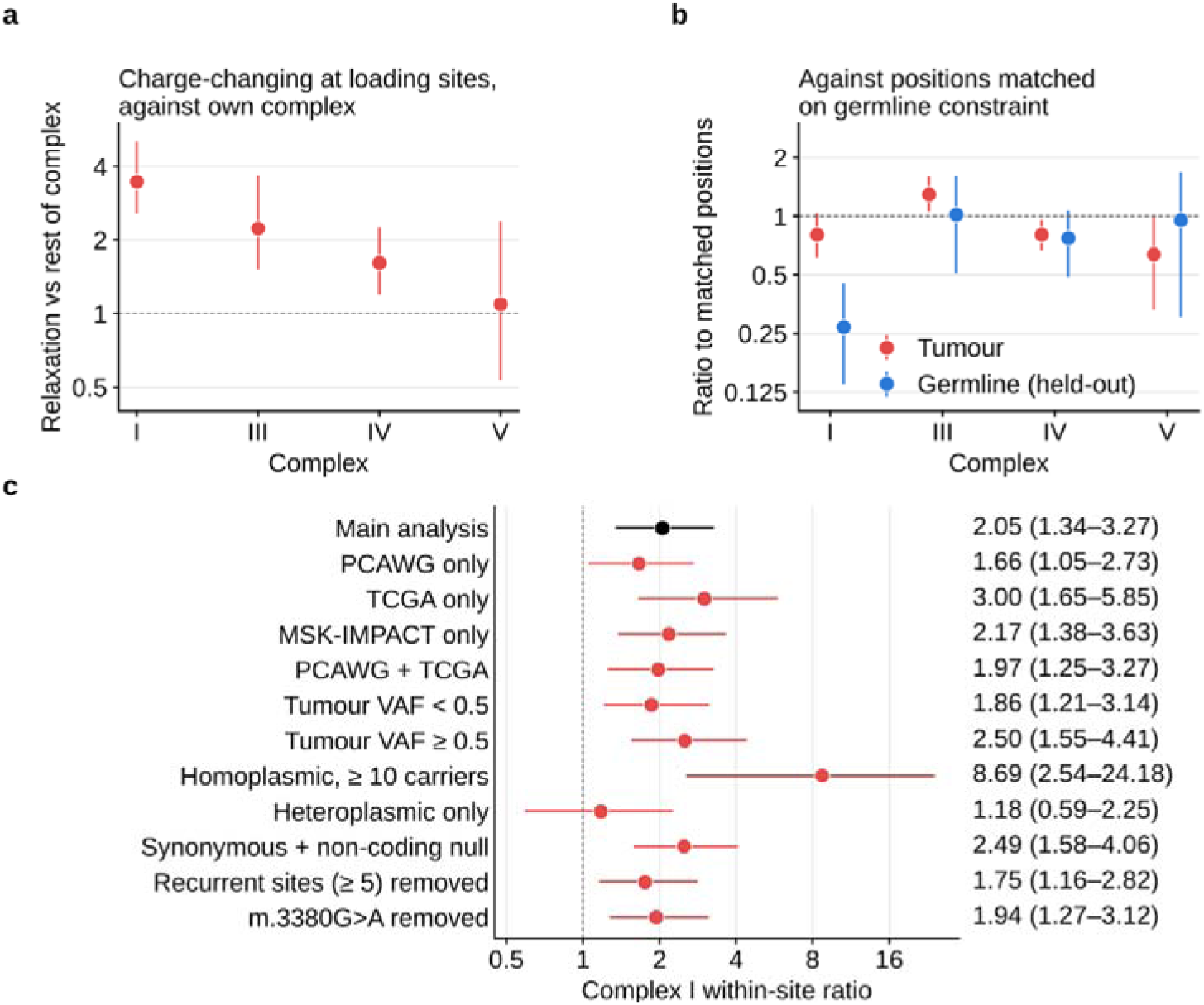
Class-level comparisons track starting constraint, and the within-site contrast is robust. a, Relaxation of charge-changing substitutions at loading sites measured against the rest of their own complex, the design of earlier class-level analyses. The apparent ordering across the chain (complex I 3.46, III 2.23, IV 1.62, V 1.10) reflects starting constraint. b, The same class compared with background positions matched on germline constraint, selected on one random half of population variants. The tumour ratio is 0.80 (0.61–1.03) in complex I, 1.29 (1.05–1.61) in III, 0.80 (0.67–0.96) in IV and 0.64 (0.33–1.01) in V, with no ordering. The germline ratio on the held-out half (blue) checks the match: it is close to 1 except in complex I (0.27, 0.14–0.45), where no background position is as constrained as the class. c, The complex I within-site contrast under each analysis choice applied to complex V, and with the 46 calls of m.3380G>A (ND1 R25Q) removed (1.94, 1.27–3.12). Error bars show 95% percentile intervals; log scales.

**Extended Data Fig. 8.**
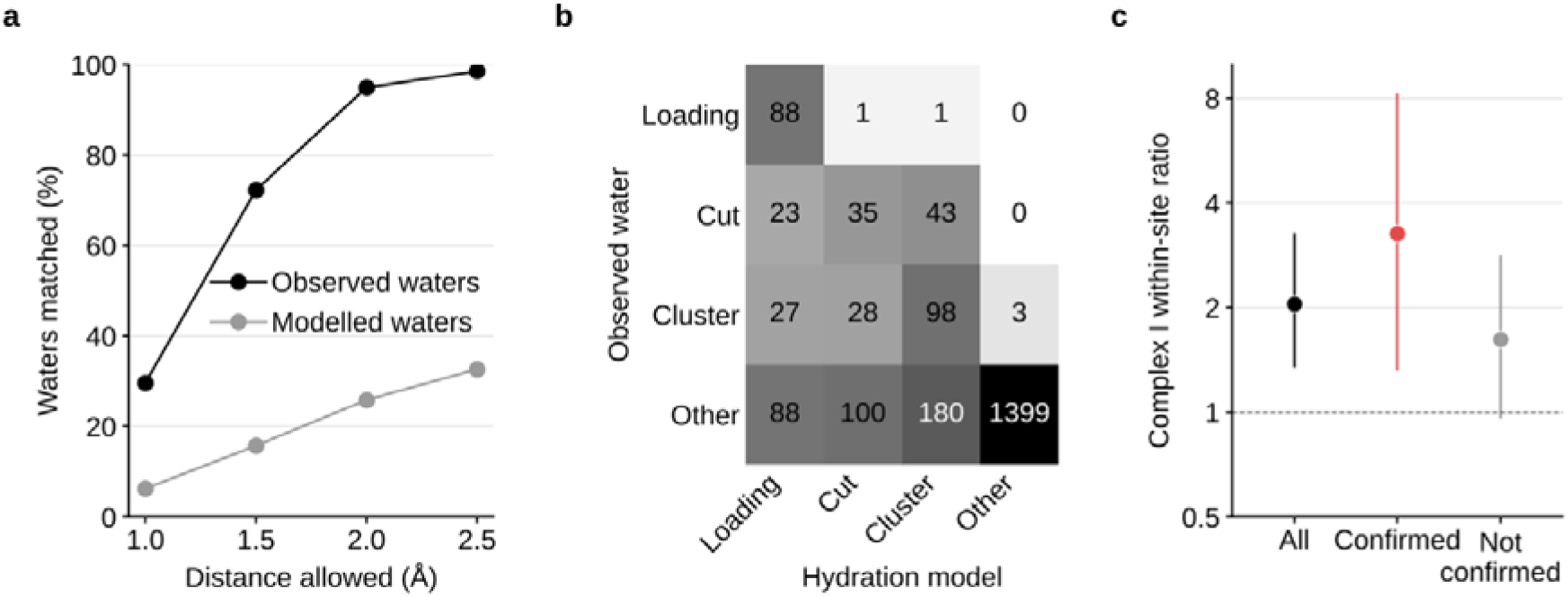
The hydration model checked against a structure that resolves its water. The proton-wire graph was built twice on mouse complex I (PDB 8OM1, 2.39 Å) with the parameters of the main analysis: once from waters placed by the hydration model and once from the buried waters the structure reports. a, Percentage of observed waters with a modelled water within the given distance, and of modelled waters with an observed one. At 2.0 Å these are 94.9% and 25.8%. b, Residue classes from the modelled water (columns) against those from the observed water (rows). The model recovers 88 of the 90 observed loading sites and adds 138. c, Complex I within-site charge contrast at all human loading sites (2.05, 1.34–3.27), at the 74 sites confirmed by observed water (3.27, 1.32–8.27) and at the rest (1.62, 0.96–2.83). Confirmed sites were carried to human residues by Needleman– Wunsch alignment. Error bars show 95% percentile intervals; log scale in c.

**Extended Data Fig. 9.**
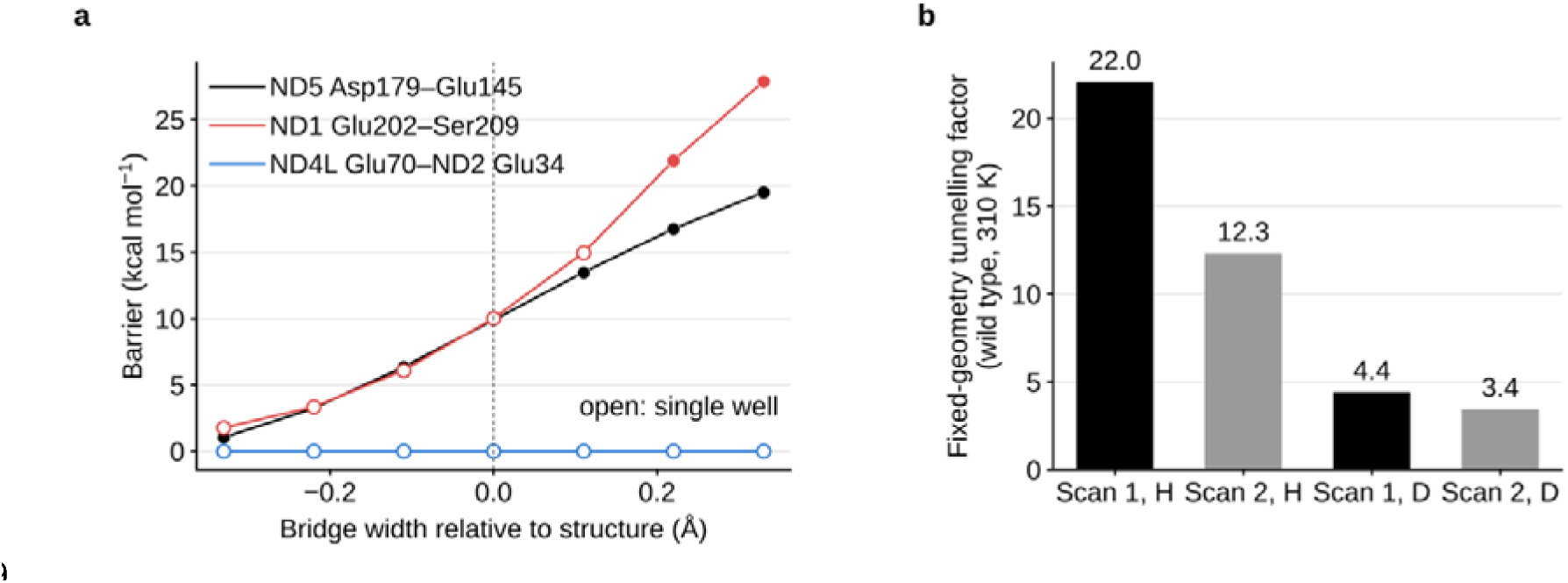
Barrier against bridge width at three loading sites, and the fixed-geometry nuclear treatment. a, Transfer barrier against the width of the bridge relative to the structure. The sites are the MT-ND5 Asp179–water–Glu145 bridge (2.89 Å), the MT-ND1 Glu202–water–Ser209 bridge (2.66 Å; the proton is handed to a hydroxyl) and the Glu70– water–Glu34 bridge between MT-ND4L and MT-ND2 (3.38 Å). Each cluster holds only the three groups of its bridge; for MT-ND5 this is the cluster without Arg176 (the Arg176His system of Fig. 5), so its barriers lie slightly below the wild-type values of Fig. 5d. Open symbols mark scans with a single well. The ND1 gradient is 40 kcal mol□¹ Å□¹, and its curve nearly coincides with that of ND5. The ND4L–ND2 step is downhill at every width sampled. b, Thermally averaged tunnelling factors for the proton (H) and deuteron (D) at 310 K, computed in the fixed-geometry potential from two scans of the same potential. Their disagreement (22.0 against 12.3 for the proton) is one reason these factors are not reported as properties of the enzyme.

**Extended Data Fig. 10.**
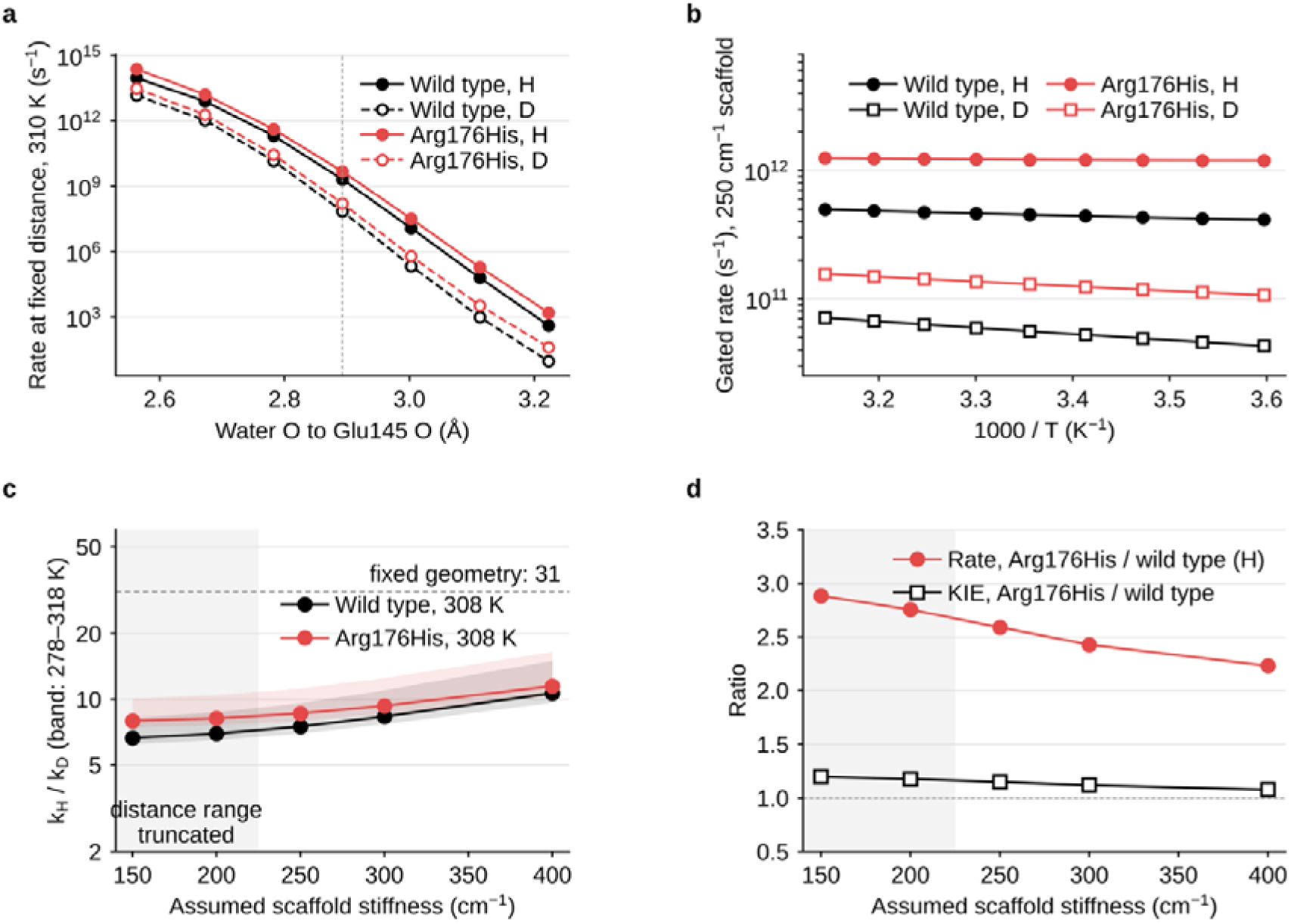
Gated proton tunnelling at the Glu145 bridge of MT-ND5. a, Rate of proton (H, filled) and deuteron (D, open) transfer at each fixed water O–Glu145 O distance at 310 K, for wild type (black) and the Arg176 charge-removal model (red). Rates were computed on the density-functional surface of Fig. 5c by semiclassical transmission from the vibrational levels of the reactant well. The dotted line marks the structural distance of 2.89 Å, where the fixed-geometry isotope effect is 30 at 310 K (31 at the 308 K used elsewhere). b, Arrhenius plot of the gated rates from 278 to 318 K, averaged over a thermal distribution of distances for a scaffold stiffness of 250 cm□¹. c, Kinetic isotope effect k_H_/k_D_ at 308 K against assumed scaffold stiffness. Bands span 278–318 K, and the dashed line marks the fixed-geometry value. d, Ratio of the Arg176 charge-removal model to wild type in the proton rate (red) and in the isotope effect (black); the dashed line marks no change. Grey regions in c and d mark stiffnesses at which the distance distribution extends past the sampled range (13% of the weight at the boundary at 150 cm□¹ and 3% at 200 cm□¹), where values are less reliable. Absolute rates depend on the cluster model and are shown for comparison only. Values are in Supplementary Table 1, sheet S2k.

**Supplementary Fig. 1.**
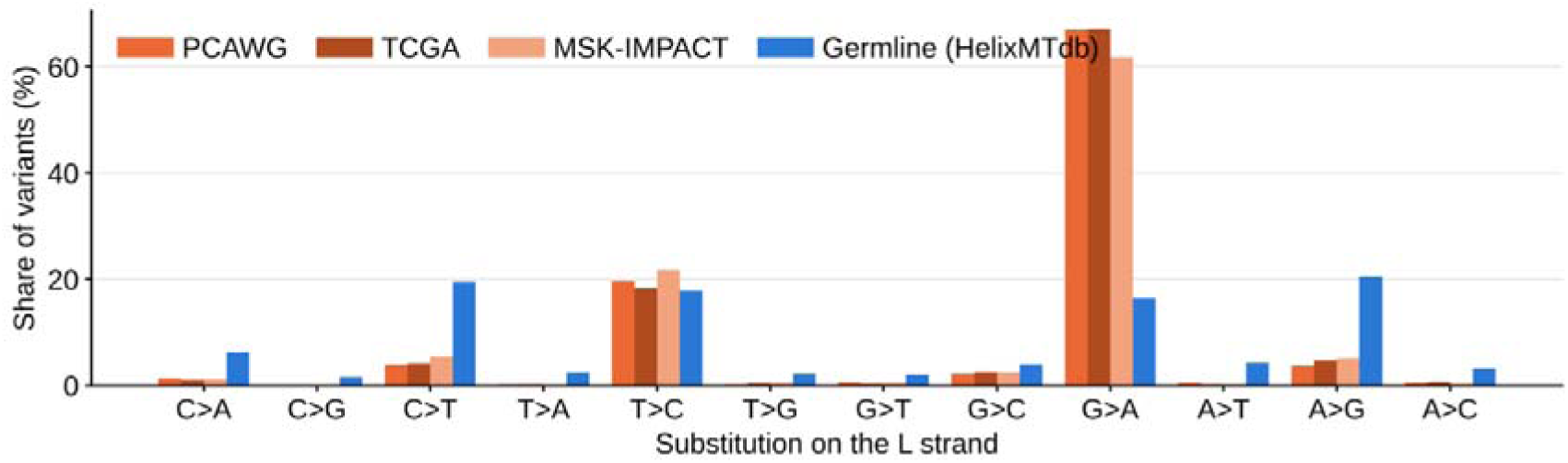
Substitution spectra of the four datasets. Shown is the share of single-nucleotide variants in each of the twelve L-strand substitution types in PCAWG, TCGA, MSK-IMPACT and HelixMTdb. Each dataset’s own 192-class strand-aware trinucleotide null model was estimated from these spectra, using all variants between rCRS positions 577 and 16,023, coding and non-coding.

**Supplementary Fig. 2.**
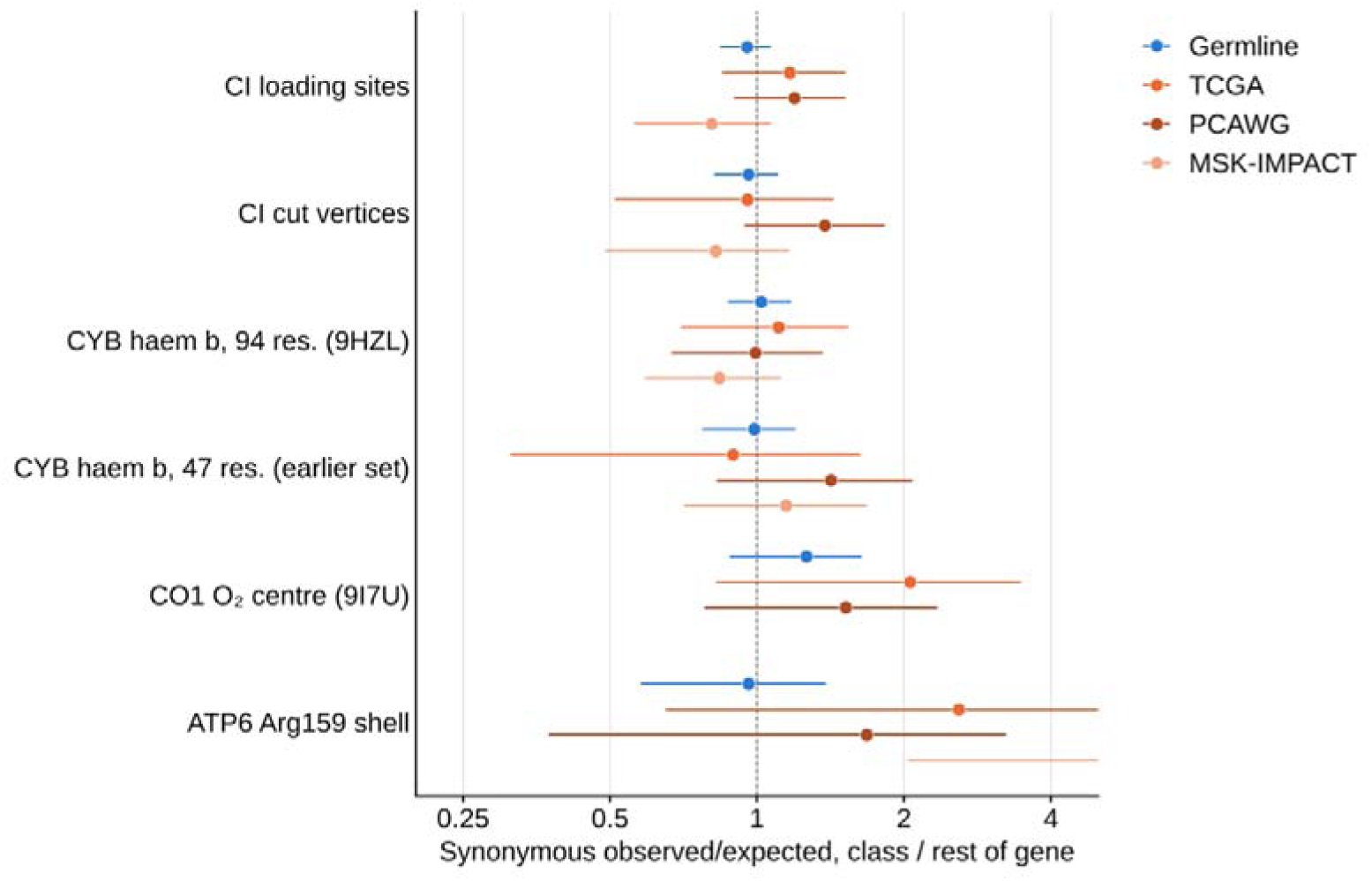
Synonymous calibration of the null model in every structural class. For each class, synonymous observed/expected is shown relative to the same ratio for the rest of the gene (or genes), in the germline and in each tumour cohort. A value of 1 means the null predicts mutational opportunity in the class correctly. The classes are complex I loading sites and cut vertices (PDB 9I4I), the MT-CYB haem b environment (94 residues in PDB 9HZL; and 47 residues as defined in our earlier complex III analysis), the MT-CO1 oxygen-reducing centre (PDB 9I7U) and the MT-ATP6 Arg159 shell (PDB 8H9F). Synonymous variants tend to be in excess at the oxygen-reducing centre in the germline (1.26, 95% CI 0.88–1.64), so the missense depletion reported there is conservative. They are also in excess at the Arg159 shell in tumours (2.24, 1.07–3.61), which would inflate the shell’s apparent tumour constraint. No point is drawn where no synonymous variant was observed. Error bars show 95% percentile intervals from the shared bootstrap resamples; log scale.

**Supplementary Fig. 3.**
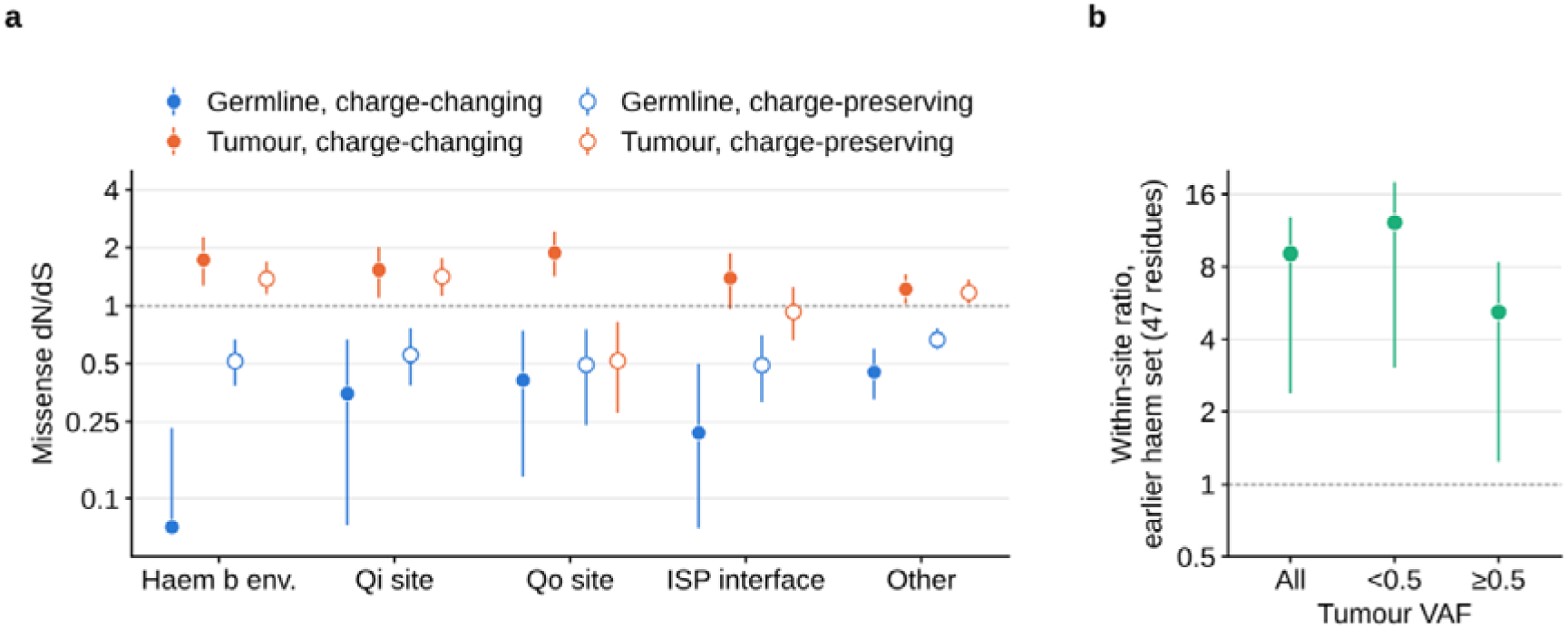
Complex III structural classes as defined previously. a, Missense dN/dS of MT-CYB residues in each structural class of our earlier complex III analysis (haem b environment, Qi site, Qo site and bL leak path, Rieske interface, other; PDB 9CG3). Values are shown for charge-changing (filled) and charge-preserving (open) substitutions, in the germline and pooled tumours, under the unified pipeline. b, Within-site ratio (charge-changing over charge-preserving fold change) for the 47-residue haem b environment of that definition: 9.07 (2.39–12.87) overall, 12.2 (3.05–18.0) below a variant allele fraction (VAF) of 0.5 and 5.20 (1.23–8.38) above. Error bars show 95% percentile intervals; log scales.

**Supplementary Fig. 4.**
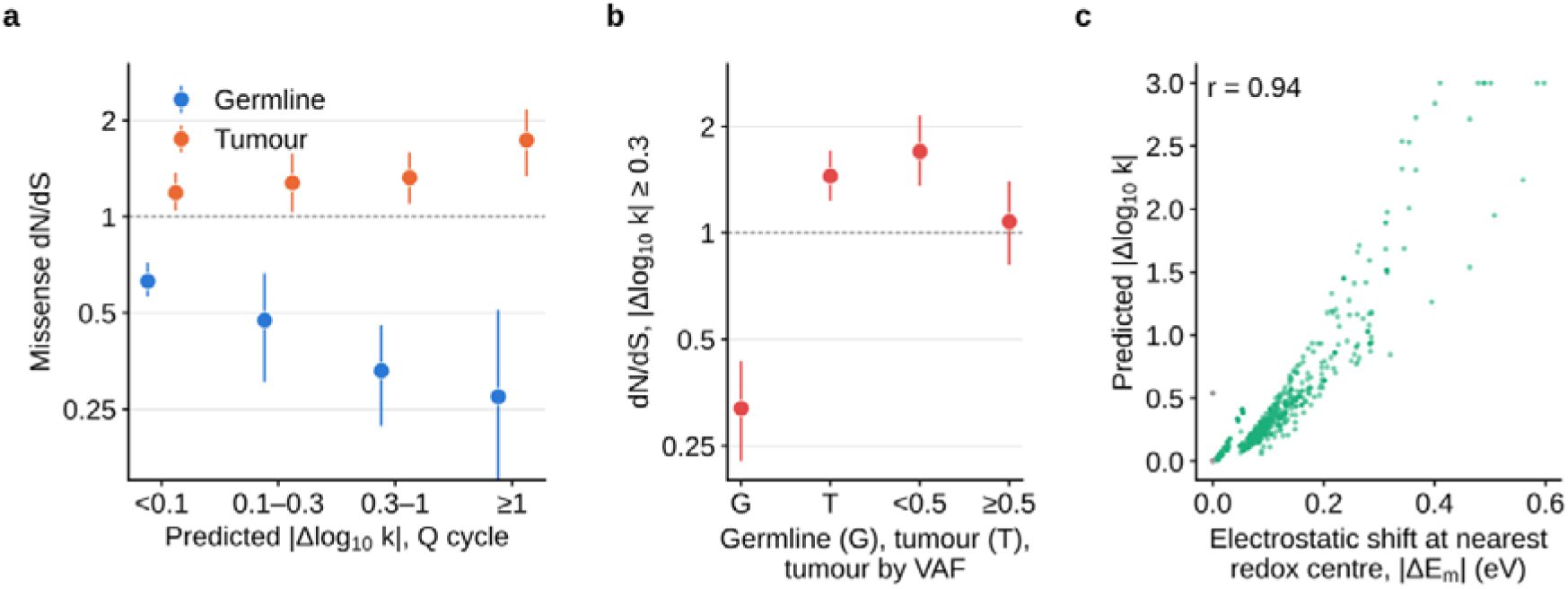
Predicted electron-transfer perturbation and its collinearity with electrostatics. For every possible MT-CYB missense substitution, the predicted change in the three Q-cycle electron-transfer rates was computed from Pathways electronic couplings and the non-adiabatic Marcus expression (PDB 9CG3; effective dielectric constant 8), as in our earlier complex III analysis. a, Missense dN/dS in the germline and pooled tumours across bins of the predicted change, |Δlog□□ k|. b, dN/dS of substitutions with |Δlog□□ k≥ 0.3 in the germline (0.32, 0.23–0.43), in all tumours (1.45, 1.23–1.71) and below (1.70) or above (1.08) a VAF of 0.5. c, Predicted |Δlog□□ k| against the screened Coulomb shift at the nearest redox centre for all possible substitutions (green, charge-changing; grey, charge-preserving). The correlation, r = 0.94, prevents the electron-transfer effect from being separated from the electrostatic one. Error bars show 95% percentile intervals; log scales in a and b.

**Supplementary Fig. 5.**
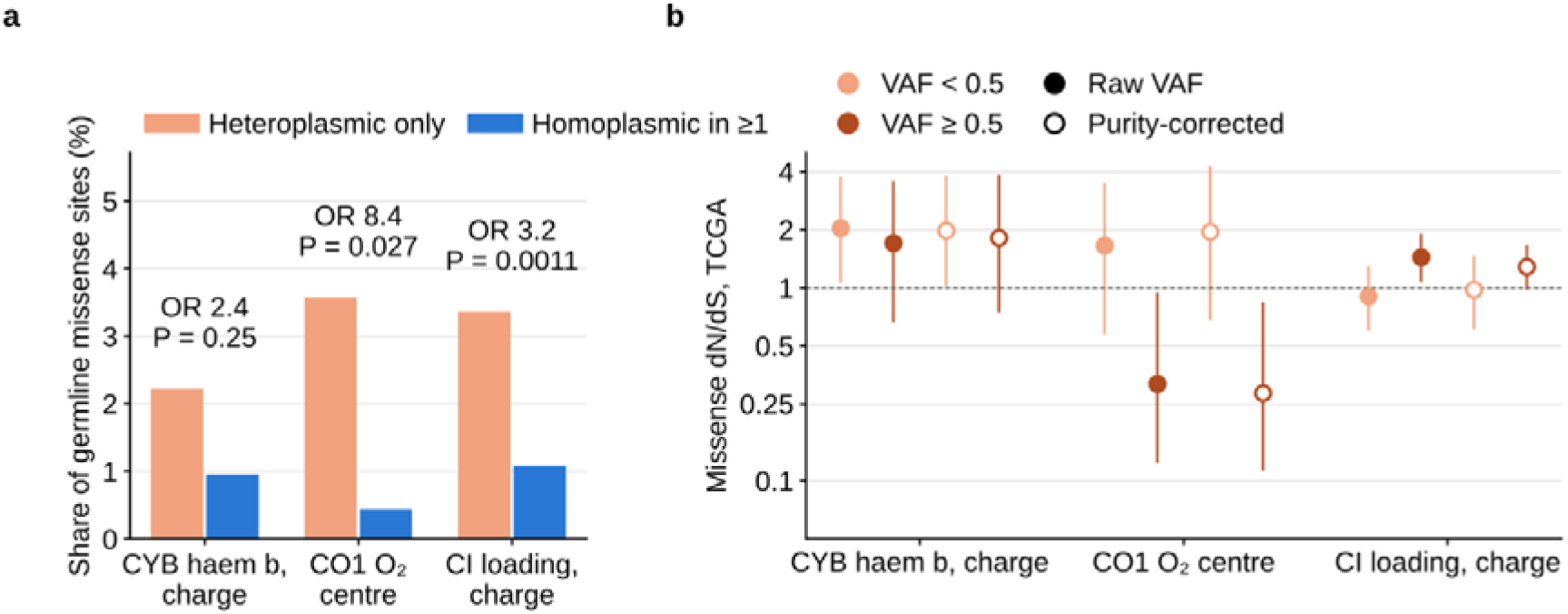
Germline heteroplasmy state and purity-corrected allele fractions. a, Share of HelixMTdb missense variant sites in each class among sites seen only in the heteroplasmic state and among sites homoplasmic in at least one carrier. The classes are charge-changing substitutions in the MT-CYB haem b environment, the MT-CO1 oxygen-reducing centre and charge-changing substitutions at complex I loading sites. Odds ratios and P values are from two-sided Fisher’s exact tests. b, Missense dN/dS of the same classes in TCGA tumours, below and above a VAF of 0.5, using the raw VAF (filled) or the VAF divided by the consensus purity estimate and capped at 1 (open). Error bars show 95% percentile intervals; log scale in b.

**Supplementary Fig. 6.**
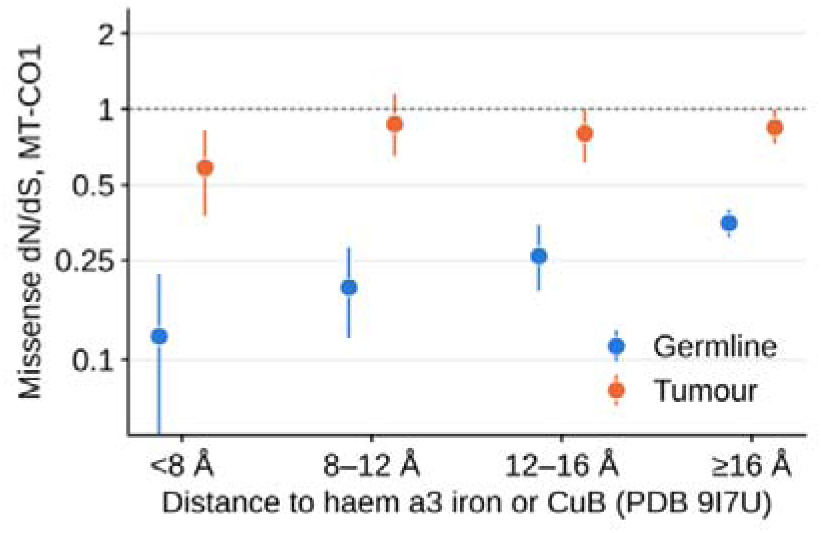
Distance gradient of constraint around the oxygen-reducing centre. Missense dN/dS of MT-CO1 residues binned by the minimum distance of any heavy atom to the haem a3 iron or CuB (PDB 9I7U), in the germline and pooled tumours. Germline dN/dS rises from 0.12 (0.05–0.22) within 8 Å to 0.35 (0.31–0.40) beyond 16 Å. Error bars show 95% percentile intervals; log scale.

**Supplementary Fig. 7.**
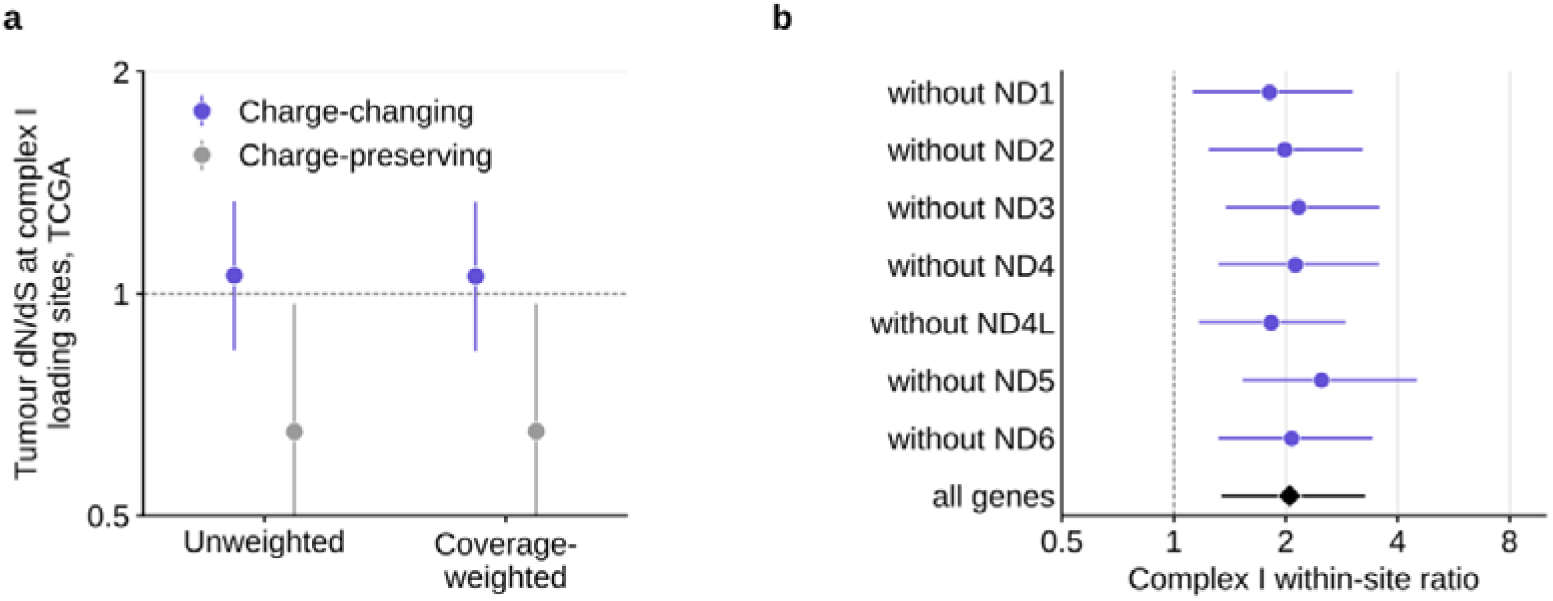
Sequencing coverage and leave-one-gene-out. a, Tumour dN/dS of charge-changing and charge-preserving substitutions at complex I loading sites in TCGA, with expected counts unweighted or weighted by the number of tumours callable at each position. The mean callable fraction is 0.779 at loading-site codons and 0.779 elsewhere in the ND genes. b, Complex I within-site ratio with each ND gene removed in turn from both the class and the synonymous denominator. Error bars show 95% percentile intervals; log scales.

**Supplementary Fig. 8.**
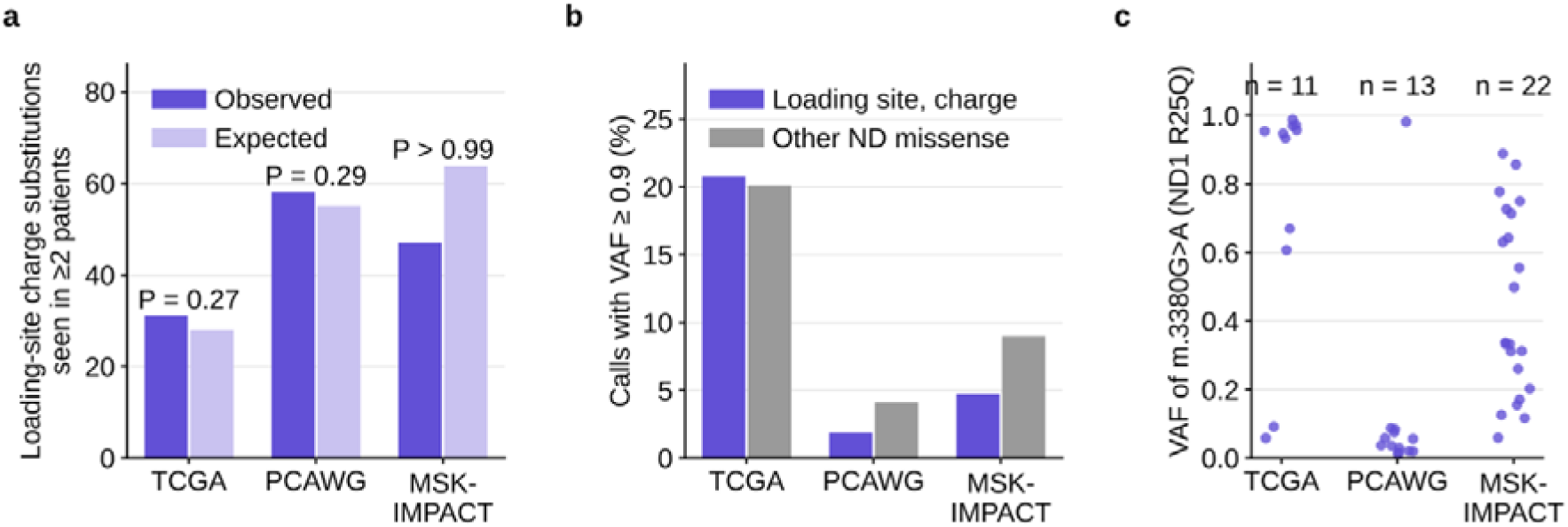
Recurrence and clonality of complex I loading-site substitutions. a, Number of charge-changing loading-site substitutions seen in two or more patients, observed and expected. The expectation comes from 2,000 multinomial draws of each cohort’s ND missense calls over all possible ND missense substitutions, weighted by the tumour spectrum. P is the fraction of draws with at least the observed number. b, Percentage of calls with a VAF of at least 0.9, for charge-changing loading-site substitutions and other ND missense substitutions. c, VAF of every call of m.3380G>A (ND1 R25Q), the most recurrent loading-site substitution, by cohort.

**Supplementary Fig. 9.**
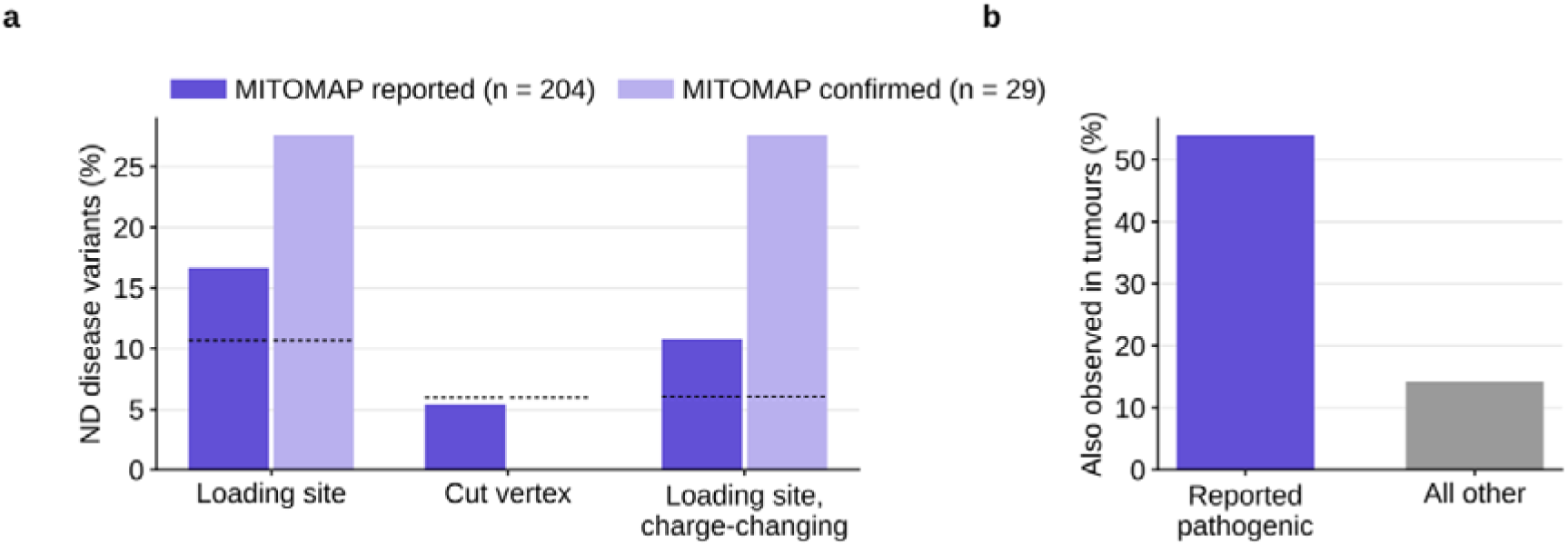
Mitochondrial disease variants on the complex I proton wire. a, Share of ND missense variants in MITOMAP (release of 25 May 2022) at loading sites, at cut vertices and among charge-changing substitutions at loading sites. Shown separately are all reported variants (n = 204) and those with confirmed pathogenic status (n = 29). Dashed lines mark the share expected from the germline spectrum over the same residues. Confirmed variants fall in the charge-changing loading-site class in 8 of 29 cases against 6.0% expected (binomial P = 2 × 10□□). No confirmed variant lies at a cut vertex, so that bar is absent. b, Percentage of the reported variants in a (n = 204) observed in at least one tumour (54%), against 14% of all other possible ND missense substitutions.

**Supplementary Fig. 10.**
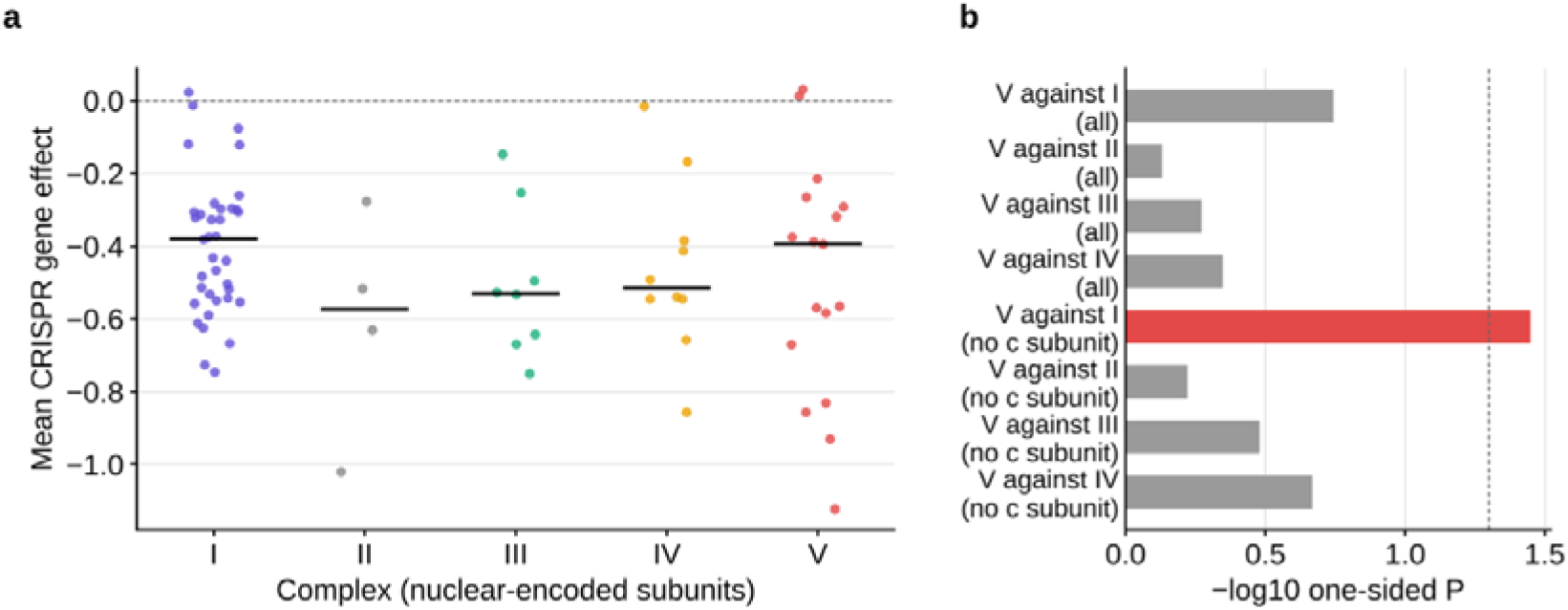
CRISPR essentiality of the respiratory complexes in cancer cell lines. a, Mean CRISPR gene effect (DepMap, 1,361 cell lines) of the nuclear-encoded structural subunits of each complex; lines mark medians. b, One-sided Mann–Whitney P values for complex V being more essential than each other complex, with all subunits or with the three c-subunit paralogues set aside, which CRISPR cannot score. With the paralogues set aside, complex V is more essential than complex I (P = 0.036) but not than complexes II, III or IV. The dashed line marks P = 0.05.

**Supplementary Fig. 11.**
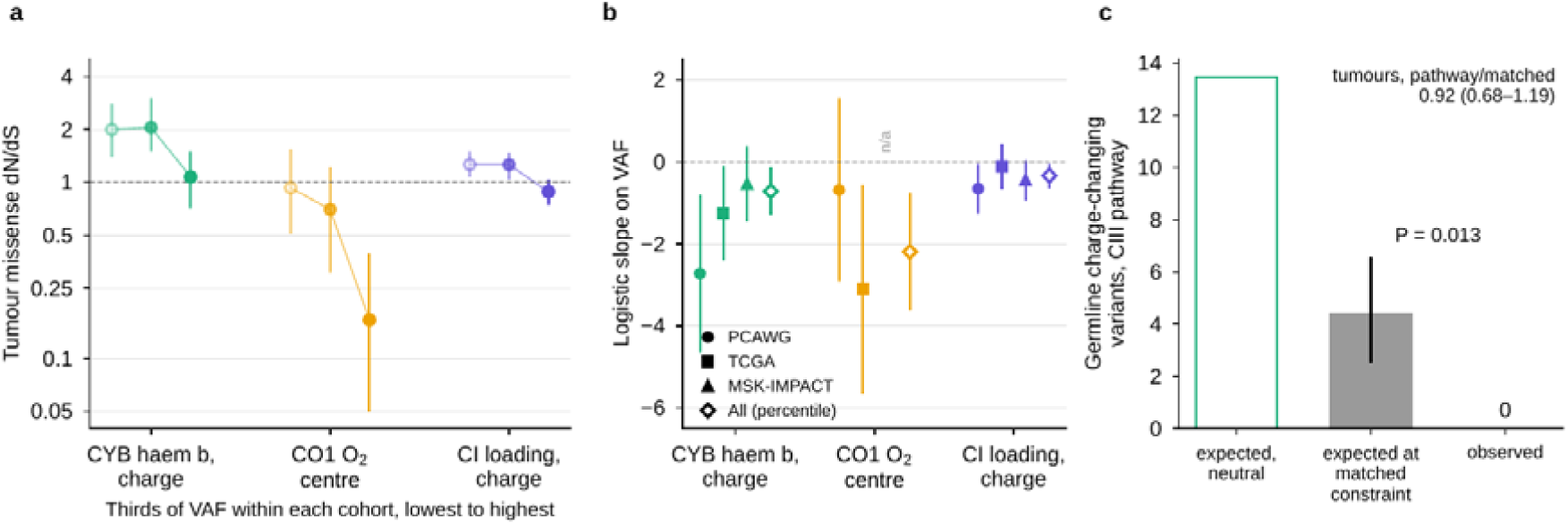
Dependence on VAF within cohorts, and charge changes on the complex III electron-tunnelling pathway against matched residues. a, Tumour missense dN/dS of charge-changing substitutions in the MT-CYB haem b environment, of missense substitutions at the MT-CO1 oxygen-reducing centre and of charge-changing substitutions at complex I loading sites, in thirds of VAF defined within each cohort (lowest to highest, left to right), so that no stratum is dominated by one cohort; rates come from the whole tumour spectrum and observations from the stratum. b, Logistic slope of class membership on VAF among the missense variants of each gene within PCAWG, TCGA and MSK-IMPACT (filled), and with VAF as a within-cohort percentile and cohort as a covariate (open). MSK-IMPACT carries no variant at the oxygen-reducing centre (n/a). c, Germline charge-changing substitutions on the complex III pathway: expected under neutrality, expected if the pathway were as constrained as residues matched on burial and distance to the cofactors (bar, with the interval from the matched dN/dS), and observed. P is the Poisson probability of observing none at the matched constraint. The tumour pathway-to-matched ratio is given above. Error bars show 95% percentile intervals from the shared resamples (a, c) or 95% Wald intervals (b); log scale in a. Values are in Supplementary Table 1, sheets S2o and S2p.

**Supplementary Table 1 Every estimate reported in this study.** One sheet per analysis, with estimates, 95% intervals, P values and the counts they rest on. Sheets S1a–S1al hold the analyses of complex V, the proton-wire graphs and the density-functional calculations. Sheets S2a–S2q hold the analyses of the complex III haem b environment, the complex IV oxygen-reducing centre, heteroplasmy, the phylogeny and the supplementary analyses, including tunnelling and its robustness analyses (S2k–S2n), the dependence on VAF within cohorts (S2o), charge changes on the complex III electron-tunnelling pathway against matched residues (S2p) and the basis-set and dispersion check of the proton-transfer potential (S2q). The first row of each sheet describes its contents.

**Supplementary Table 2 Residue classifications.** The proton-wire classes (loading site, cut vertex, cluster member, reference) of every mtDNA-encoded residue in PDB 9I4I, 9HZL, 9I7U and 8H9F, and the structural regions of MT-ATP6 and MT-ATP8 in PDB 8H9F. Also given are the MT-CYB haem b environment at 6 and 8 Å in PDB 9HZL and 9CG3, with the quinone-site and Rieske-interface exclusions and the categories of our earlier complex III analysis, and the MT-CO1 oxygen-reducing centre in PDB 9I7U and 5B1A. Also given are the residues of each electron-tunnelling pathway with their loss of coupling.

**Supplementary Table 3 Candidate variants for a test of the two classes. mtDNA substitutions in subunit a of ATP synthase on the charge axis (axis A) and the wire-severing axis (axis B).** For each, the table gives the protein and mtDNA change, wire class, number of tumours carrying it, HelixMTdb heteroplasmic and homoplasmic carrier counts, and distance to the Arg159 guanidinium.

**Supplementary Table 4 Recurrent loading-site substitutions and substitutions fixed on the phylogeny.** Charge-changing substitutions at complex I loading sites in tumours, with the number of patients carrying each and their median VAF. Also listed are the 14 ND missense substitutions at loading sites that define branches of PhyloTree build 17, and every call of m.3380G>A with its cohort, VAF and cancer type.

