## Supplementary figures and images for "Evolutionary constraints on mitochondrial bioenergetics are reshaped in cancer"

### Extended data figure 1

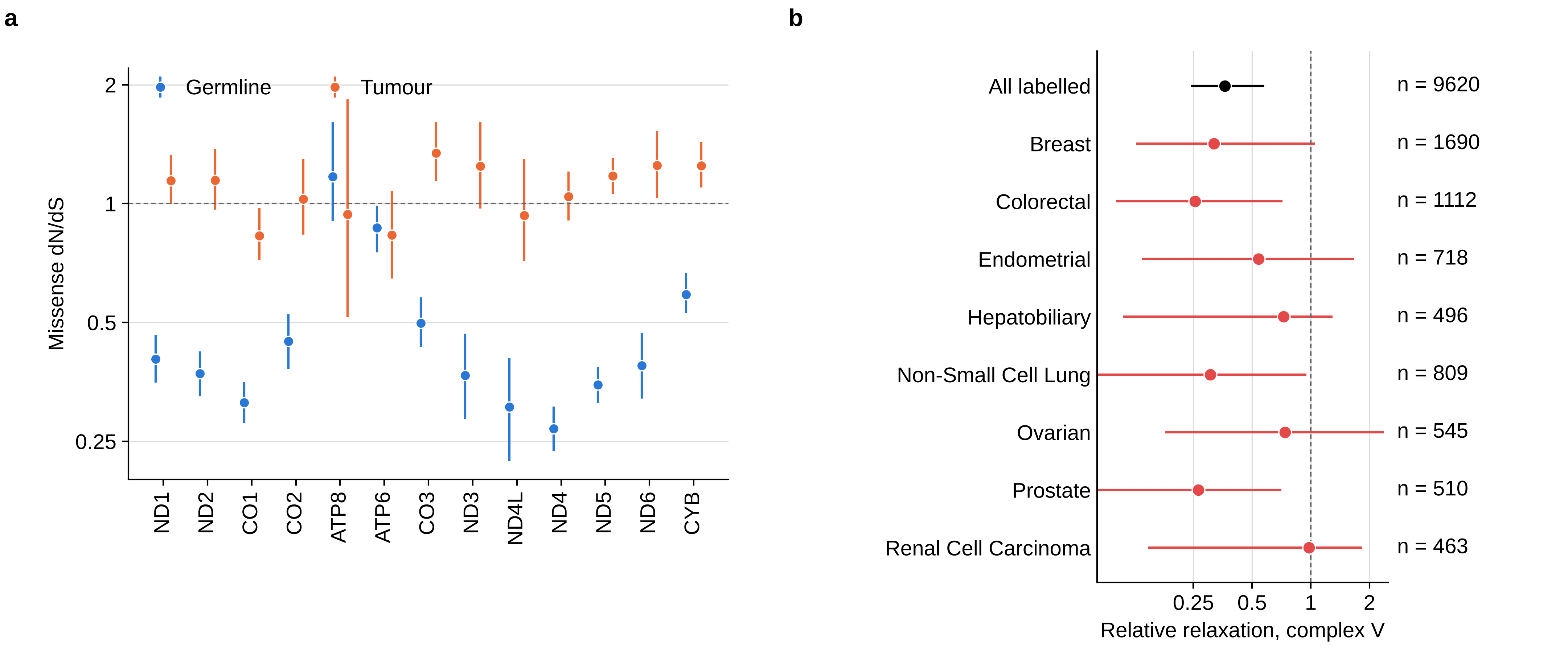

### Extended data figure 2

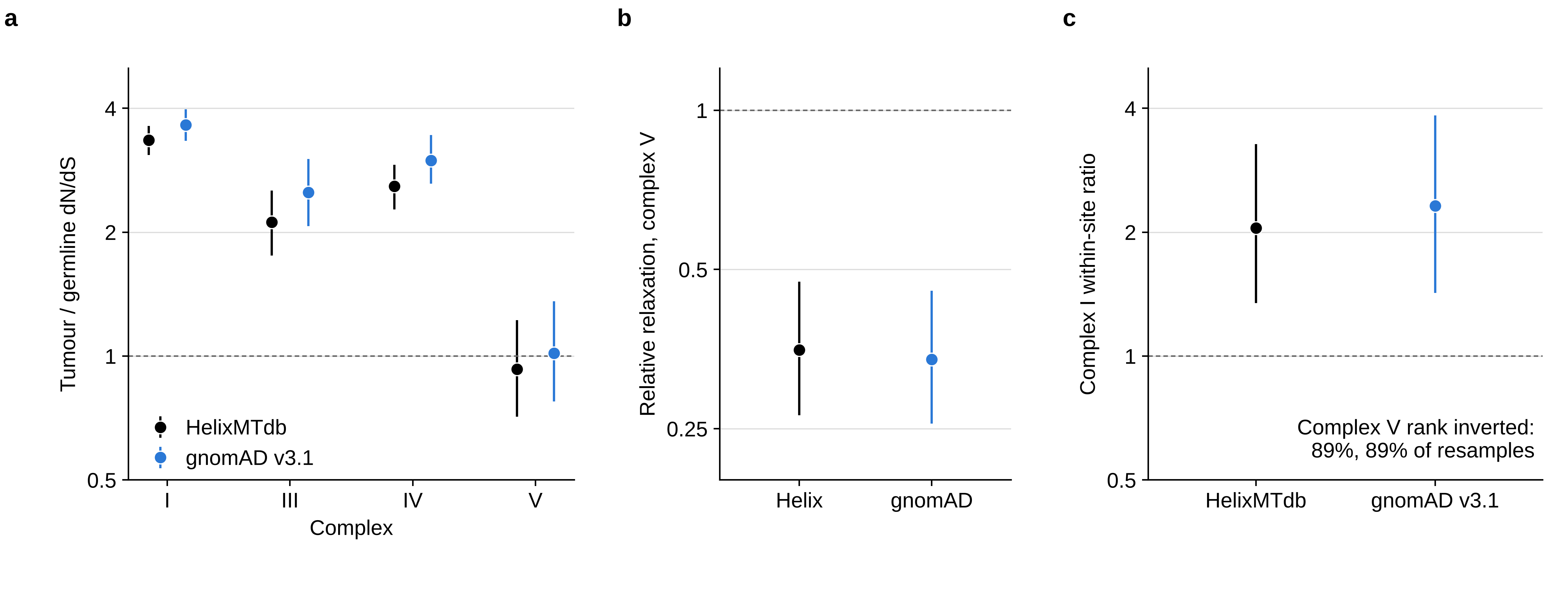

### Extended data figure 3

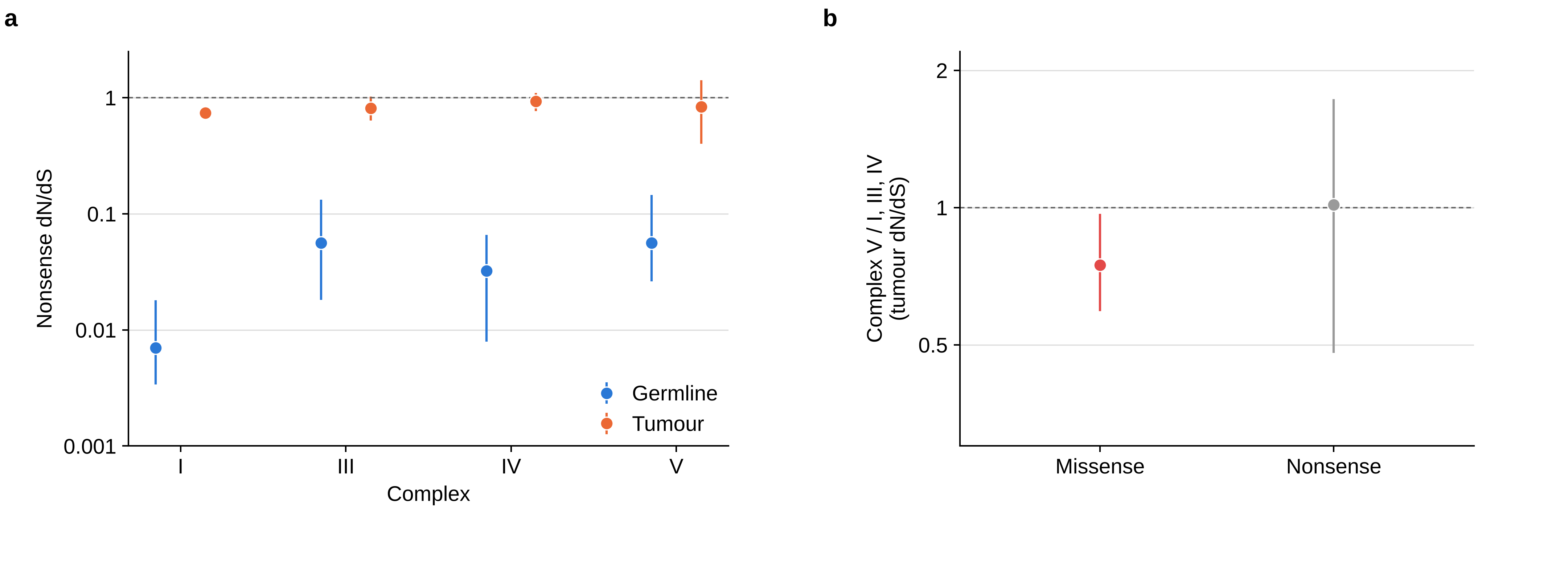

### Extended data figure 4

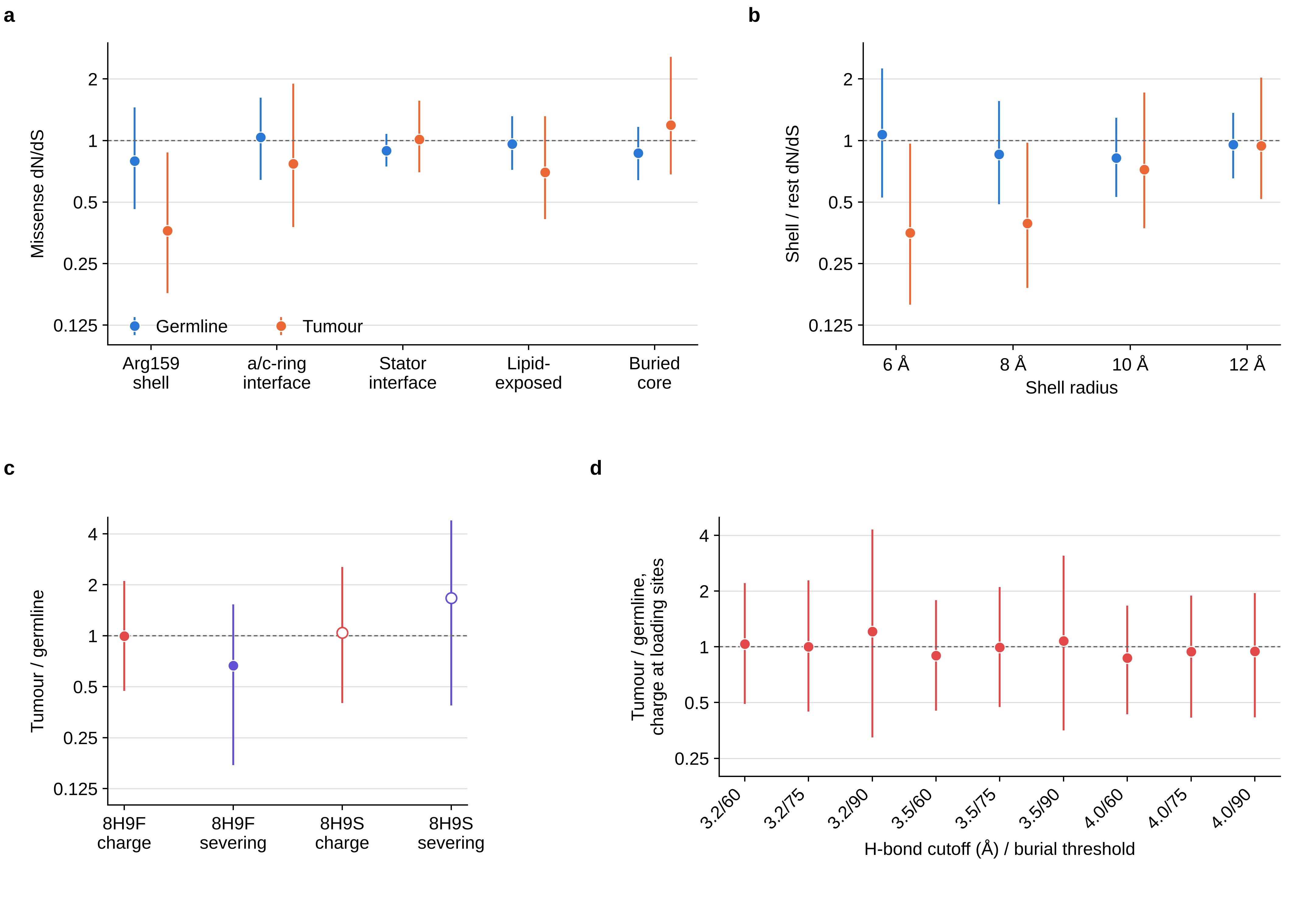

### Extended data figure 5

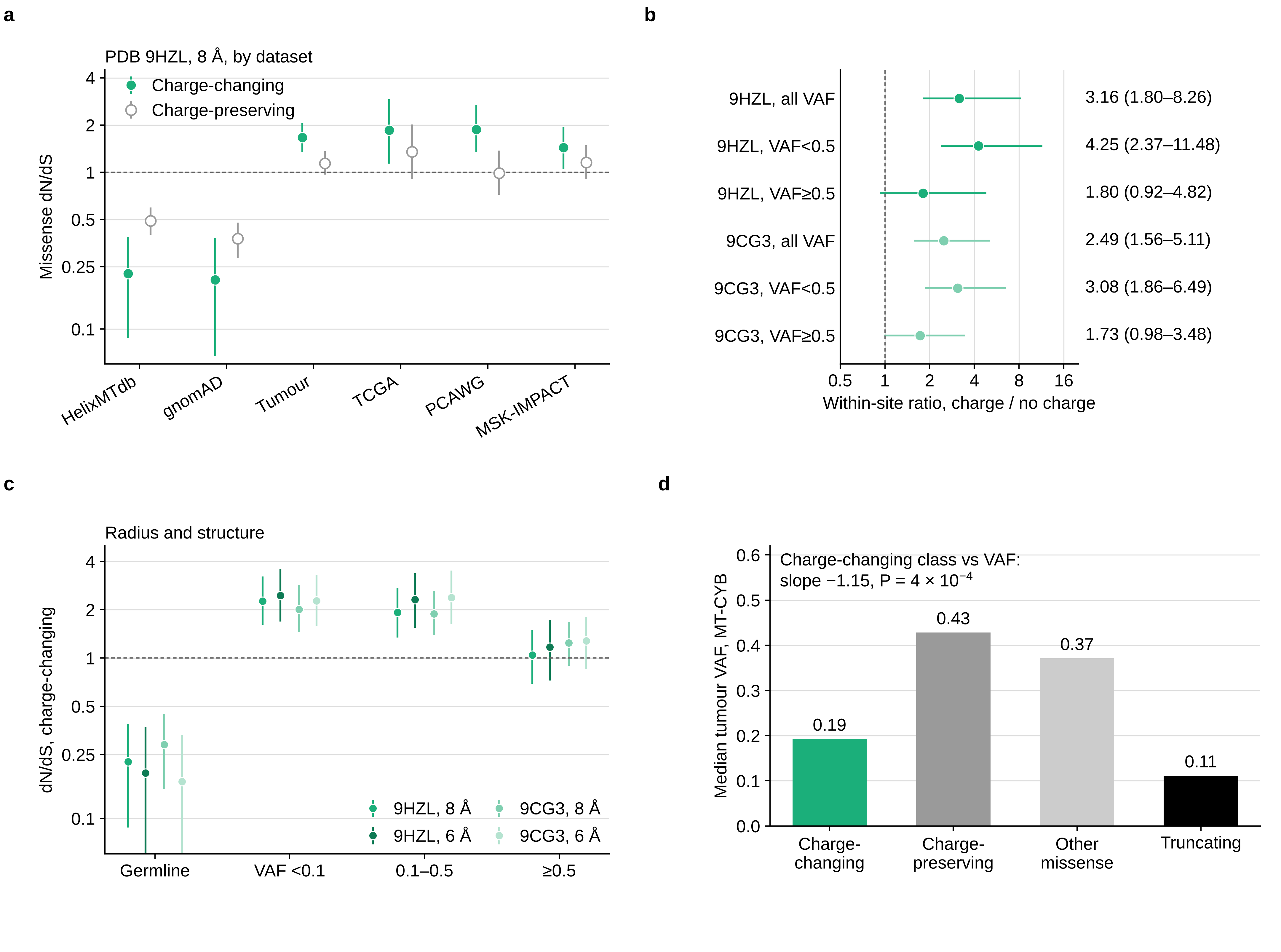

### Extended data figure 6

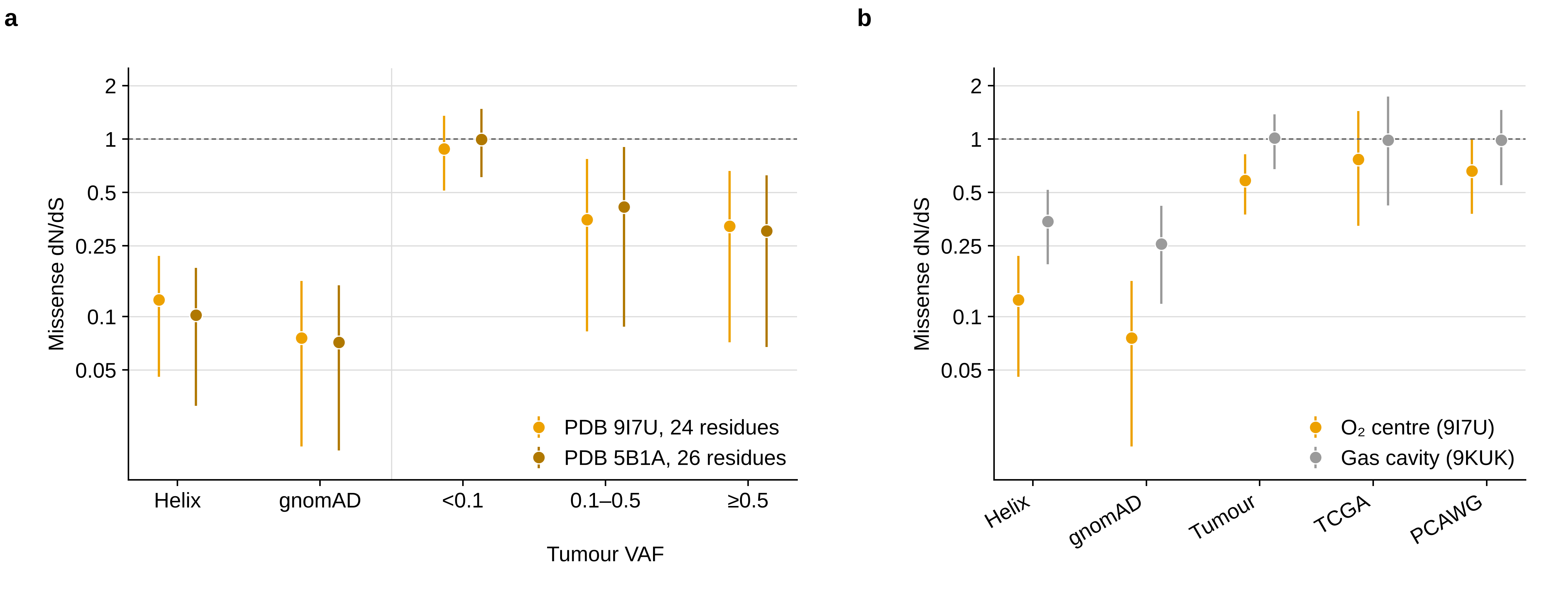

### Extended data figure 7

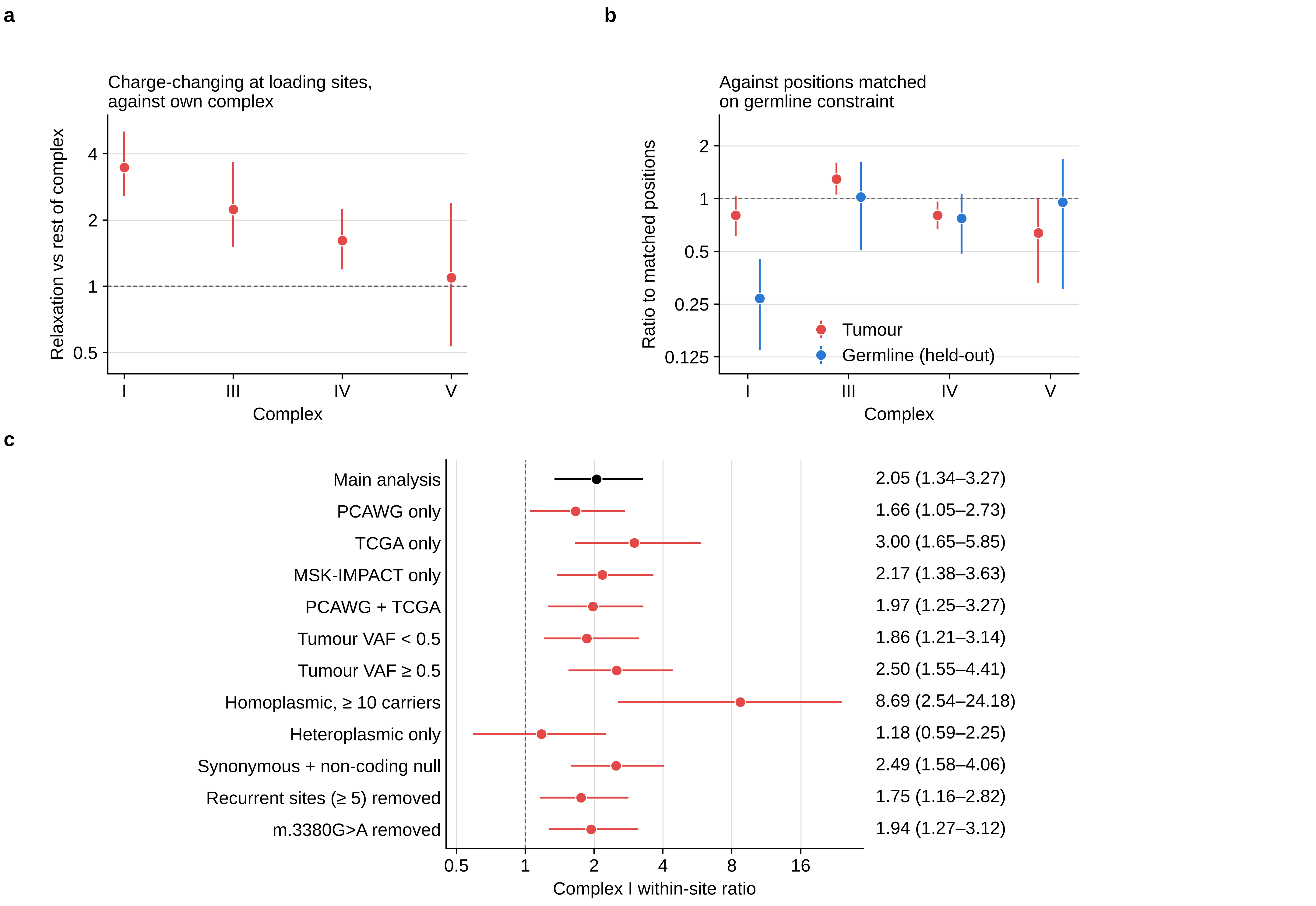

### Extended data figure 8

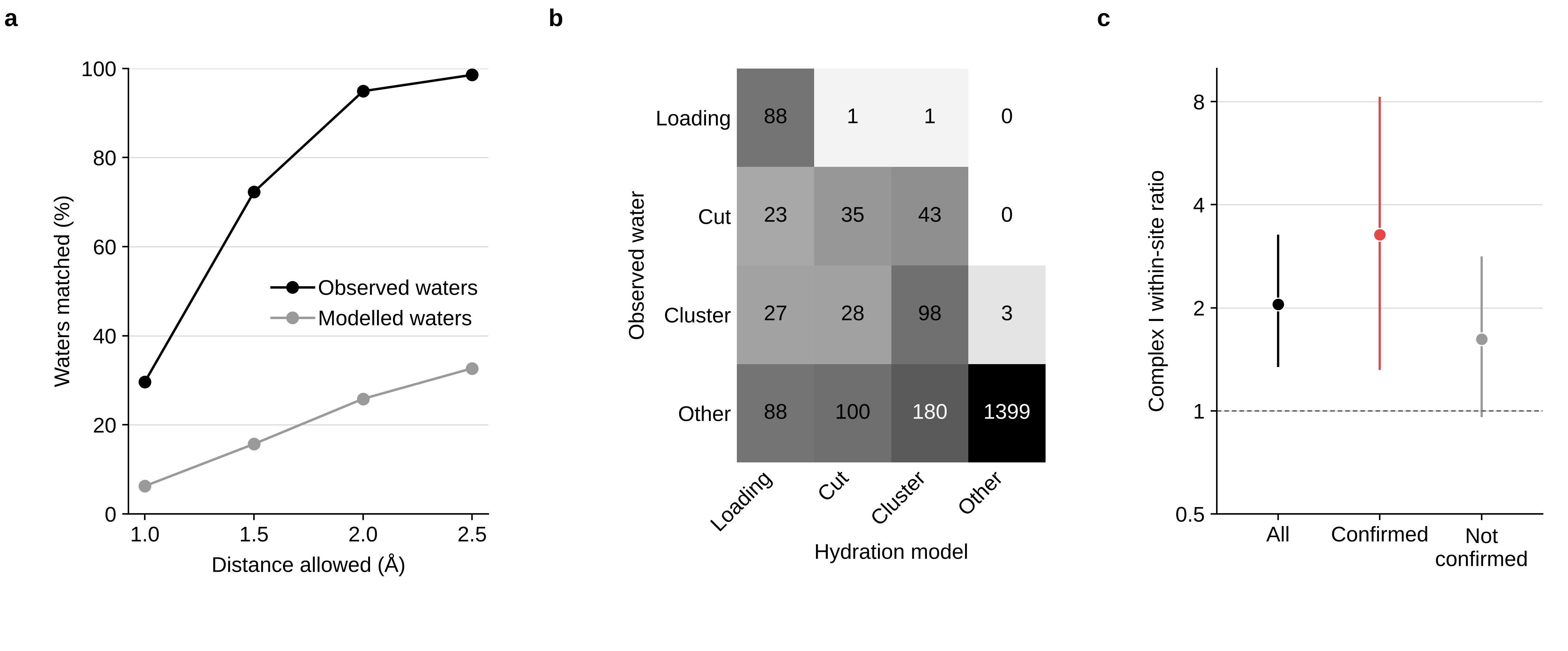

### Extended data figure 9

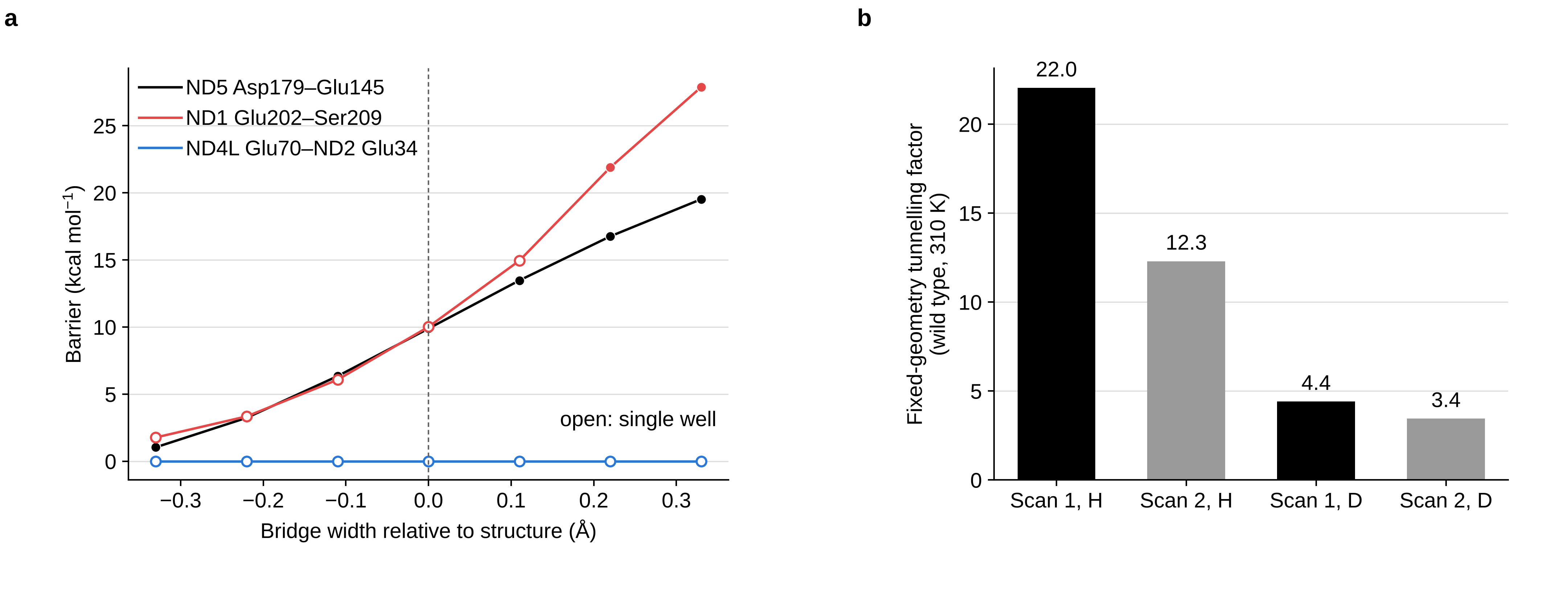

### Extended data figure 10

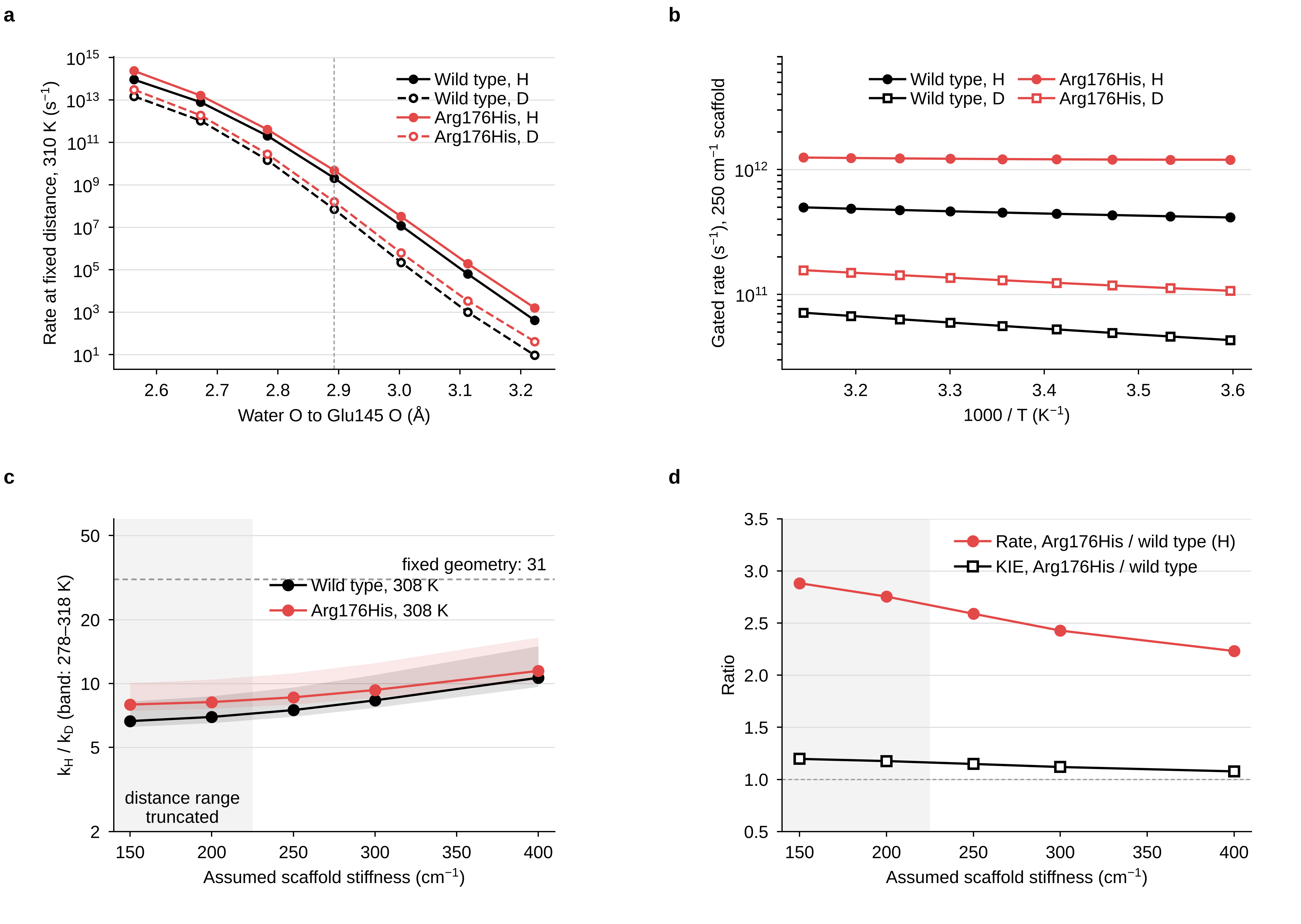

### supple fig1

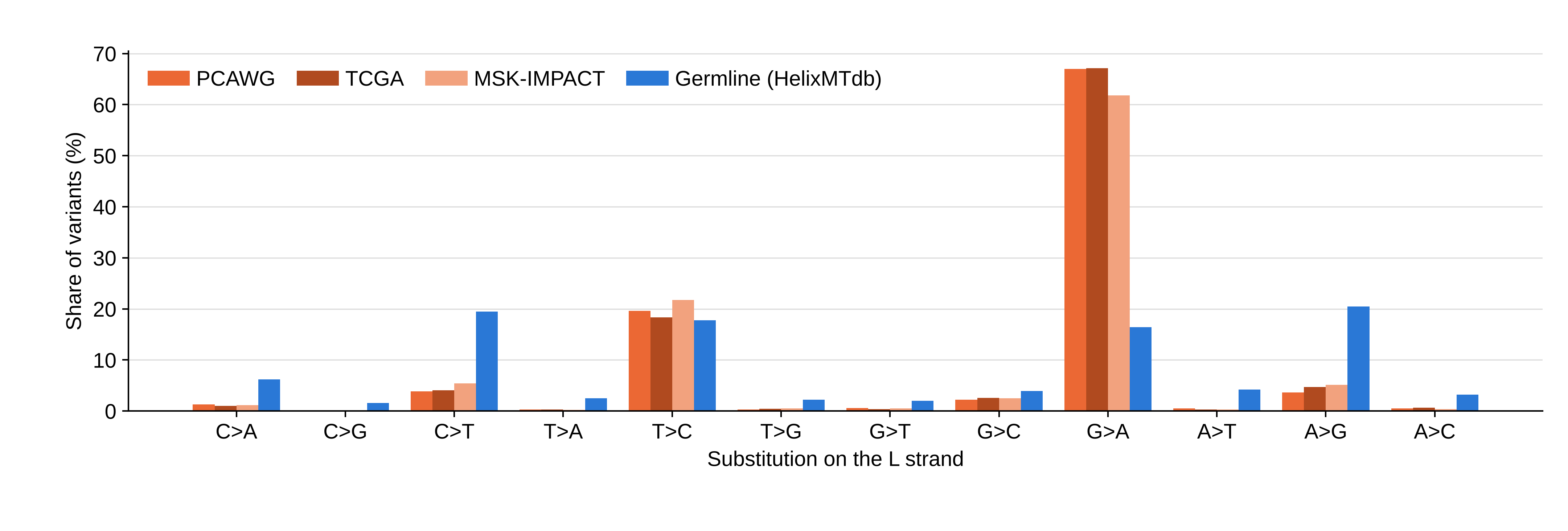

### supple fig2

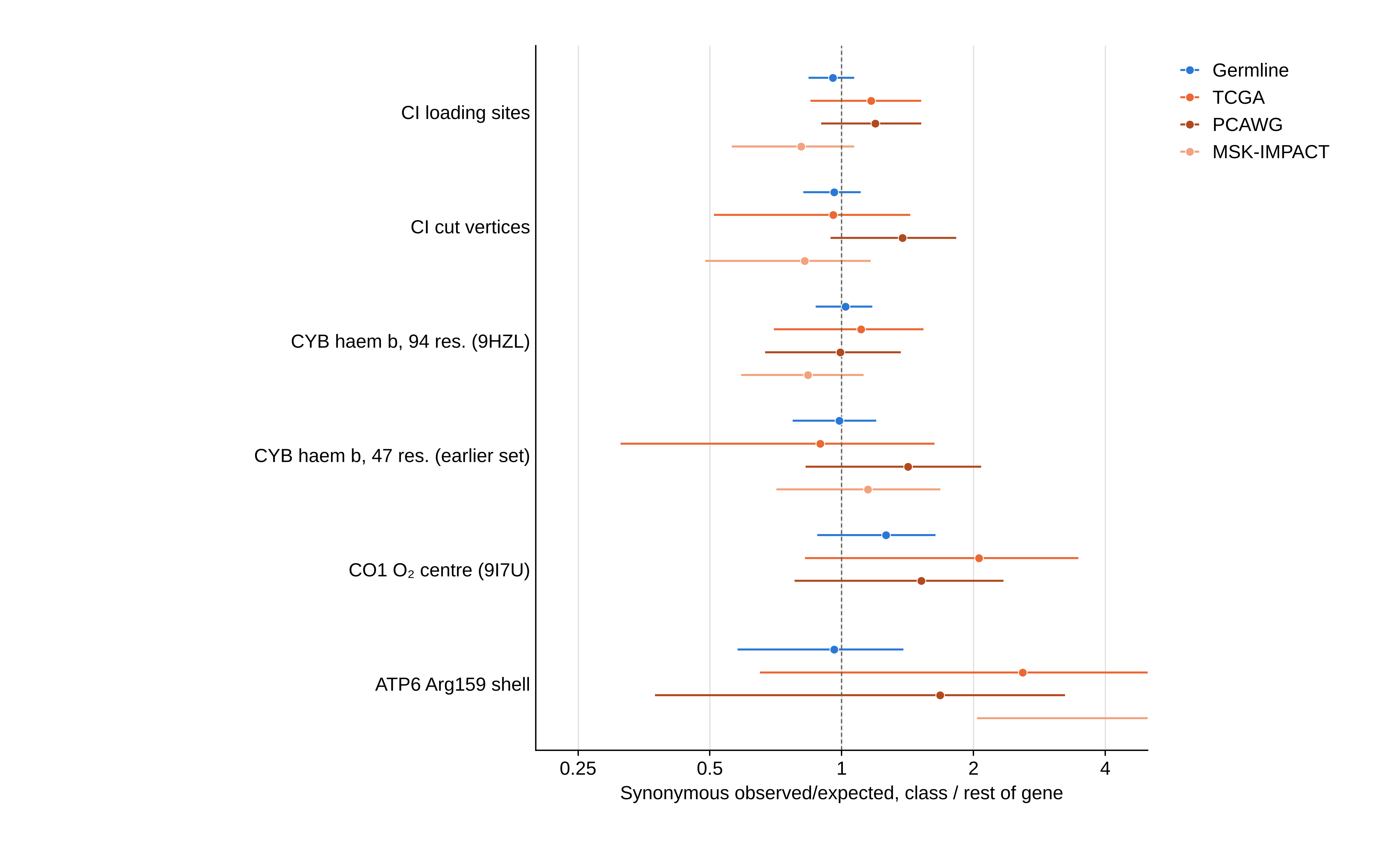

### supple fig3

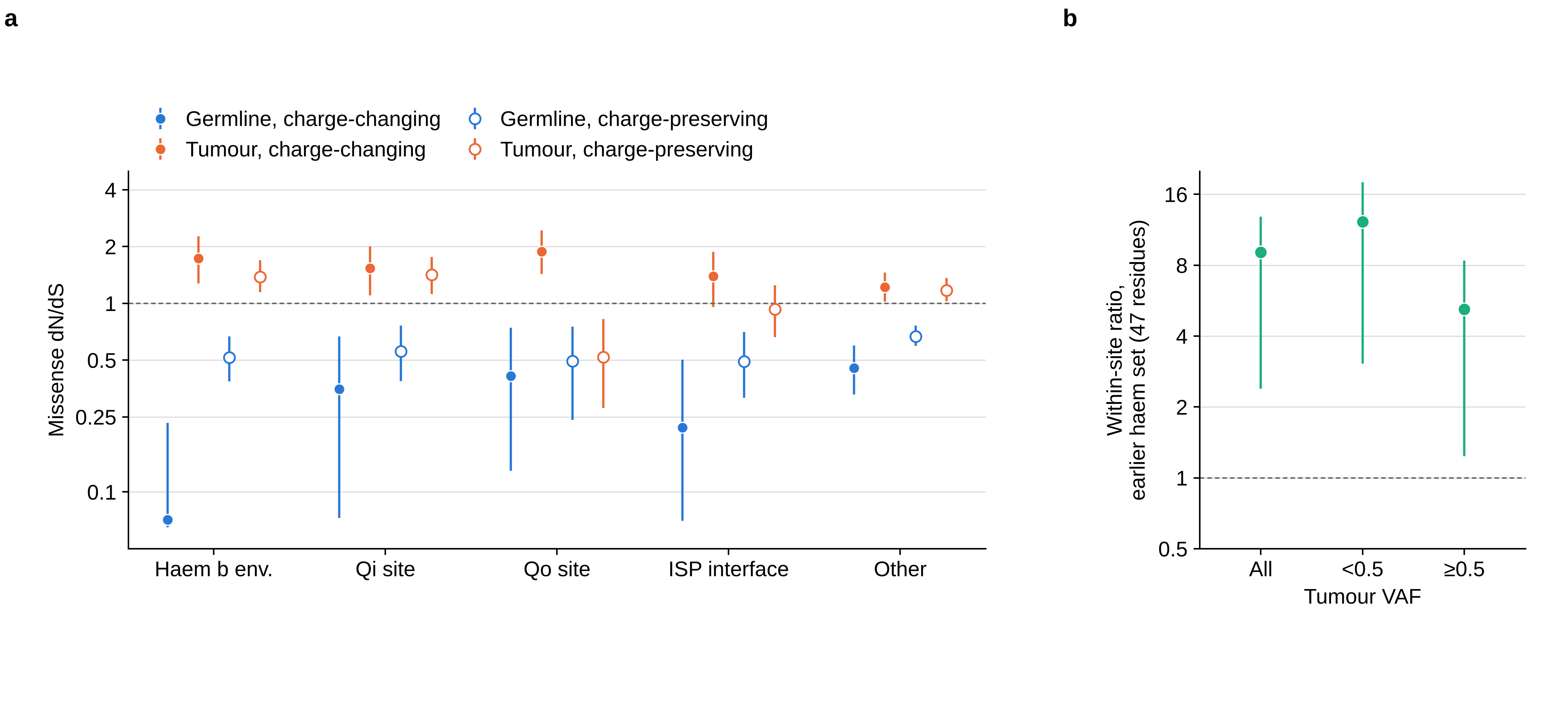

### supple fig4

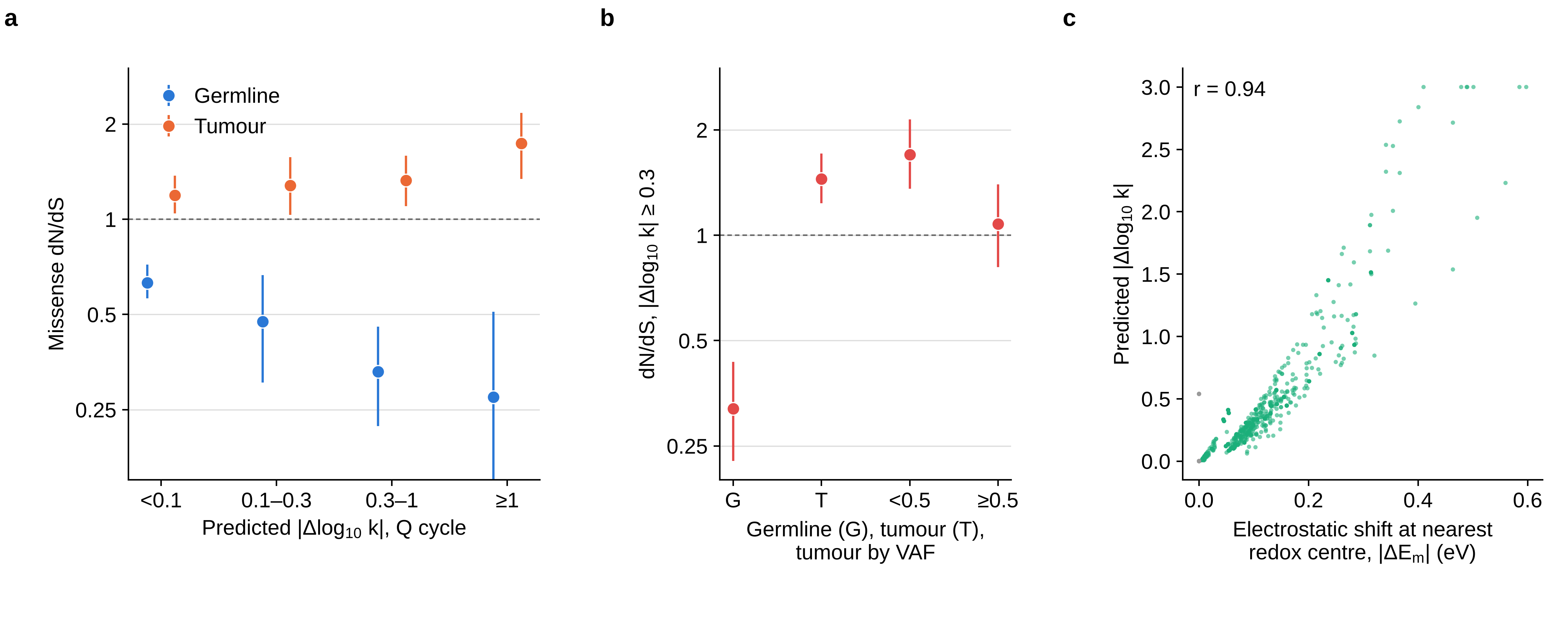

### supple fig5

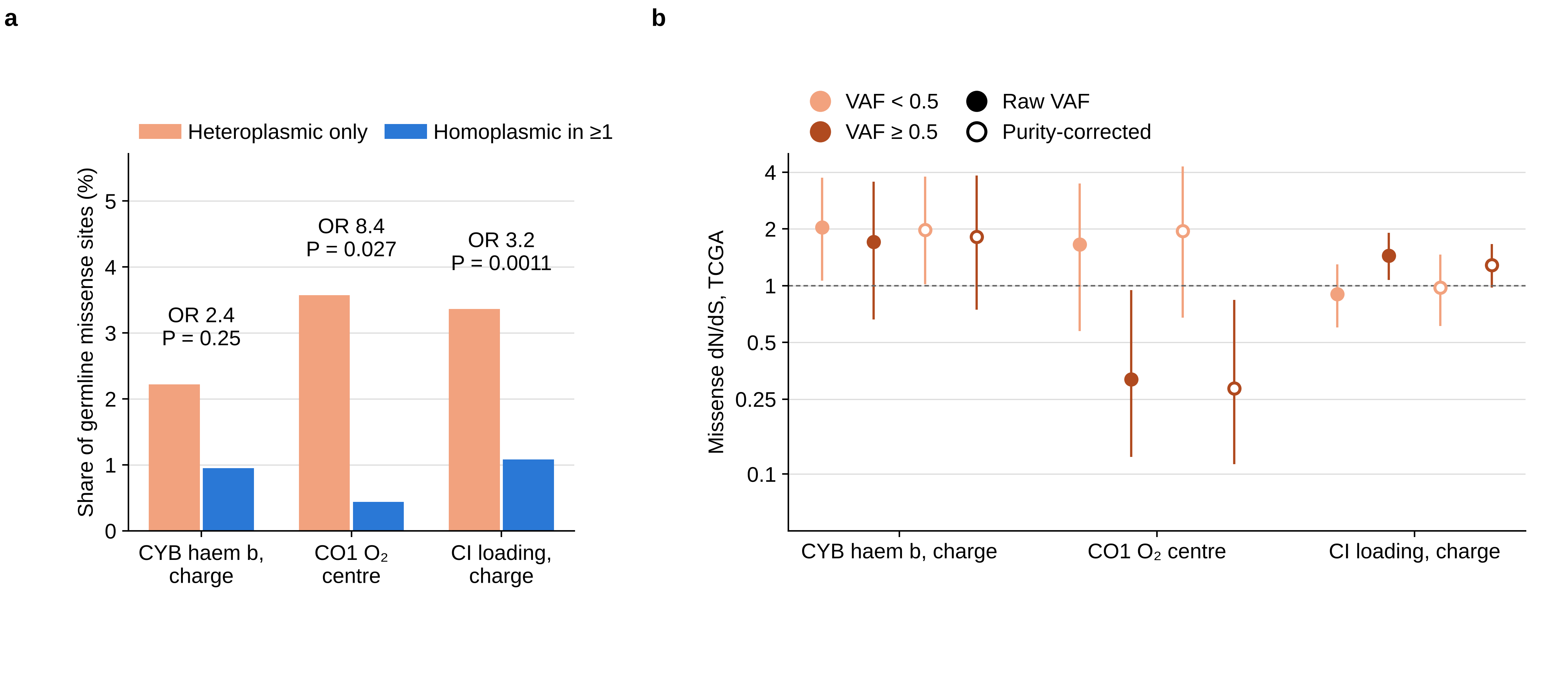

### supple fig6

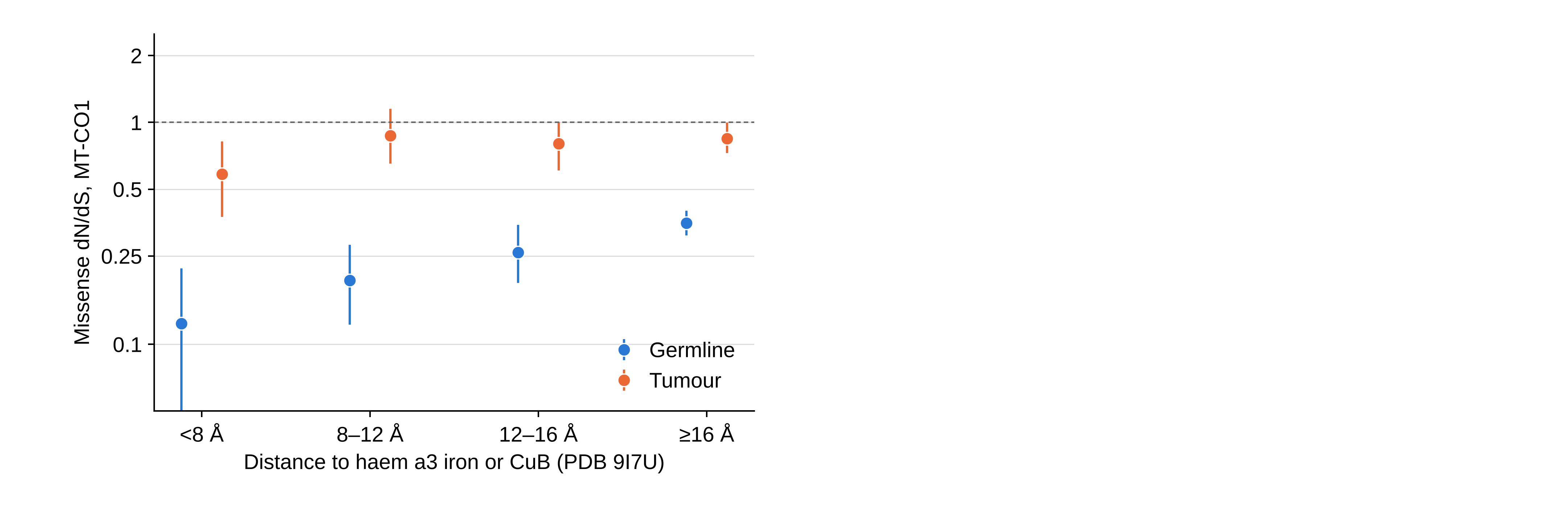

### supple fig7

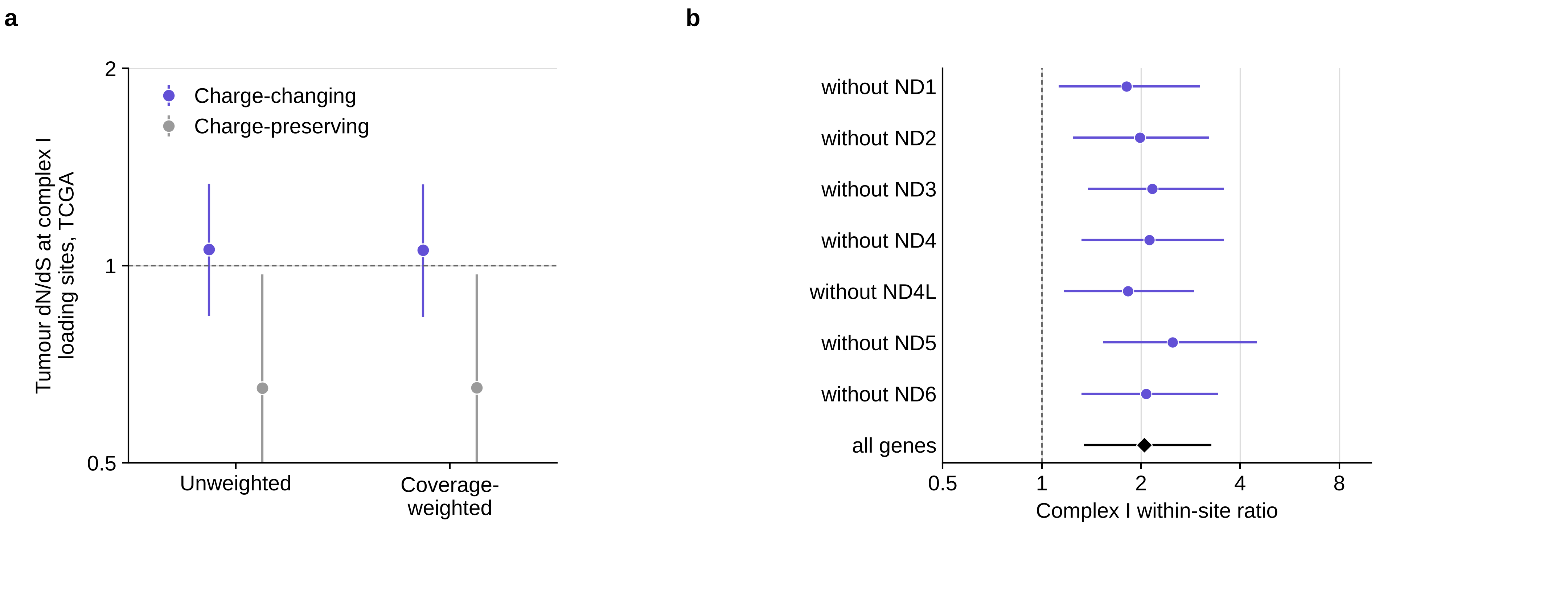

### supple fig8

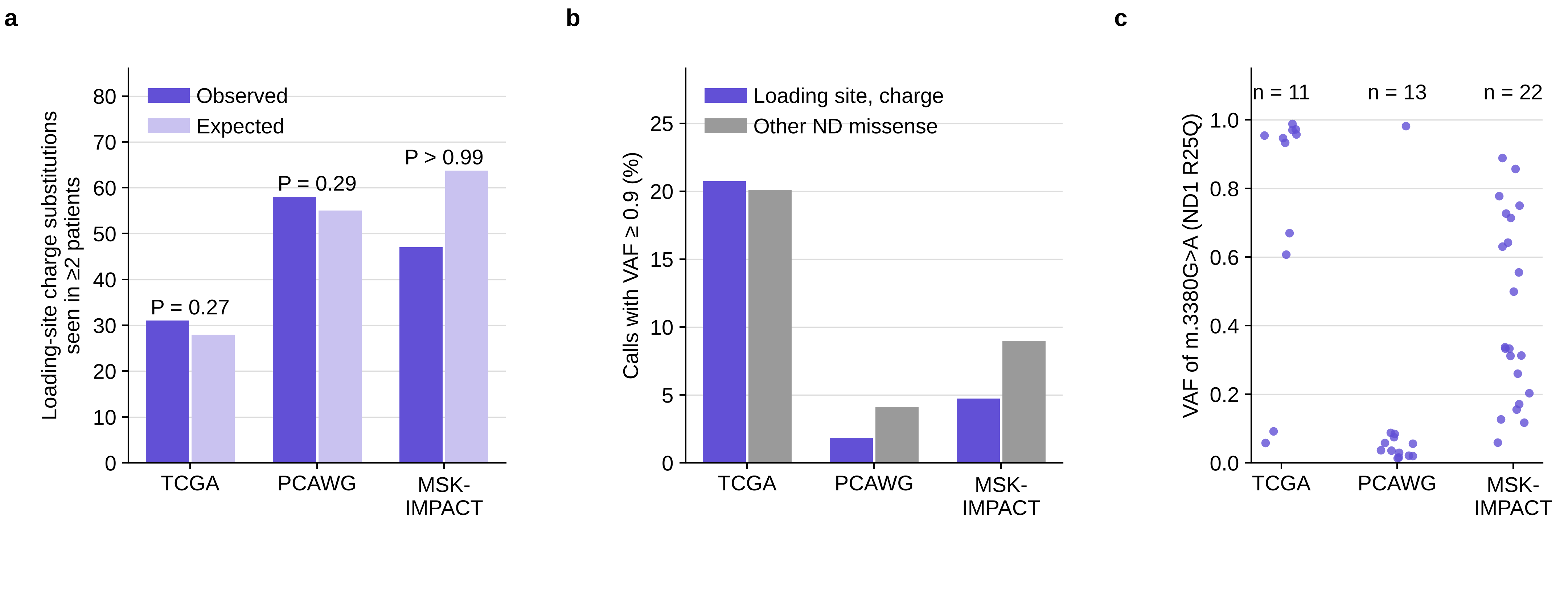

### supple fig9

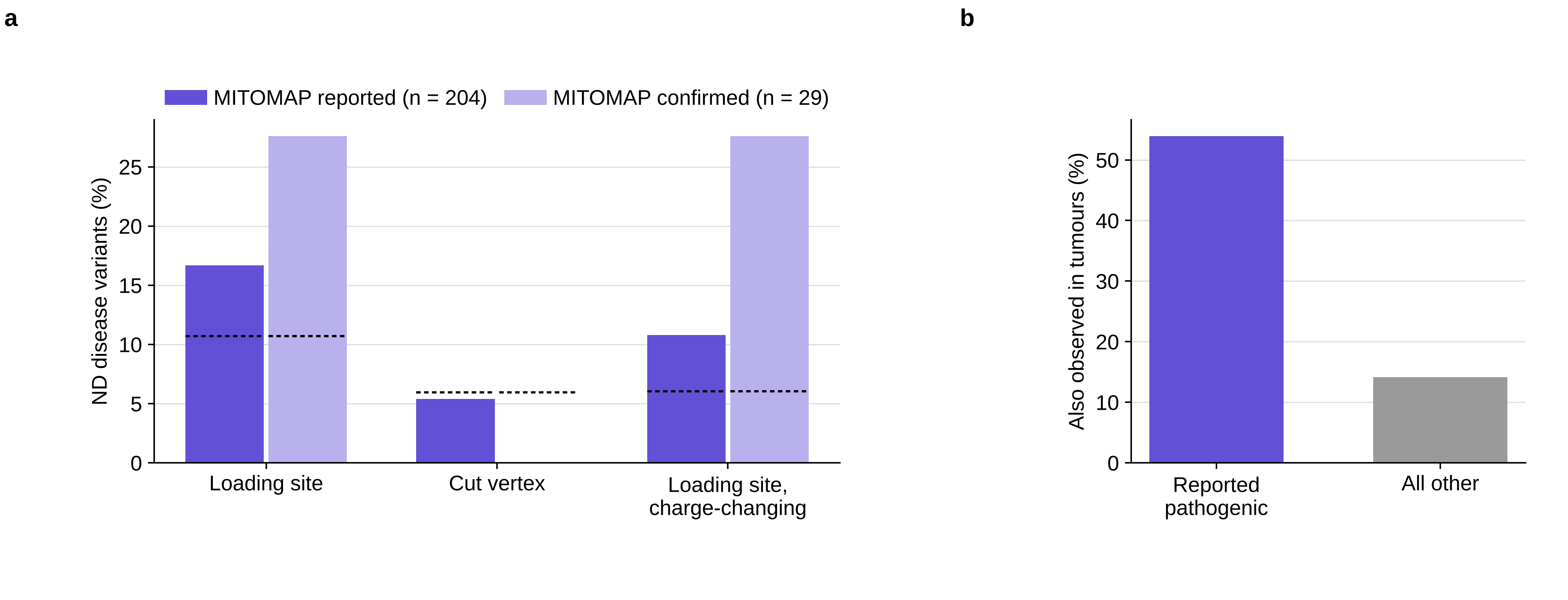

### supple fig10

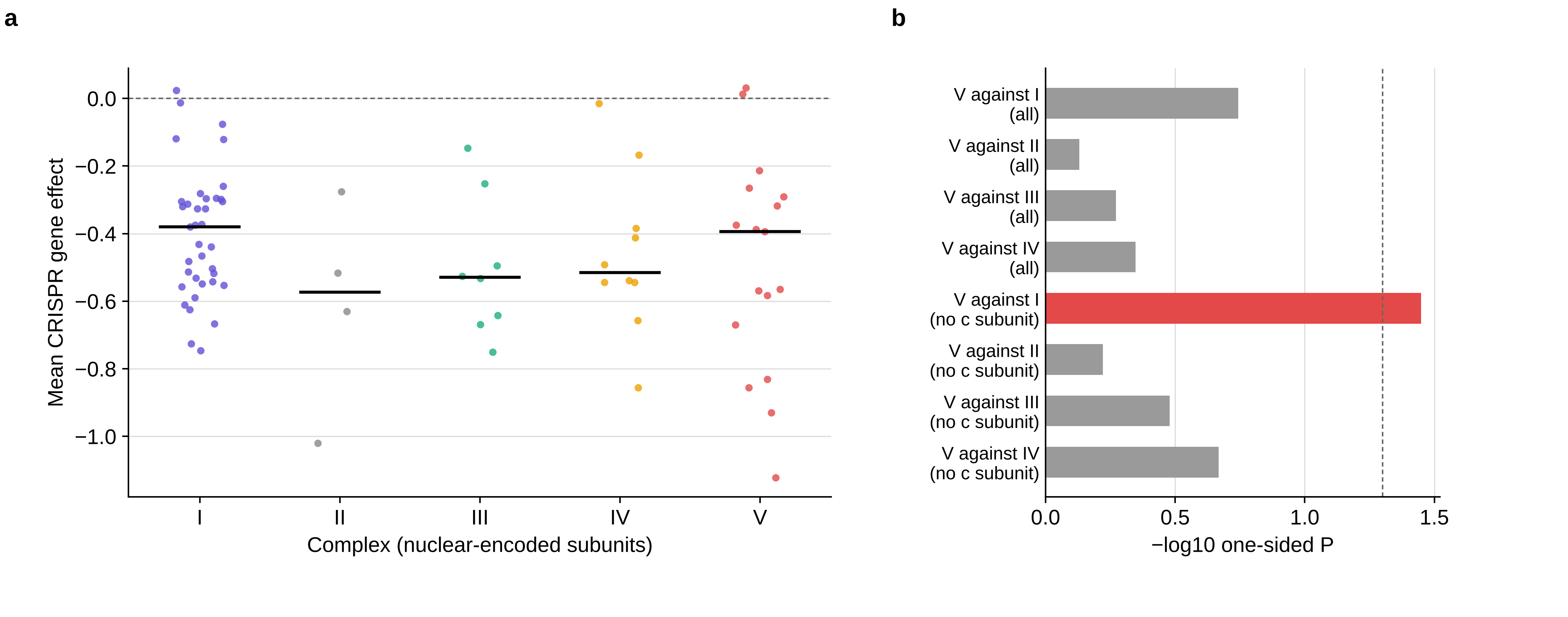

### supple fig11

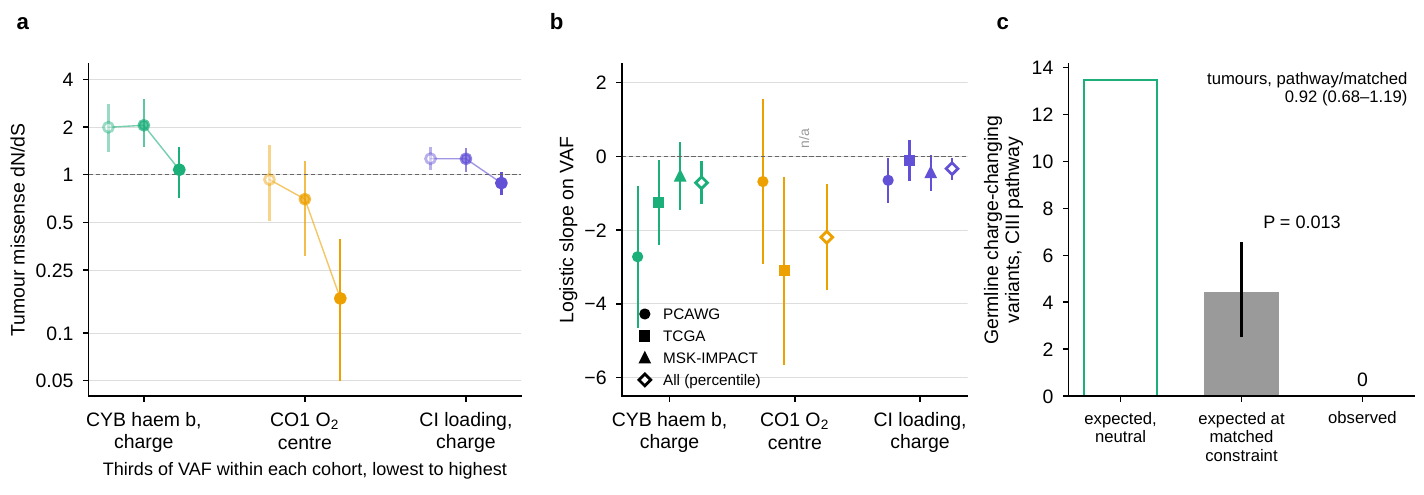
